# Systemic viral vector vaccination induces brain resident memory T cells to drive anti-glioblastoma immunity

**DOI:** 10.64898/2026.06.18.733241

**Authors:** Emily E. Steffke, Laila Latifi, Taijun Hana, Ayaka Hara, Morgan Coombs, Jo Spurgeon, James McAuliffe, Vinnycius Pereira-Almeida, Amanda Wicki, Sara Abdel Malak, Laurine Noblecourt, Caitlin Huguely Wilkinson, John Hancock, Silvia Panetti, Hye Kim, Brita Anderson, Chen Makranz, Nicole Briceno, Meili Zhang, Wei Zhang, Dionne Davis, Hua Song, Michael Bryan, Hideho Okada, Mark Gilbert, Carol Sze Ki Leung, Benoit J. Van den Eynde, Masaki Terabe

**Affiliations:** Laboratory of Integrative Cancer Immunology, Center for Cancer Research, National Cancer Institute, National Institutes of Health, Bethesda, MD, 20892, USA; Neuro-Oncology Branch, National Cancer Institute, Center for Cancer Research, National Cancer Institute, National Institutes of Health, Bethesda, MD, 20892, USA; Ludwig Institute for Cancer Research, Nuffield Department of Medicine, University of Oxford, Oxford, OX3 7DQ, UK; Centre for Immuno-Oncology, Nuffield Department of Medicine, University of Oxford, Oxford, OX3 7DQ, UK; Department of Neurological Surgery, University of California, San Francisco, San Francisco, CA, 94143, USA; Ludwig Institute for Cancer Research, de Duve Institute/UCLouvain, Brussels, Belgium; WELBIO Department, WEL Research Institute, Brussels, Belgium

**Author notes:** Corresponding authors: these authors equally contributed to this work (Carol Sze Ki Leung, Benoit J. Van den Eynde, Masaki Terabe). Masaki Terabe, Ph.D. Laboratory of Integrative Cancer Immunology, Center for Cancer Research, National Cancer Institute, NIH Building 37/Room 1016A, 37 Convent Drive, Bethesda, Maryland 20892.

**Keywords:** Vaccine, glioblastoma, resident memory T cells, tumor immunology, CD8^+^ T-cell

## Abstract

Glioblastoma is a lethal brain tumor that is unresponsive to current cancer immunotherapeutic approaches, including immune checkpoint blockade (ICB). This suggests that initial priming of T cells, rather than their expansion and licensing as effectors, is a restricting feature in this tumor setting. To overcome the limited initiation of CD8^+^ T cell responses, we employed a strong heterologous prime-boost vaccination with the simian adenovirus ChAdOx1 and poxvirus modified vaccinia Ankara (MVA). Vaccination conferred therapeutic efficacy against orthotopic, immune checkpoint-blockade (ICB)-refractory SB28 murine glioblastoma. Vaccination was effective against both the murine tumor antigen, P1A, and a newly identified glioblastoma-associated antigen, Gpr149. Additional treatment with ICB provided no additional benefit. Systemic ChAdOx1/MVA vaccination induced robust infiltration of antigen-specific T cells in tumor-challenged brains, the majority of which exhibited a CD103^+^CD69^+^CD8^+^ tissue-resident memory (TRM)-like phenotype. These cells were polyfunctional, durable in brains with sustained tumor control, and mediated tissue-specific immunological memory. Moreover, intracranial adoptive transfer of glioblastoma-derived antigen-specific TRM-like cells was sufficient to protect naïve recipients from subsequent orthotopic tumor challenge. Together, these findings establish that viral vector vaccination can generate tumor-specific TRM-like cells that mediate effective anti-glioblastoma immunity, providing a rationale for clinical evaluation of ChAdOx1/MVA-based strategies in glioblastoma.

## Introduction

Glioblastoma is an aggressive, immunologically cold tumor with a limited capacity to induce CD8^+^ T cell responses^1^. While immunotherapies enhancing T cell-mediated anti-tumor immunity have transformed outcomes in many cancers, none have proven effective against glioblastoma. Despite evidence for anti-glioblastoma activity of immune checkpoint blockade (ICB) in mouse models using the commonly employed line GL261, poor outcomes for glioblastoma patients have been observed in multiple ICB clinical trials^2–5^. This suggests that initial priming of T cell responses in the brain, rather than their expansion and licensing as effectors, is a restricting feature in the glioblastoma setting. There is therefore strong rationale to use vaccines to promote *de novo* induction of glioblastoma-specific T cell populations.

Heterologous prime-boost vaccination with the simian adenovirus ChAdOx1 followed by modified vaccinia Ankara (MVA) poxvirus induces some of the highest magnitude CD8^+^ T cell responses in humans, and tumor-antigen targeted ChAdOx1/MVA mediates therapeutic efficacy across multiple tumor models, prompting clinical trials for prostate cancer and NSCLC^6–11^. However, whether this strategy can elicit effective CD8^+^ T cell responses in the brain remains unknown. Mounting evidence suggests that control of solid tumors can be achieved with vaccines that elicit tumor-infiltrating CD8^+^ T cells with a tissue-resident memory (TRM)-like phenotype^12–15^. TRMs are typically characterized by expression of CD103 and CD69, which promote tissue retention and restrict egress^12,16^. However, despite extensive testing of peptide-, nucleic acid-, and dendritic cell-based vaccines in glioblastoma, TRM induction has not been observed clinically, and no approach has yet succeeded in phase III trials^17^. Viral vector-based approaches in glioblastoma have largely focused on oncolytic virotherapy, while antigen-encoding viral vector vaccines remain understudied. It is unclear whether ChAdOx1/MVA vaccination induces TRMs and whether vaccine-induced TRMs can mediate anti-tumor immunity in brain tumors.

We therefore employed ChAdOx1/MVA vaccination against orthotopic, syngeneic SB28 mouse glioblastomas. The SB28 model recapitulates key features of human glioblastoma when implanted orthotopically, including high parenchymal invasiveness, low mutational burden, and poor T cell infiltration^2,18^. Unlike the widely-used GL261 model, SB28 is non-responsive to anti-PD-1 and anti-CTLA-4 treatment^2,18^. During our evaluations, we identified a novel endogenous SB28 tumor-associated antigen, Gpr149, as an effective vaccination target. Furthermore, we reveal that ChAdOx1/MVA vaccination via a systemic route induces robust, polyfunctional brain CD103^+^CD69^+^CD8^+^ TRM-like cells that are sufficient to mediate anti-glioblastoma immunity, identifying a clinically deployable strategy to drive CD8^+^ T cells to the brain.

## Results

### ChAdOx1/MVA vaccination demonstrates prophylactic and therapeutic efficacy against P1A-expressing glioblastomas

As endogenous SB28 antigens were unknown, we created a transgenic SB28 line expressing P1A, the murine homologue of MAGE tumor antigens (Supplementary Fig. 1A). P1A mediates anti-tumor immunity in C57BL/6 mice via cytotoxic T lymphocyte responses against P1A_43-51_ presented on H-2D^b^^7^. While constitutive MHC-I expression on SB28 is low, as in typical human glioblastomas, our SB28.P1A clone exhibited no MHC-I expression without IFNγ treatment (Supplementary Fig. 1B). Pre-treating SB28.P1A cells with IFNγ facilitated recognition by *ex vivo* expanded P1A_43-51_-specific CD8^+^ T cells from ChAdOx1/MVA-P1A-vaccinated mice (Fig. 1A, Supplementary Fig. 1C), confirming endogenous processing and presentation of P1A_43-51_ by SB28.P1A, though responses were markedly lower than to P1A_43-51_-peptide-pulsed controls. This modest immunogenicity may reflect that of many human glioblastoma antigens.

**Figure 1.**
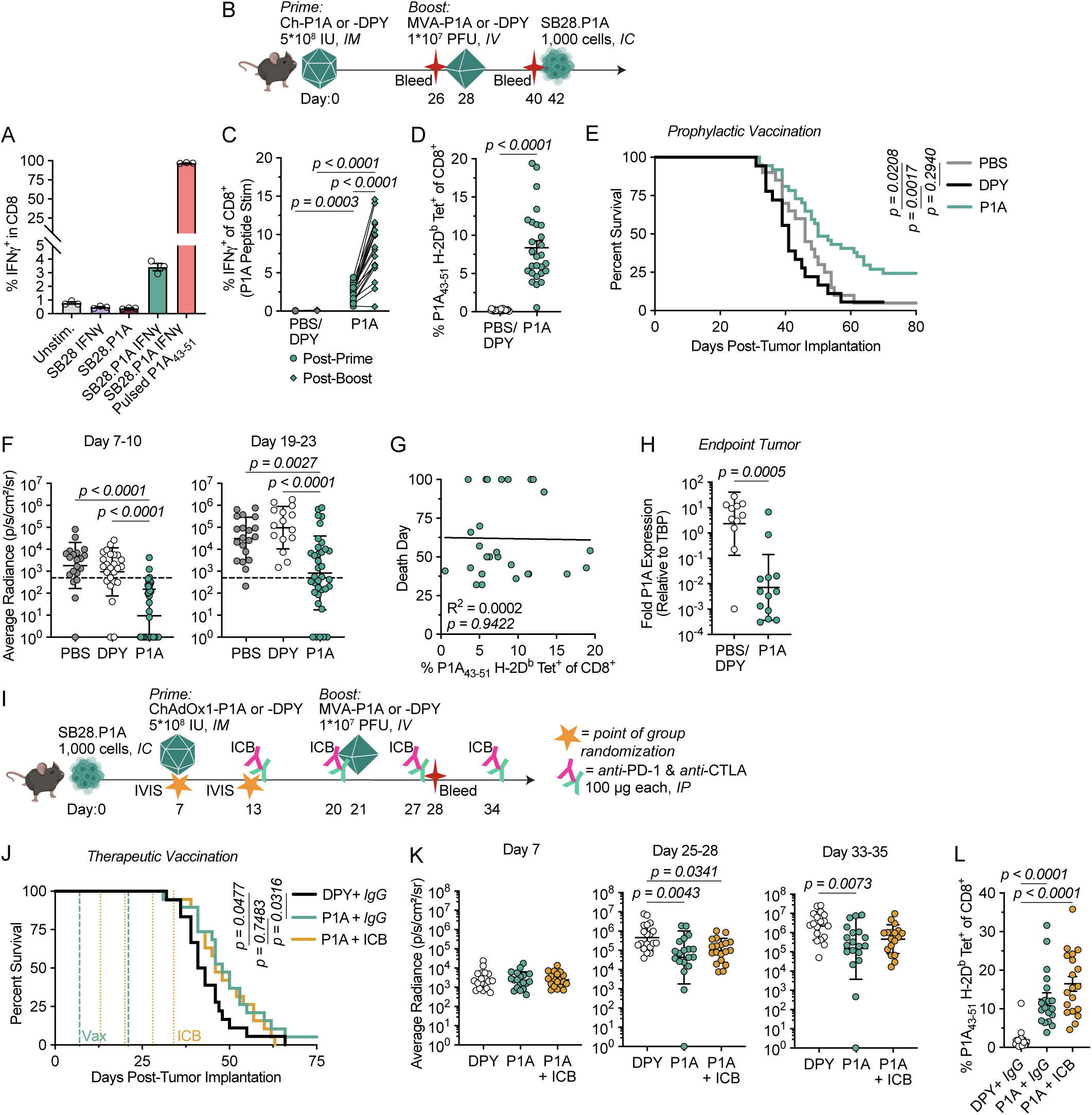
ChAdOx1/MVA vaccination demonstrates prophylactic and therapeutic efficacy against P1A-expressing glioblastomas. (A) Proportion of *ex vivo* expanded P1A-specific CD8^+^ T cells expressing IFNγ after co-culture with wild-type (WT), P1A-transfected, or P1A-peptide pulsed SB28 cells, some of which were pre-treated with IFNγ (*n* = 3 technical replicates). (B) Experimental scheme for (C-H): C57BL/6 mice were primed with ChAdOx1 encoding P1A or DPY, or PBS, then boosted IV 1 month later with MVA encoding P1A or DPY, or PBS. Mice were challenged intracranially with SB28.P1A; blood was sampled 26 days post-prime and 12 days post-boost. (C) Proportion of IFNγ-expressing CD8^+^ T cells in blood post-prime (*circles*) or post-boost (*diamonds*) after *ex vivo* stimulation with P1A peptides (*n* = 18, pooled from 2 experiments). (D) Proportion of CD8^+^ T cells in blood post-boost (D40) binding P1A_43-51_/H-2D^b^-tetramers (*n = 27-28*, pooled from 3 experiments). (E) Survival of SB28.P1A-challenged mice (*n =* 18-37, pooled from 4 experiments). (F) Intracranial IVIS signal on D7-10 (*n =* 20-46, pooled from 5 experiments) and D19-23 (*n* = 15-36, pooled from 4 experiments). (G) Spearman rank correlation for date of death and proportion of P1A_43-51_/H-2D^b^-tetramer^+^CD8^+^ T cells in blood on D40 (*n = 27*, pooled from 3 experiments). (H) Tumor mRNA expression of P1A relative to TATA-box Binding Protein (TBP) at endpoint quantified via RT-qPCR (*n =* 11*-*13, pooled from 2 experiments). (I) Experimental scheme for (J-L): C57BL/6 mice were challenged intracranially with SB28.P1A and screened for tumor presence prior to vaccination with ChAdOx1 vaccines. Mice were screened again on D13 prior to treatment with anti-PD-1 and anti-CTLA-4 (ICB; 100 μg each /dose), non-ICB-treated mice received IgG2a isotype control, ICB treatment was administered weekly for 4 weeks. MVA vaccination occurred 21 days post-tumor implantation. (J) Survival of SB28.P1A-bearing mice (*n* = 19, pooled from 2 experiments). (K) Intracranial IVIS signal (*n =* 19, pooled from 2 experiments). (L) CD8^+^ T cells in blood binding P1A_43-51_/H-2D^b^-tetramers post-boost (*n* = 19, pooled from 2 experiments). Data are presented as mean ± SEM for (A), (D), (L), and geometric mean ± geometric SD for (F), (H), (K). Statistical analysis was performed using RM two-way ANOVA with Fisher’s LSD multiple comparisons test for (C), two-tailed t-test with Welch’s correction for (D), log-rank test for (E), simple linear regression for (G), Kruskal-Wallis test with Dunn’s multiple comparisons for (F), (K), (L), two-tailed Mann-Whitney test for (H), and Gehan-Breslow-Wilcoxon test for (J).

To assess prophylactic efficacy, mice were vaccinated with ChAdOx1/MVA vectors encoding P1A or DPY (an irrelevant antigen) in a prime-boost regimen before intracranial SB28.P1A challenge (Fig. 1B). ChAdOx1-P1A primed robust circulating P1A-specific CD8^+^ T cell responses that were amplified by MVA-P1A (Fig. 1C-D). P1A, but not DPY, vaccination reduced tumor burden and prolonged survival, with 25% achieving long-term survival (Fig. 1E-F). IFNγ pre-treatment of SB28.P1A prior to implantation did not significantly impact survival (Supplementary Fig. 1D). Circulating P1A-specific CD8^+^ T cell magnitude did not correlate with survival (Fig. 1G). However, endpoint tumors from P1A-vaccinated mice exhibited reduced P1A expression, consistent with antigen-specific immune pressure and immunoediting in non-cleared tumors (Fig. 1H).

Given this prophylactic efficacy, we next evaluated therapeutic ChAdOx1/MVA vaccination. Because a 28-day prime-boost interval was incompatible with rapid SB28.P1A growth, we optimized alternative regimens. Thirteen strategies varying dose, route, and timing were tested in tumor-free mice to identify those rapidly inducing functional antigen-specific CD8^+^ T cells (Supplementary Fig. 2A). Priming with 1×10^8^ IU ChAdOx1-P1A intravenously (IV) or 5×10^8^ IU intramuscularly (IM), followed two weeks later by IV boosting with 1×10^7^ PFU of MVA-P1A (Strategies 11 and 12) induced the highest frequencies of circulating P1A-specific CD8^+^ T cells by day 20 post-prime (Supplementary Fig. 2B-C). Compared to IV/IV, IM/IV vaccination elicited greater T cell polyfunctionality both post-prime and post-boost (Supplementary Fig. 2D-E).

The IM/IV two-week regimen was therefore applied therapeutically, with priming following confirmation of tumor establishment. Though anti-PD-1 and anti-CTLA-4 (ICB) treatment is ineffective against intracranial SB28^2^, we investigated whether ICB enhances vaccine efficacy (Fig. 1I). P1A vaccination significantly reduced tumor burden and prolonged survival relative to DPY controls (Fig. 1J-K, Supplementary Fig. 1F).

Adjuvant ICB conferred no additional benefit, despite modestly increasing circulating P1A_43-51_/H-2D^b^-tetramer^+^CD8^+^ T cells (P1A_43-51_-specific T cells) (Fig. 1L, Supplementary Fig. 1E). As in the prophylactic setting, circulating P1A_43-51_-specific T cell magnitudes did not correlate with survival (Supplementary Fig. 1F, G). While prior studies have demonstrated vaccine-driven control of immunologically “warmer” glioblastoma models expressing xenogeneic antigens, these data demonstrate that ChAdOx1/MVA vaccines targeting a murine antigen can control immunologically cold SB28 tumors, underscoring the translational relevance of this approach.

### ChAdOx1/MVA vaccination is prophylactically and therapeutically effective against a novel SB28 TAA, Gpr149

To enhance the clinical relevance of our vaccination model beyond a transfected antigen, we sought to identify endogenous SB28 vaccine targets. We employed whole-exome and mRNA sequencing of SB28 cells and mouse brains and astrocytes to identify neoantigens and tumor-associated antigens (TAAs) (Supplementary Fig. 3A). SB28 had fewer neoantigen candidates than the GL261 glioblastoma model, though it harbored similar numbers of TAA candidates (Supplementary Fig. 3B-C). No SB28 neoantigens with both high expression and strongly predicted MHC-I binding epitopes were identified (Supplementary Fig. 3D-F, Supplemental Data Table 1A-B). Sc5d was the only expressed neoantigen with a “weakly” predicted MHC-I epitope (Supplemental Data Table 1B). However, we identified 16 TAAs with high relative SB28 expression by mRNA-Seq and minimal expression in adult mice based on NCBI Gene, five of which – *Gpr149*, *Cdh6*, *Fbn2, Shox2,* and *Ifg2bp1 –* showed high SB28 and negligible tissue expression by RT-qPCR (Fig. 2A, Supplementary Fig. 3G, Supplementary Fig. 4A-B, Supplemental Data Table 1C). *Cdh6*, *Fbn2,* and *Shox2* were also overexpressed in human TCGA glioblastoma samples (Supplemental Data Table 1D). All five TAAs contained multiple predicted MHC-I binding epitopes throughout their coding sequences (Supplementary Fig. 4C); accordingly, we encoded complete TAA sequences in our vaccine constructs where possible.

**Figure 2.**
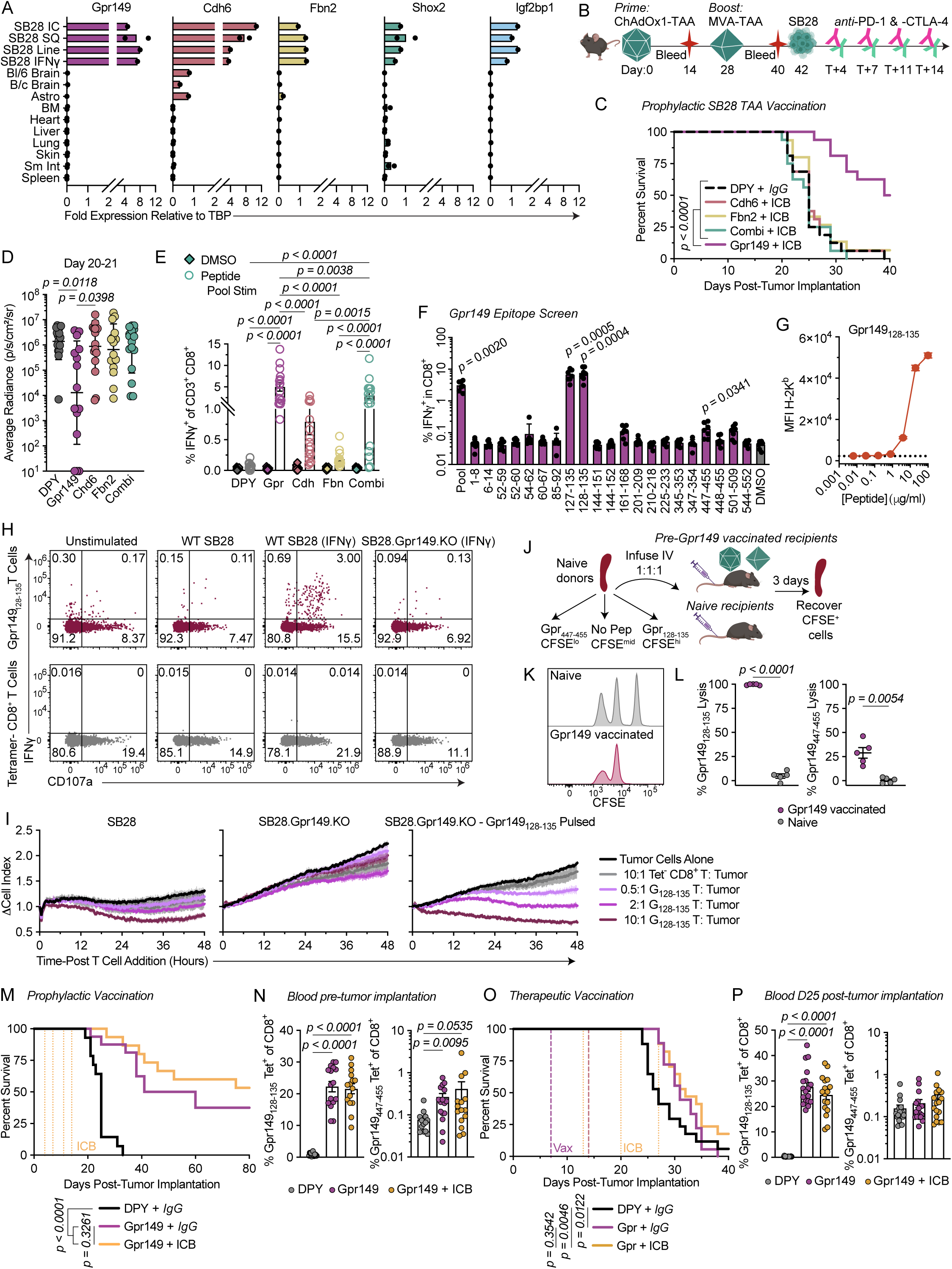
ChAdOx1/MVA vaccination is prophylactically and therapeutically effective against a novel SB28 TAA, Gpr149. (A) Expression of TAA candidate genes relative to TBP as measured by RT-qPCR. IC, intracranial; SQ, subcutaneous; IFNγ, IFNγ-pretreated; Bl/6, C57BL/6; B/c, BALB/c; Astro, astrocytes; BM, bone marrow; Sm Int, small intestine (*n =* 1-2). (B) Experimental scheme for (C-E): C57BL/6 mice were vaccinated with ChAdOx1/MVA vaccines encoding SB28 TAAs or DPY one month apart, then challenged intracranially with 500 SB28 tumor cells; all mice receiving SB28 TAA vaccines also received anti-PD-1 and anti-CTLA-4 (ICB; 100 μg each /dose) on days 4, 7, 11, and 14 post-tumor implantation. Blood was sampled 14 days post-prime and 12 days post-boost. (C) Survival of SB28-challenged mice (*n* = 15-16, pooled from 2 experiments). (D) Intracranial IVIS signal (*n =* 15-16, pooled from 2 experiments). (E) Proportion of CD8^+^ T cells expressing IFNγ after *ex vivo* restimulation of PBMCs with the relevant pool of minimal epitope peptides (*circles*) or without stimulation (DMSO, *diamonds*) 2 weeks post-boost (*n* = 16, pooled from 2 experiments). (F) Frequency of CD8^+^ T cells expressing IFNγ after *ex vivo* restimulation of splenocytes from Gpr149-vaccinated mice with minimal epitope peptides (*n* = 7, pooled from 2 experiments). (G) RMA-S H-2K^b^ median fluorescence intensity (MFI) after culture with peptide at increasing concentrations; dotted line indicates baseline (*n* = 3 technical replicates). (H) Reactivity of FACS-isolated Gpr149_128-135_-specific or nonspecific T cells against SB28 cells; IFNγ indicates tumor cells pre-treated with IFNγ prior to co-culture. (I) Fold change over time in SB28 cell growth during co-culture with sorted T cells. Thick lines represent mean change in cell index, thin vertical lines represent SEM (*n* = 3 technical replicates). (J) Experimental scheme for (K-L): Syngeneic splenocytes from donor mice were pulsed with peptides and labeled with different concentrations of CFSE prior to IV infusion into naïve or vaccinated recipients. (K) Splenocyte CFSE signals 3 days post-cell infusion. (L) Proportional *in vivo* lysis of peptide-pulsed splenocytes (*n* = 5). (M) Survival of SB28-challenged mice (*n* = 14-16, pooled from 2 experiments). Mice were prophylactically vaccinated as in (B), except some Gpr149-vaccinated mice did not receive ICB. (N) Tetramer-binding CD8^+^ T cells in blood post-boost (D40) for mice in (M) (*n =* 14-16, pooled from 2 experiments). (O) Survival of SB28-bearing mice therapeutically treated with ChAdOx1-Gpr149 or - DPY on D7 and MVA-Gpr149 or -DPY on D14. Some mice also received anti-PD-1 + anti-CTLA-4 (ICB) on D13, 20, 27, and 34 (*n* = 17-18, pooled from 2 experiments). (P) Tetramer-binding CD8^+^ T cells in blood post-boost (D25) for mice in (O) (*n* = 17-18, pooled from 2 experiments). Data are presented as mean for (A), mean ± SEM for (E), (L), (N), (P), geometric mean ± geometric SD for (D), and mean ± SD for (F-G). Statistical analysis was performed using log-rank test for (C), (M), Kruskal-Wallis test with Dunn’s multiple comparisons for (D), (F), (N), (P), two-way ANOVA with Tukey’s multiple comparisons test for (E), unpaired t-test with Welch’s correction for (L), and Gehan-Breslow-Wilcoxon test for (O).

We constructed four novel sets of ChAdOx1 and MVA vectors: Gpr149 and Cdh6 encoded independently and entirely; Fbn2 encoded by 11 minigenes aligned to top predicted immunogenic regions; and Shox2, Igf2bp1, and the Sc5d neoepitope encoded in tandem “Combi” vectors (Supplementary Fig. 4D, Supplemental Data Table 1E).

CD8^+^ T cells expanded *ex vivo* from splenocytes of TAA-immunized mice produced IFNγ when co-cultured with corresponding transgenic DC2.4 clones expressing identical TAA inserts to those in all vaccines except Fbn2, indicating endogenous processing and presentation of immunogenic Gpr149, Cdh6, Shox2, and Igf2bp1 epitopes on H-2^b^ molecules (Supplementary Fig. 4E-F).

To determine the efficacy of TAA-targeted vaccination against SB28, the vaccines were evaluated prophylactically with adjuvant ICB (Fig. 2B). Though all TAA vaccines were immunogenic, only Gpr149 vaccination reduced tumor burden and prolonged survival, with half of mice surviving long-term (Fig. 2C-E). Gpr149 vaccination elicited the highest frequency of antigen-specific CD8^+^ T cells in blood post-prime and post-boost, measured by IFNγ secretion following restimulation with peptide pools containing top predicted epitopes for each TAA (Fig. 2E, Supplementary Fig. 5A, Supplemental Data Table 2), whereas Combi vaccines induced more polyfunctional responses (Supplementary Fig. 5B). No CD4^+^ T cell responses were detected (Supplementary Fig. 5C). Overall, prophylactic Gpr149-targeted vaccination was sufficient to protect against orthotopic SB28 tumors.

Gpr149_128-135_ (VCYNFYTM) was the dominant minimal epitope, with secondary responses to Gpr149_447-455_ (RSVFNTITV) and minor responses to additional epitopes detected by TNF secretion (Fig. 2F, Supplementary Fig. 5D). Immunogenic epitopes were also identified within the other TAAs, though the Sc5d neoepitope was not immunogenic (Supplementary Fig. 5D). Using TAP-deficient RMA-S cells, we verified the MHC-I restriction and approximated the binding affinities of each epitope, finding Gpr149_128-135_ bound H-2K^b^ and Gpr149_447-455_ bound H-2D^b^ (Fig. 2G, Supplementary Fig. 5E).

Tetramers were generated for the strongest TAA epitopes (Supplementary Fig. 5F). Gpr149_128-135_/H-2K^b^-tetramer^+^CD8^+^ (Gpr149_128-135_-specific) T cells but not tetramer^-^CD8^+^ T cells sorted from vaccinated mice expressed IFNγ and CD107a when exposed to IFNγ-pre-treated WT SB28, but not to untreated or Gpr149-knockout (KO) SB28 (Fig. 2H, Supplementary Fig. 5G). In a real-time growth monitoring assay, Gpr149_128-135-_specific but not tetramer^-^CD8^+^ T cells lysed WT but not Gpr149.KO SB28 in a dose-dependent manner (Fig. 2I); Gpr149_128-135_ peptide-pulsing of Gpr149.KO cells restored killing. An *in vivo* lysis assay in Gpr149-vaccinated mice demonstrated ∼100% and ∼25% specific lysis of Gpr149_128-135_- and Gpr149_447-455_-pulsed targets, respectively (Fig. 2J-L). Thus, ChAdOx1/MVA-Gpr149 vaccination protects against SB28 tumors primarily through induction of Gpr149_128-135_-specific CD8^+^ T cells mediating antigen-specific cytolysis.

Prophylactic ChAdOx1/MVA-Gpr149 vaccination prolonged survival of SB28-challenged mice without adjuvant ICB (Fig. 2M). Prior to tumor implantation, >20% and ∼0.2% of CD8^+^ T cells in the blood were Gpr149_128-135_ and Gpr149_447-455_-specific, respectively (Fig. 2N).

Despite using a compressed IM/IV prime-boost schedule to treat rapidly growing WT SB28 tumors, therapeutic Gpr149 vaccination significantly prolonged survival and induced high frequencies of circulating Gpr149_128-135_-specific T cells (Fig. 2O-P).

Combinatorial ICB conferred only a marginal, non-significant benefit. Together, these data show that ChAdOx1/MVA vaccines targeting either P1A or the endogenous TAA Gpr149 significantly improve survival in mice with established, ICB-refractory orthotopic SB28 tumors, highlighting the translational potential of this approach for glioblastoma.

### Vaccination remodels tumor immune infiltrate and induces high levels of CD103^+^CD69^+^ TRM-like CD8^+^ T cells

While antigen-specific CD8^+^ T cells were observed in the blood of immunized mice, their infiltration of the brain was unclear. We therefore collected tumor-challenged and contralateral brain hemispheres from prophylactically P1A- or DPY-vaccinated and/or SB28.P1A-challenged mice for flow cytometric analysis (Fig. 3A). We identified 32 distinct immune subsets that varied in relative abundance by experimental group and tumor clearance status (Fig. 3B-D, Supplementary Fig. 6A-H). While CD8^+^ T cells were sparse in PBS-treated tumor-bearing mice, P1A and DPY vaccination expanded the CD8^+^ T cell compartment (Fig. 3B-D and Supplementary Fig. 6A-F, H). Meanwhile, tumor-associated macrophages (TAMs), eosinophils, mMDSCs, and T_reg_ cells were increased in the tumor-bearing hemispheres of DPY-vaccinated mice relative to contralateral hemispheres (Supplementary Fig. 6F-G), and in P1A-vaccinated tumor-challenged hemispheres compared to non-challenged brains (Supplementary Fig. 6E, G). However, these immunosuppressive populations did not differ between tumor-challenged and contralateral hemispheres of P1A-vaccinated mice (Supplementary Fig. 6C, G), with tumor-cleared hemispheres showing reduced TAM and mMDSC infiltration and resembling brains of non-challenged, P1A-vaccinated mice (Supplementary Fig. 6A).

**Figure 3.**
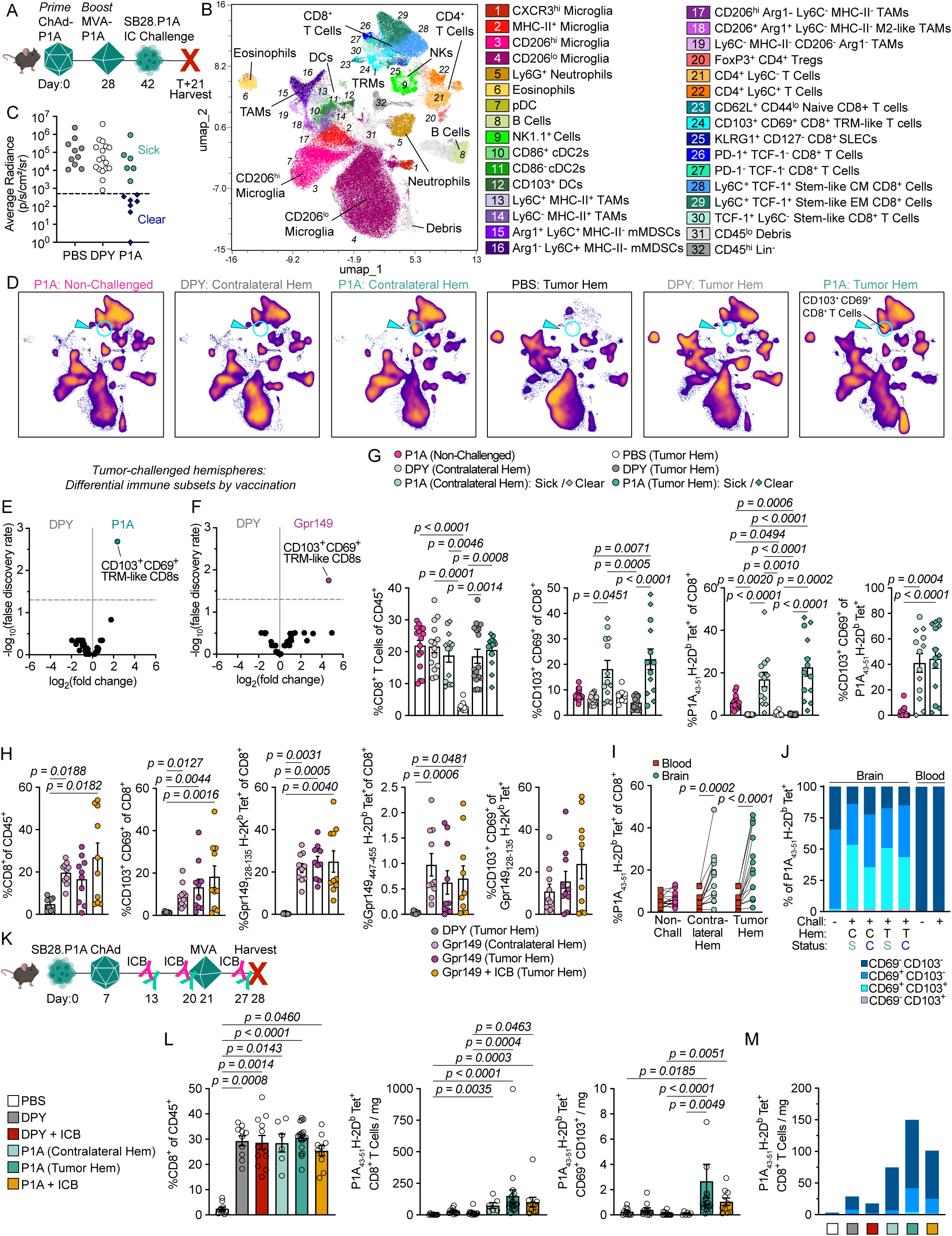
Vaccination remodels tumor immune infiltrate and induces high levels of CD103^+^CD69^+^ TRM-like CD8^+^ T cells. (A) Prophylactic experimental scheme for (B-F), (H), and (J-K). Mice were vaccinated with ChAdOx1/MVA-P1A or -DPY, or were non-vaccinated (PBS), then challenged with SB28.P1A tumors (except non-challenged group). Brains and blood were collected 21 days post-tumor challenge for flow cytometry. (B) UMAP of flow cytometry data for CD45^+^ cells from 50 brain samples. (C) Intracranial IVIS signal on D20 prior to brain collection (*n =* 10-17, pooled from 3 experiments). (D) Heatmap distribution of cells in (B) by treatment and brain hemisphere (Hem) (*n* = 5-10). (E) Comparative expression of subsets in (B) for tumor-bearing hemispheres (*n =* 5-10). (F) Mice were treated prophylactically with ChAdOx1/MVA-Gpr149 or -DPY as in Fig. 2B, and brains were collected 21 days post-tumor implantation. Comparative expression of subsets listed in Supplementary Fig. 7A for tumor-bearing hemispheres (*n =* 3-5). (G) Flow cytometry analysis of immune cells in brains. Data from “tumor-cleared” mice (see C) are designated by diamonds (*n* = 10-17, pooled from 3 experiments). (H) Flow cytometry analysis of brain immune cells (*n =* 8-10, pooled from 2 experiments). (I) P1A_43-51_/H-2D^b^-tetramer-binding of brain or blood CD8^+^ T cells (*n* = 13-16, pooled from 3 experiments). (J) CD103/CD69 expression on P1A_43-51_/H-2D^b^-tetramer^+^CD8^+^ T cells (*n* = 6-16, pooled from 3 experiments). (K) Therapeutic experimental scheme for (L-M): brains were collected 28 days post-tumor challenge for flow cytometry. (L) Flow cytometry analysis of brain immune cells (*n* = 6-19, pooled from 2-3 experiments except for “P1A (Contralateral Hem)” group). (M) Numbers of brain P1A_43-51_/H-2D^b^-tetramer^+^ T cells by CD103/CD69 expression (*n* = 6-19, pooled from 2-3 experiments except for “P1A (Contralateral Hem)” group). Data are presented as mean for (J), (M), and mean ± SEM for (G-H), (L). Statistical analysis was performed using edgeR for (E-F), Kruskal-Wallis test with Dunn’s multiple comparisons for (G-H), (L), and RM two-way ANOVA with Sídǎk’s multiple comparisons test (Blood vs. Brain) for (I).

Of the 32 immune subsets, only CD8^+^ T cells expressing the canonical CD103 and CD69 TRM markers differed significantly following antigen-specific vaccination (Fig. 3E). These cells were singularly increased in tumor-challenged brains by both P1A and Gpr149 vaccines compared to DPY vaccines (Fig. 3E-H, Supplementary Fig. 7A-D), indicating that TRM-like cells may contribute to differential survival outcomes.

P1A_43-51_-specific T cells were enriched in both hemispheres of tumor-challenged, vaccinated mice relative to non-challenged brains, consistent with tumor-specific recruitment or expansion (Fig. 3G, Supplementary Fig. 6I). Gpr149 vaccination similarly induced T cells recognizing both Gpr149 epitopes in SB28 tumors, with no effect of adjuvant ICB (Fig. 3H, Supplementary Fig. 7D). Antigen-specific CD8^+^ T cells were significantly more frequent in tumor-challenged brains than in blood at the same timepoint in both P1A- and Gpr149-vaccinated mice (Fig. 3I, Supplementary Fig. 7E), indicating that systemic ChAdOx1/MVA vaccination promotes central nervous system (CNS) trafficking.

Over 40% of P1A_43-51_-specific T cells in tumor-challenged brains exhibited a TRM-like CD103^+^CD69^+^ double-positive (DP) phenotype (Fig. 3G, J). CD69, but not CD103, was expressed on P1A_43-51_-specific T cells from non-challenged mice, suggesting that tumor exposure drives CD103 acquisition (Fig. 3J). CD103 and CD69 expression were comparable between tumor-challenged and contralateral hemispheres and between tumor-bearing (sick) and tumor-cleared P1A-vaccinated mice, indicating that CD103 expression was not constrained to cells in direct tumor contact (Fig. 3G, J). This CD103/CD69 upregulation was specific to CD8^+^ T cells, as CD4^+^ T cell frequencies and phenotypes were similar across groups (Supplementary Fig. 6J-K, Supplementary Fig. 7F). In contrast to brain-resident cells, P1A_43-51_-specific T cells in blood lacked CD69 and CD103 expression (Fig. 3J). While a smaller fraction of Gpr149_128-135_-specific T cells exhibited a DP phenotype (16%) compared to P1A (45%), most expressed CD69 and additional TRM markers, with 58% CD39^+^CD49a^+^Runx3^+^ (Fig. 3H, Supplementary Fig. 7G-H). Similar TRM marker expression was observed on Gpr149_447-455_-specific CD8^+^ T cells (Supplementary Fig. 7I). TRM phenotypes were unaffected by ICB (Supplementary Fig. 7D, J). Collectively, prophylactic tumor-targeted systemic ChAdOx1/MVA vaccination significantly increased tumor-infiltrating, antigen-specific TRM-like CD8^+^ T cells in orthotopic glioblastoma.

Therapeutic vaccination also remodeled the tumor immune compartment relative to PBS-treated controls. Both DPY and P1A vaccines increased CD8^+^ T cell infiltration and reduced M2-like TAMs, mMDSCs, and T_reg_ cells (Fig. 3K, Supplementary Fig. 6L-N).

P1A vaccination induced greater numbers of total and TRM-like P1A_43-51_-specific T cells than DPY vaccination, though most P1A_43-51_-specific T cells were CD69^+/-^CD103^-^ at the timepoint investigated (Fig. 3L-M). Given that cellular memory develops over weeks^19^, fewer TRM-like cells were expected in the therapeutic setting one week post-boost relative to the prophylactic setting five weeks post-boost. Adjuvant ICB had minimal impact on CD8^+^ T cell numbers or phenotypes, consistent with its lack of survival benefit (Fig. 1J, 2O). Together, these data indicate that ChAdOx1/MVA vaccination expands tumor-specific CD8^+^ T cells and reduces immunosuppressive myeloid populations, slowing SB28 tumor progression and prolonging survival.

### Vaccine-induced CD8^+^ T cells in brain tumors are phenotypically distinct from blood CD8^+^ T cells and are not terminally exhausted

Given the lack of correlation between circulating tumor-specific CD8^+^ T cell magnitude and survival, we investigated phenotypic differences between blood and brain-tumor infiltrating CD8^+^ T cells to better understand their capacities for exerting anti-tumor immunity. Unsupervised clustering of CD8^+^ T cells from P1A, DPY, or non-prophylactically vaccinated (PBS-treated) mice revealed 21 subsets, most composed predominantly of either brain- or blood-derived cells but not both (Fig. 4A-C). A similar analysis in Gpr149-vaccinated, SB28-challenged mice showed comparable segregation of blood and brain phenotypes (Supplementary Fig. 8A-E), indicating that circulating CD8^+^ T cell phenotypes do not predict those infiltrating the brain.

**Figure 4.**
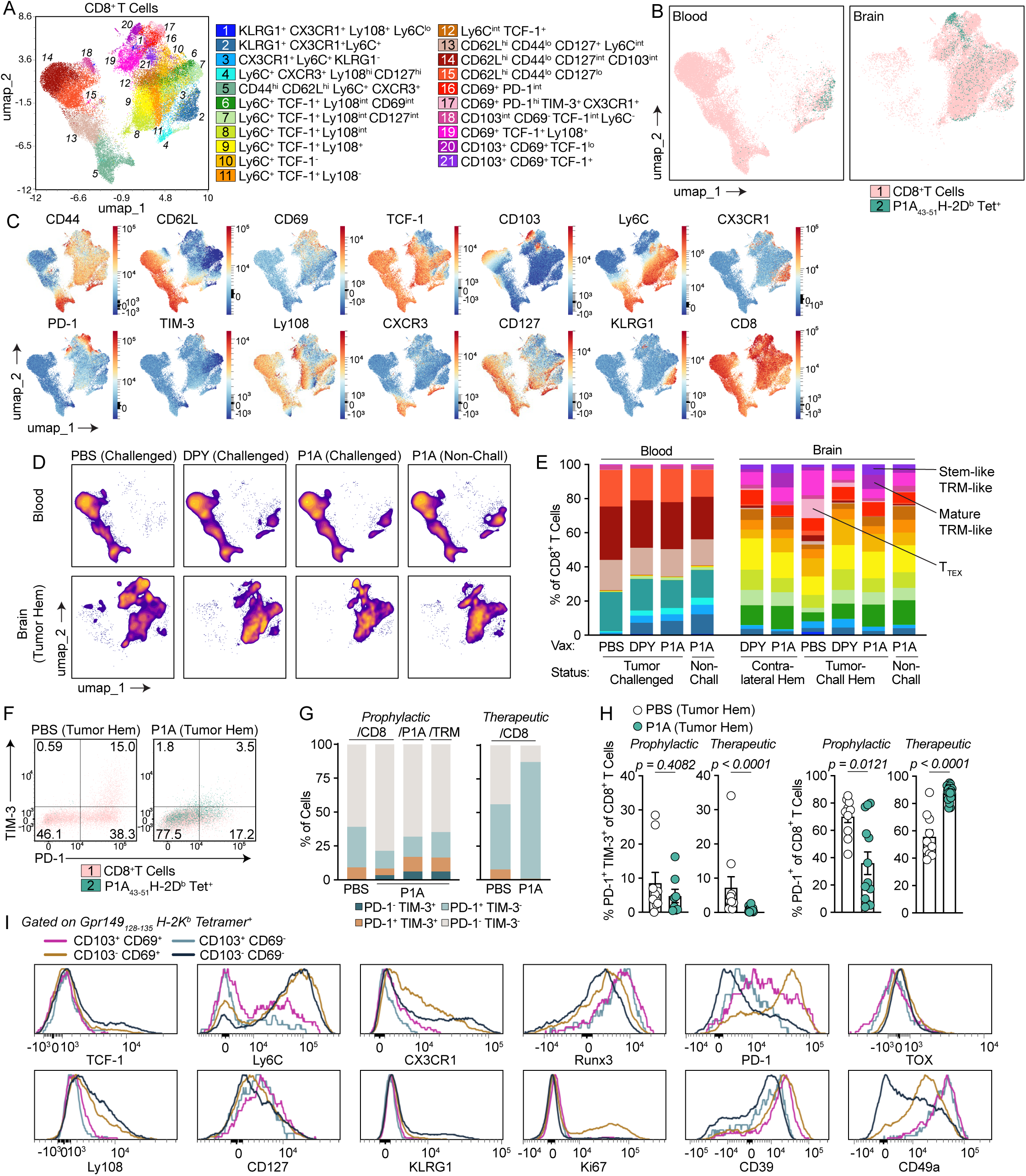
Vaccine-induced CD8^+^ T cells in brain tumors are phenotypically distinct from blood CD8^+^ T cells and are not terminally exhausted. Panels (A-H) refer to flow cytometry data from brains and blood of P1A-prophylactically vaccinated mice as described in Figure 3A. Panels (G-H) additionally refer to flow cytometry data from brains of therapeutically P1A-vaccinated mice as described in Supplementary Fig. 6. Panel (I) refers to flow cytometry data from brains of Gpr149-prophylactically vaccinated mice as described in Supplementary Fig. 7. (A) UMAP of CD8^+^ T cells from 50 brain and 35 blood samples. (B) Overlay of P1A_43-51_/H-2D^b^-tetramer^+^ cells atop UMAP in (A) for blood or brains (*n =* 35-50). (C) Heatmap of phenotypic markers for UMAP in (A) (n = 85). (D) Heatmap distribution of cells in (A) by treatment and tissue (*n* = 5-10). (E) Proportion of CD8^+^ cells in brains or blood corresponding to subsets in (A), stacked ascending from bottom (*n =* 5-10). (F) TIM-3/PD-1 expression on brain CD8^+^ and P1A_43-51_/H-2D^b^-tetramer^+^ T cells from PBS-treated or P1A-vaccinated mice (concatenated from *n =* 5-10). (G-H) Proportion of TIM-3/PD-1-expressing cells among brain CD8^+^, P1A_43-51_/H-2D^b^-tetramer^+^ (P1A), or CD103^+^CD69^+^ (TRM) T cells from PBS-treated or P1A-vaccinated mice (*n* = 10-19, each pooled from 2-3 experiments). (I) Expression of phenotypic markers on brain Gpr149_128-135_/H-2K^b^-tetramer^+^CD8^+^ T cells by CD103/CD69 expression (concatenated from *n =* 20 mice). Data are presented as mean for (E), (G), and mean ± SEM for (H). Statistical analysis was performed using two-tailed Mann-Whitney test for (H).

Most P1A_43-51_-specific T cells in the blood were short-lived effector cells (SLECs) in Cluster 2 (KLRG1^+^CX3CR1^+^Ly6C^+^), the only cluster with notable brain-blood overlap. In contrast, most P1A_43-51_-specific T cells in brains were TRM-like cells in Cluster 20 (CD103^+^CD69^+^TCF-1^lo^) (Fig. 4B). Overall, cluster composition varied more by tissue than by treatment, though treatment-specific signatures were observed (Fig. 4D-E). P1A but not DPY vaccination or PBS upregulated TRM-like clusters (20 and 21) in tumor-challenged brains, whereas PBS-treated tumors harbored proportionally more exhausted-like cells in Cluster 17 (CD69^+^PD-1^hi^TIM-3^+^CX3CR1^+^) (Fig. 4E).

Consistent with this, tumor CD8^+^ T cells from prophylactically P1A-vaccinated mice showed reduced co-expression of PD-1 and TIM-3 inhibitory receptors compared to PBS-treated controls, indicating reduced levels of terminal exhaustion (Fig. 4F-G). The small fraction of TIM-3^+^PD-1^+^ terminally exhausted (T_TEX_) cells in P1A-vaccinated tumors were primarily P1A_43-51_-specific (Fig. 4F-G). Therapeutic P1A vaccination similarly reduced tumor T_TEX_ frequencies (Fig. 4H). PD-1 expression was increased with therapeutic but reduced with prophylactic P1A vaccination relative to PBS-treated controls (Fig. 4H), likely reflecting different antigen encounter dynamics. Overall, few CD8^+^ T cells in P1A-vaccinated tumors were T_TEX_, suggesting preserved cytotoxic functionality.

Stem-like TCF-1^+^PD-1^+^ progenitor exhausted (T_PEX_) tumor-infiltrating lymphocytes (TILs) have been linked to vaccine-mediated tumor control^20^. Therapeutic P1A vaccination increased T_PEX_ proportions, while prophylactic P1A vaccination decreased both T_PEX_ and TCF-1^-^PD-1^+^ populations (Supplementary Fig. 8F-H). T_PEX_ levels were inversely correlated with those of immunosuppressive myeloid cells (Arg1^+^CD11b^+^) in tumors (Supplementary Fig. 8I). Among prophylactically vaccinated mice, <25% of P1A_43-51_-specific and TRM-like T cells were TCF-1^+^ compared to ∼40% of total CD8^+^ T cells (Supplementary Fig. 8F-H), consistent with terminal TRM differentiation.

To clarify the association between expression of TRM-associated molecules and other phenotypic markers on vaccine-induced brain TILs, we examined Gpr149_128-135_- and P1A_43-51_-specific T cells stratified by CD103/CD69 expression. DP Gpr149_128-135_-specific cells expressed higher levels of the tissue residency markers CD49a and Runx3 than CD103^-^CD69^+^ single-positive (SP) cells, suggesting that DP cells are more mature TRMs, though CD39 expression was similar (Fig.4I). Meanwhile, SP cells expressed higher levels of markers for stemness (Ly108) and proliferation (Ki67) than DP cells (Fig.4I), suggesting that DP cells may originate from SP cells upon propagation and differentiation. PD-1 expression was higher on DP P1A_43-51_-specific T cells but lower on DP Gpr149_128-135_-specific T cells relative to SP cells (Fig. 4I, Supplementary Fig. 8J-K). DP and SP cells were more similar to each other than to double-negative (DN) cells, which expressed the highest TCF-1 and Ly108 and lowest PD-1 levels (Fig. 4I, Supplementary Fig. 8J-K). Together, these data indicate heterogeneous differentiation states among antigen-specific TILs within the tumor niche.

### CD103^+^CD69^+^ TRM-like CD8^+^ T cells in brain tumors are highly polyfunctional

To assess functionality of antigen-specific CD103/CD69 subsets, brain immune cells from P1A-vaccinated, SB28.P1A-challenged mice were restimulated *ex vivo* with P1A peptides. Stimulation did not alter CD103/CD69 expression on CD8^+^ T cells (Fig. 5A, Supplementary Fig. 9A), though P1A_43-51_-specific T cells were over-represented among the DP subset (Fig. 5B-C). DP cells exhibited superior polyfunctionality, and expressed the most CD107a, granzyme B (GZMB), IFNγ, IL-2, and TNF in isolation (Fig. 5D-E, Supplementary Fig. 9B-C). After accounting for antigen-specific enrichment by gating on P1A_43-51_/H-2D^b^-tetramer^+^ and/or CD3^lo^CD8^+^ cells, DP cells remained significantly more tri-functional than SP cells, with both exceeding DN cells (Supplementary Fig. 9D-F).

**Figure 5.**
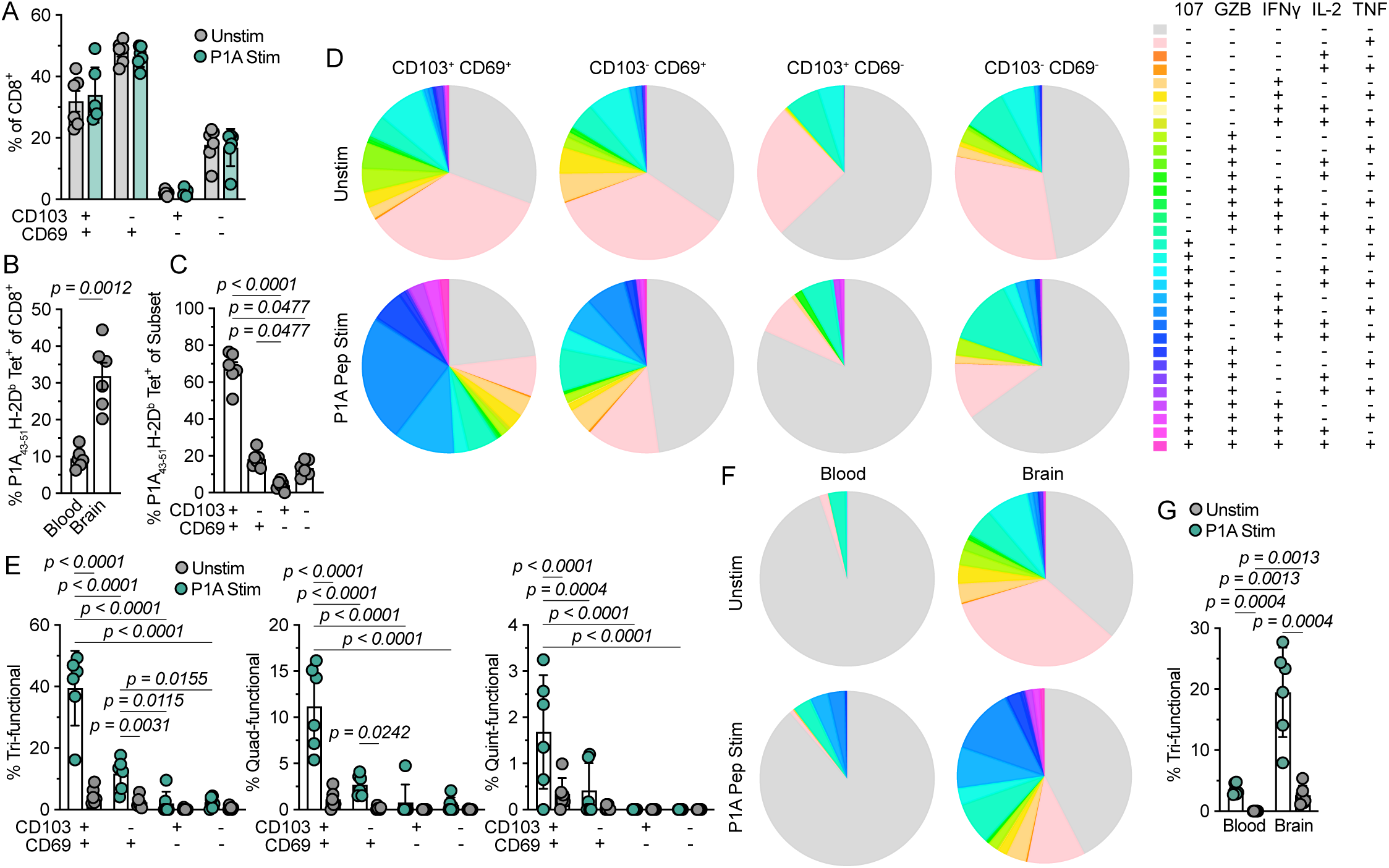
CD103^+^CD69^+^ TRM-like CD8^+^ T cells in brain tumors are highly polyfunctional. For all panels (A-G), mice were prophylactically vaccinated with ChAdOx1/MVA-P1A and challenged with SB28.P1A tumors prior to isolation of immune cells from brains and blood 21 days post-tumor challenge (as in Figure 3A) for *ex vivo* restimulation and flow cytometry. (A) Proportion of CD103/CD69-expressing CD8^+^ T cells in brains with/without stimulation (*n =* 6, all *n.s.*). (B) Brain or blood P1A_43-51_/H-2D^b^-tetramer-binding CD8^+^ T cells without stimulation (*n =* 6). (C) Brain P1A_43-51_/H-2D^b^-tetramer-binding CD8^+^ T cells by CD103/CD69 expression, without stimulation (*n =* 6). (D) Brain CD8^+^ T cells polyfunctionality by CD103/CD69 expression with/without stimulation (*n* = 6). (E) Brain CD8^+^ T cells simultaneously expressing ≥3-5 markers in (D) by CD103/CD69 expression with/without stimulation (*n* = 6). (F) Brain or blood CD8^+^ T cells polyfunctionality with/without stimulation (*rows*), utilizing the color key in (D) (*n* = 6). (G) Brain or blood CD8^+^ T cells simultaneously expressing 3 markers in (F) with/without stimulation (*n* = 6). Data are presented as mean ± SEM for (A-C), and mean ± SD for (E), (G). Statistical analysis was performed using RM two-way ANOVA with Sídǎk’s multiple comparisons test (Stimulated vs. Unstimulated) for (A), two-tailed Welch’s t test for (B), Kruskal-Wallis test with Dunn’s multiple comparisons for (C), and two-way ANOVA with Tukey’s multiple comparisons test for (E), (G).

Untreated SB28.P1A-bearing mice harbored significantly fewer brain CD8^+^ and P1A_43-51_-specific T cells, but their DP cells still trended toward increased polyfunctionality upon stimulation, though stimulation augmented the DP fraction (Supplementary Fig. 9G-K). Finally, irrespective of CD103/CD69 expression, compared to blood-derived CD8^+^ T cells, brain-derived CD8^+^ T cells from vaccinated mice exhibited disproportionately greater polyfunctionality despite enrichment for P1A_43-51_-specificity (Fig. 5B, F-G).

Together, these findings indicate that vaccine-induced CD8^+^ T cells in the brain, and particularly CD103^+^CD69^+^ TRM-like cells, retain high functional capacity to respond to antigen and exert anti-tumor immunity.

### Route of prophylactic vaccination impacts the magnitude of circulating but not brain-infiltrating CD8^+^ T cells

While vaccination can drive CD8^+^ T cells to the CNS in infectious disease settings, the magnitude of antigen-specific TRM-like cells induced by ChAdOx1/MVA vaccination was higher than anticipated. To determine whether induction of tumor-specific TRM-like cells and survival were influenced by vaccination route, we compared IM/IM, IM/IV, and IV/IV administration of ChAdOx1/MVA-P1A at a four-week interval (Supplementary Fig. 10A). As observed previously^7^, IV/IV vaccination induced the highest magnitude of circulating P1A_43-51_-specific T cells (Supplementary Fig. 10B). However, IV/IV-induced P1A_43-51_-specific T cells were significantly less polyfunctional than those induced by IM/IM or IM/IV vaccination (Supplementary Fig. 10C, D). Vaccination route did not significantly affect survival following intracranial SB28.P1A challenge (Supplementary Fig. 10E), consistent with the lack of correlation between circulating antigen-specific CD8^+^ T cells and survival. Notably, vaccination route also did not affect induction of CD8^+^ T cells, TRMs, or P1A_43-51_-specific T cells in tumor-challenged brain hemispheres (Supplementary Fig. 10F). We therefore concluded that IV boosting is not required for induction of brain TRM-like CD8^+^ T cells or survival benefit in orthotopic glioblastoma.

### Both intramuscular and intravenous vaccine boosting drive antigen-specific CD8^+^ T cells to multiple tissues

To assess the tissue trafficking and potential neurotropism of antigen-specific CD8^+^ T cells, we collected twelve tissues from tumor-naïve mice vaccinated IM/IM or IM/IV with ChAdOx1/MVA-P1A. P1A_43-51_-specific T cells were detected across tissues, including liver, lung, meninges, muscle, brain, and skull bone marrow, indicating no preferential neurotropism, with no significant differences in P1A_43-51_-specific CD8αβ^+^ T cell proportions by vaccination route within individual tissues (Fig. 6A-B, Supplementary Fig. 10G-H).

**Figure 6.**
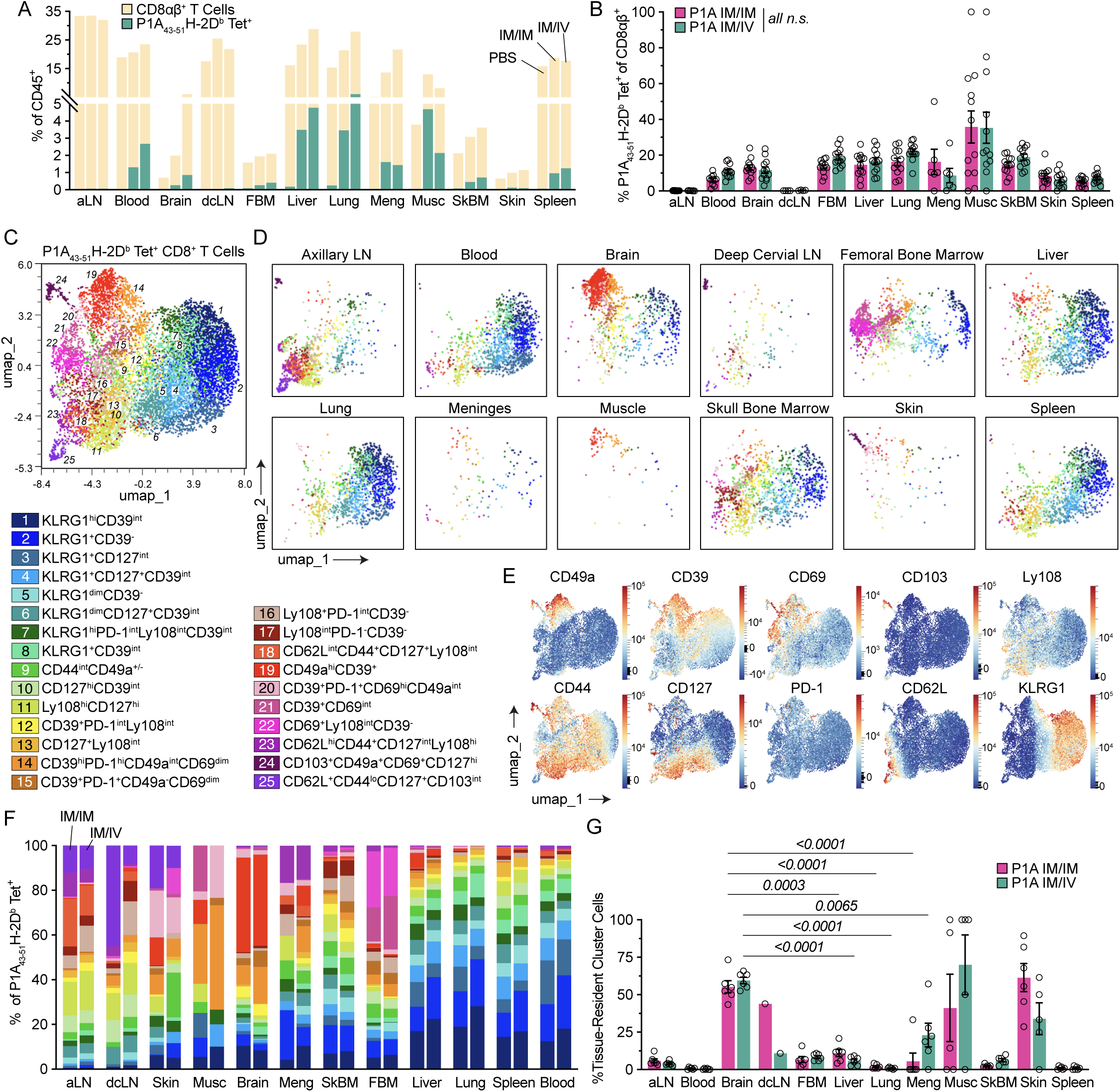
Both intramuscular and intravenous vaccine boosting drive antigen-specific CD8^+^ T cells to multiple tissues. For all panels (A-G), mice were prophylactically vaccinated IM/IM or IM/IV with ChAdOx1/MVA-P1A four weeks apart, and tissues were collected two weeks post-boost for flow cytometry. (A) CD8αβ^+^ or P1A_43-51_/H-2D^b^-tetramer^+^CD8αβ^+^ among CD45^+^ cells by tissue after PBS (*left*), IM/IM vaccination (*middle*), or IM/IV vaccination (*right*) in various tissues (aLN, axillary lymph nodes; dcLN, deep cervical lymph nodes; FBM; femoral bone marrow; Meng, meninges; Musc, muscle; SkBM, skull bone marrow) (*n =* 8-12, pooled from 2 experiments). (B) P1A_43-51_/H-2D^b^-tetramer-binding among CD8αβ^+^ T cells for vaccinated mice (*n =* 8-12, pooled from 2 experiments). (C) UMAP of P1A_43-51_/H-2D^b^-tetramer^+^CD8αβ^+^ T cells from all tissues of vaccinated mice (*n =* 131 total). (D) P1A_43-51_/H-2D^b^-tetramer^+^CD8αβ^+^ T cell distribution by tissue, pooled regardless of vaccination route (*n* = 10-12). (E) Heatmap of phenotypic markers for UMAP in (C) (*n* = 131). (F) P1A_43-51_/H-2D^b^-tetramer^+^CD8αβ^+^ T cells in subsets from (C) by tissue and vaccination route (IM/IM *left*, IM/IV *right*), stacked ascending from bottom (*n =* 5-6). (G) P1A_43-51_/H-2D^b^-tetramer^+^CD8αβ^+^ T cells in tissue-resident-like clusters (i.e. 14, 19, 20, 24) by tissue and vaccination route (*n =* 5-6). Data are presented as mean for (A), (F), and mean ± SEM for (B), (G). Statistical analysis was performed by two-way ANOVA with Sídǎk’s multiple comparisons test (IM/IM vs. IM/IV) for (B), and two-way ANOVA with Tukey’s multiple comparisons for (G) with significant comparisons shown only for brain vs. liver, lung, meninges, and skin within a given vaccination strategy, and brain IM vs. IV (all *n.s.*).

Unsupervised clustering of tissue-derived P1A_43-51_-specific T cells identified 25 subsets with tissue-dependent distributions (Fig. 6C-F). P1A_43-51_-specific T cells from liver, lung, spleen, and blood were phenotypically similar and dominated by KLRG1^+^ SLECs, while brain-derived cells more closely resembled those in muscle and skin and were enriched for TRM markers, including CD49a, CD39, and CD69 (Fig. 6F-I, Supplementary Fig. 10I-M). Consistent with earlier findings in non-tumor-challenged mice, brain P1A_43-51_-specific T cells were primarily CD103^-^, while CD103 expression was highest in skin-derived cells (Supplementary Fig. 10M). Together, these data show that non-local ChAdOx1/MVA immunization drives widespread infiltration of tumor-specific CD8^+^ T cells with tissue-specific phenotypes, supporting its potential to protect against solid cancers in diverse anatomical compartments.

### Antigen-specific TRM-like cells display tissue-specific memory and long-term tissue durability to protect against glioblastoma

To determine whether TILs expressing TRM markers function as sentinel, non-recirculating cells, we assessed tissue-specific immunological memory by comparing control of ectopic and orthotopic secondary tumors. P1A-vaccinated tumor-challenge survivors rechallenged with SB28.P1A either in the flank or original brain hemisphere rejected intracranial but not subcutaneous tumors (Fig. 7A-B). A separate cohort rechallenged simultaneously at both sites all developed flank tumors, but exhibited reduced intracranial tumor uptake compared to naïve controls, confirming tissue-specific memory (Fig. 7C-D).

**Figure 7.**
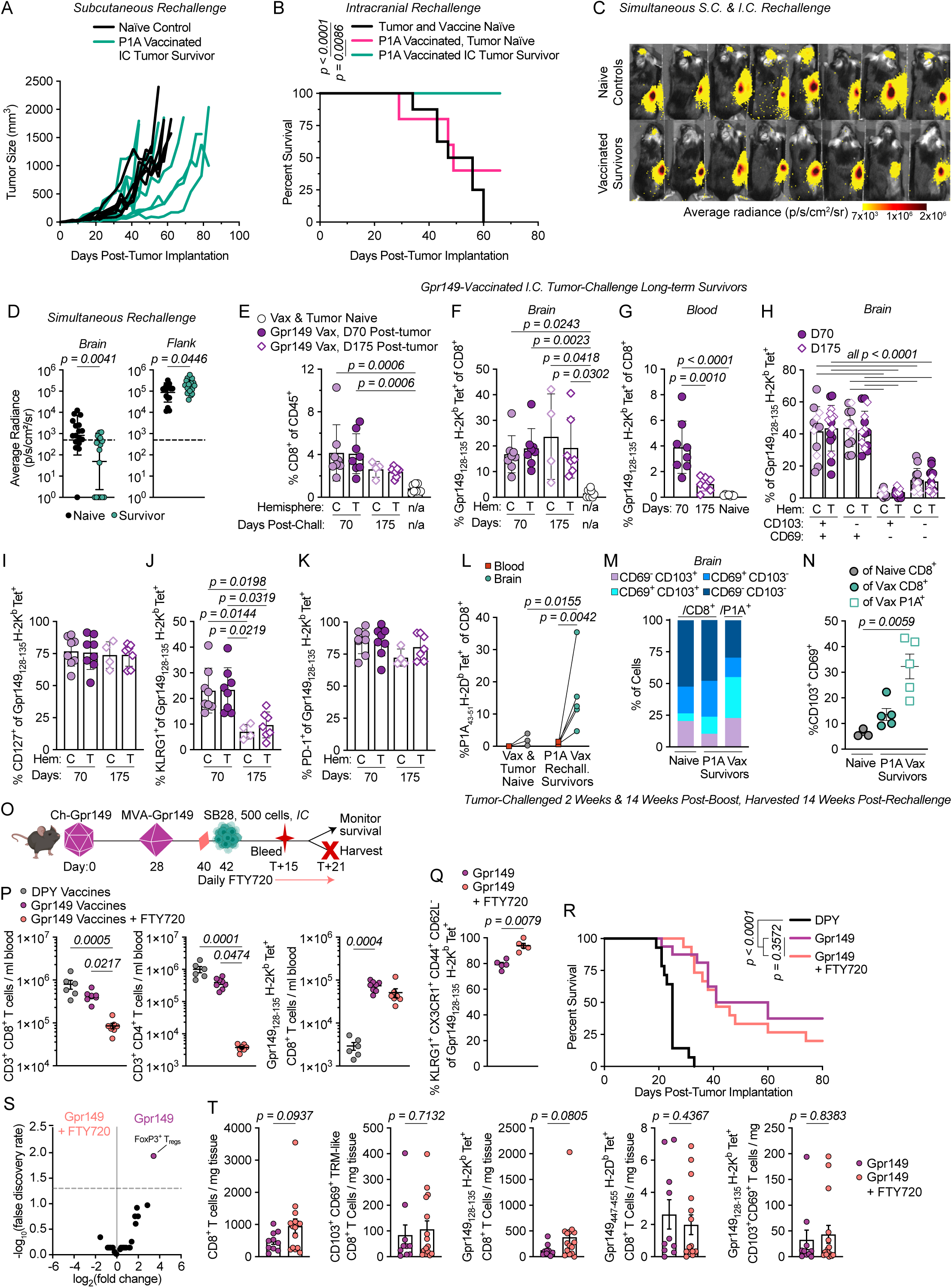
Antigen-specific TRM-like cells display tissue-specific memory and long-term tissue durability to protect against glioblastoma. (A) Individual tumor growth for prophylactically P1A-vaccinated mice surviving IC SB28.P1A tumor challenge or naïve controls, rechallenged 20 weeks post-IC tumor with subcutaneous SB28.P1A flank tumors (*n =* 6-9). (B) Survival for prophylactically P1A-vaccinated mice which survived IC SB28.P1A tumor challenge or were non-challenged, or naïve controls, all rechallenged 12 weeks post-original challenge date with IC SB28.P1A tumors (*n =* 5-9). (C) IVIS images 7 days post-rechallenge for prophylactically P1A-vaccinated mice surviving IC SB28.P1A tumor challenge or naïve controls, rechallenged 15 weeks post-IC tumor with both subcutaneous and IC SB28.P1A tumors (*n =* 8 representative animals, pooled from 2 experiments). (D) D7 intracranial or flank IVIS signal for mice in (C) (*n =* 17, pooled from 2 experiments). For (E-K): brains and blood of prophylactically Gpr149-vaccinated mice surviving IC SB28 challenge were collected 70 or 175 days post-challenge for flow cytometry. T, tumor-challenged brain hemisphere (*dark purple*); C, contralateral hemisphere (*lavender*). (E) Brain CD8^+^ T cells among CD45^+^ cells (*n =* 4-8, pooled from 2 experiments). (F-G) Brain (F) or blood (G) Gpr149_128-135_/H-2K^b^-tetramer^+^ cells among CD8^+^ T cells (*n =* 4-8, pooled from 2 experiments). (H) Brain Gpr149_128-135_/H-2K^b^-tetramer^+^ cells by CD103/CD69 expression (*n =* 12-15, pooled from 2 experiments). (I-K) Phenotypes of brain Gpr149_128-135_/H-2K^b^-tetramer^+^ cells (*n =* 4-8, pooled from 2 experiments). For (L-N), brains and blood of naïve or prophylactically P1A-vaccinated mice surviving 2 consecutive IC SB28.P1A tumor challenges (as in (B)) were harvested 14 weeks post-rechallenge. (L) Blood or brain P1A_43-51_/H-2D^b^-tetramer^+^ cells among CD8^+^ T cells (*n =* 3-5). (M-N) CD103/CD69 expression of brain CD8^+^ or P1A_43-51_/H-2D^b^-tetramer^+^CD8^+^ T cells (*n* = 3-5) (O) Experimental scheme for (P-T). Mice were vaccinated with ChAdOx1/MVA-Gpr149 or -DPY; some mice began receiving daily injections of FTY720 (1 mg/kg) two days prior to intracranial SB28 tumor challenge. (P) Flow cytometry analysis of PBMCs 15 days post-tumor implantation (*n* = 6-8). (Q) Flow cytometry analysis of PBMCs 21 days post-tumor implantation (*n* = 5). (R) Survival of SB28-challenged mice (*n* = 14-15, pooled from 2 experiments). (S) Comparative expression of subsets listed in Supplementary Fig. 7 for tumor-challenged brain hemispheres (*n =* 5). (T) Flow cytometry analysis of brain immune cells 21 days post-tumor implantation (*n* = 10-15, pooled from 2 experiments). Data are presented as geometric mean ± geometric SD for (D), mean ± SD for (E-K), mean for (M), and mean ± SEM for (N, P-Q), (T). Statistical analysis was performed using log-rank test for (B), (R), lognormal Welch’s t test for (D), Kruskal-Wallis test with Dunn’s multiple comparisons for (E-F), (I-K), (N), (P), one-way ANOVA with Tukey’s multiple comparisons for (G), two-way ANOVA with Tukey’s multiple comparisons for (H), two-way ANOVA with Fisher’s LSD multiple comparisons test for (L), edgeR for (S), and two-tailed Mann-Whitney test for (Q), (T).

To evaluate the long-term persistence of brain TRM-like cells, we profiled immune cells from surviving Gpr149-vaccinated mice 70 or 175 days after intracranial SB28 challenge. Survivors maintained higher levels of brain CD8^+^ T cells than naïve controls, driven by Gpr149_128-135_-specific T cells, which comprised nearly 20% of all CD8^+^ T cells in both contralateral and tumor-challenged hemispheres (Fig. 7E-F). While frequencies of Gpr149_128-135_-specific T cells remained stable in the brain between days 70 and 175, they declined in the blood, indicating superior durability of brain-resident cells (Fig. 7F-G). Most brain Gpr149_128-135_-specific T cells were DP or SP TRMs, with stable CD103/CD69 phenotypes over time (Fig. 7H). The majority expressed PD-1 and CD127 at both timepoints, while the KLRG1^+^ minority declined significantly from day 70 to day 175, consistent with loss of SLECs (Fig. 7I-K). Thus, vaccine-induced antigen-specific TRM-like cells persist long-term in brain tissue following intracranial tumor challenge.

We next examined the durability of antigen-specific CD8^+^ T cells in P1A-vaccinated mice surviving multiple intracranial tumor challenges. At 200 days post-MVA-P1A boost, the frequency of P1A_43-51_-specific CD8^+^ T cells in the brain was over 20-fold higher than in the blood (Fig. 7L). The proportion of DP cells among P1A_43-51_-specific T cells exceeded that of total CD8^+^ T cells in naïve controls, and a subset of P1A_43-51_-specific T cells were CD103^+^CD69^-^ (Fig. 7M-N). Together, these findings show that vaccinated mice maintain functional brain TRM-like cells long after tumor exposure.

To determine whether antigen-specific CD8^+^ T cells persist in the brain without tumor challenge, we analyzed immune cells from tumor-naïve, P1A-vaccinated mice 12 weeks post-boost. The frequency of P1A_43-51_-specific CD8^+^ T cells remained higher in the brain than blood (Supplementary Fig. 10N), despite declining by approximately half relative to levels 5 weeks post-boost (Fig. 3G). Brain P1A_43-51_-specific CD8^+^ T cells were predominantly SP at both timepoints, though the DP subset was increased at 12 weeks (Supplementary Fig. 10O, Fig. 3J). Therefore, without tumor exposure, ChAdOx1/MVA-induced antigen-specific T cells gradually decline yet persist in the brain, accompanied by increasing CD103 and decreasing CD69 expression.

### Vaccination seeds the brain with tumor-lysing CD8^+^ T cells to protect against glioblastoma without the need for further immune cell recruitment

We next evaluated whether seeding brains with antigen-specific CD8^+^ T cells can protect against intracranial tumors without the recruitment of circulating cells. ChAdOx1/MVA-Gpr149-vaccinated mice were treated with FTY720 to block lymphocyte egress from lymphoid tissues (Fig. 7O). FTY720 significantly reduced blood levels of CD4^+^ and CD8^+^ T cells, though it did not significantly reduce Gpr149_128-135_-specific T cells, likely because these cells were predominantly CD44^+^CD62L^-^ (Fig. 7P-Q).

However, these cells were almost exclusively KLRG1^+^ (Fig. 7Q), and CD103^+^ TRMs are known to derive from cells lacking KLRG1^21^. FTY720 did not diminish the prolonged survival of Gpr149-vaccinated mice challenged with SB28 tumors, indicating that brain-resident T cells were sufficient for protection, while circulating CD8^+^ T cell recruitment was dispensable (Fig. 7R). Foxp3^+^ T_reg_ cells were the only immune subset in tumor-challenged brains altered by FTY720 treatment, indicating that SB28 typically recruits T_reg_ cells from the circulation (Fig. 7S, Supplementary Fig. 7A-C, K). FTY720 treatment did not impact the magnitude of CD8^+^ T cells, Gpr149_128-135_-specific cells, or TRM-like cells, suggesting that DP TILs arise from SP cells seeded before tumor implantation and mediate anti-tumor immunity independently of circulating T cell recruitment (Fig. 7T, Supplementary Fig. 7L).

### Antigen-specific brain TRM-like cells are sufficient to protect against glioblastoma

Finally, we asked whether antigen-specific TRM-like cells were sufficient to mediate anti-glioblastoma immunity by intracranially adoptively transferring sorted CD8^+^ T cells to naïve recipients before tumor challenge (Fig. 8A). Strikingly, brain-derived DP P1A_43-51_-specific TRM-like cells were sufficient to prolong survival of tumor-challenged mice, while brain-derived non-P1A-specific CD8^+^ T cells and blood-derived DN P1A-specific CD8^+^ T cells were not (Fig. 8B). This demonstrates that antigen-specific CD103^+^CD69^+^ brain TRM-like cells induced by ChAdOx1/MVA vaccination are sufficient to protect against glioblastoma and exert superior anti-tumor immunity to circulating DN tumor-specific T cells.

**Figure 8.**
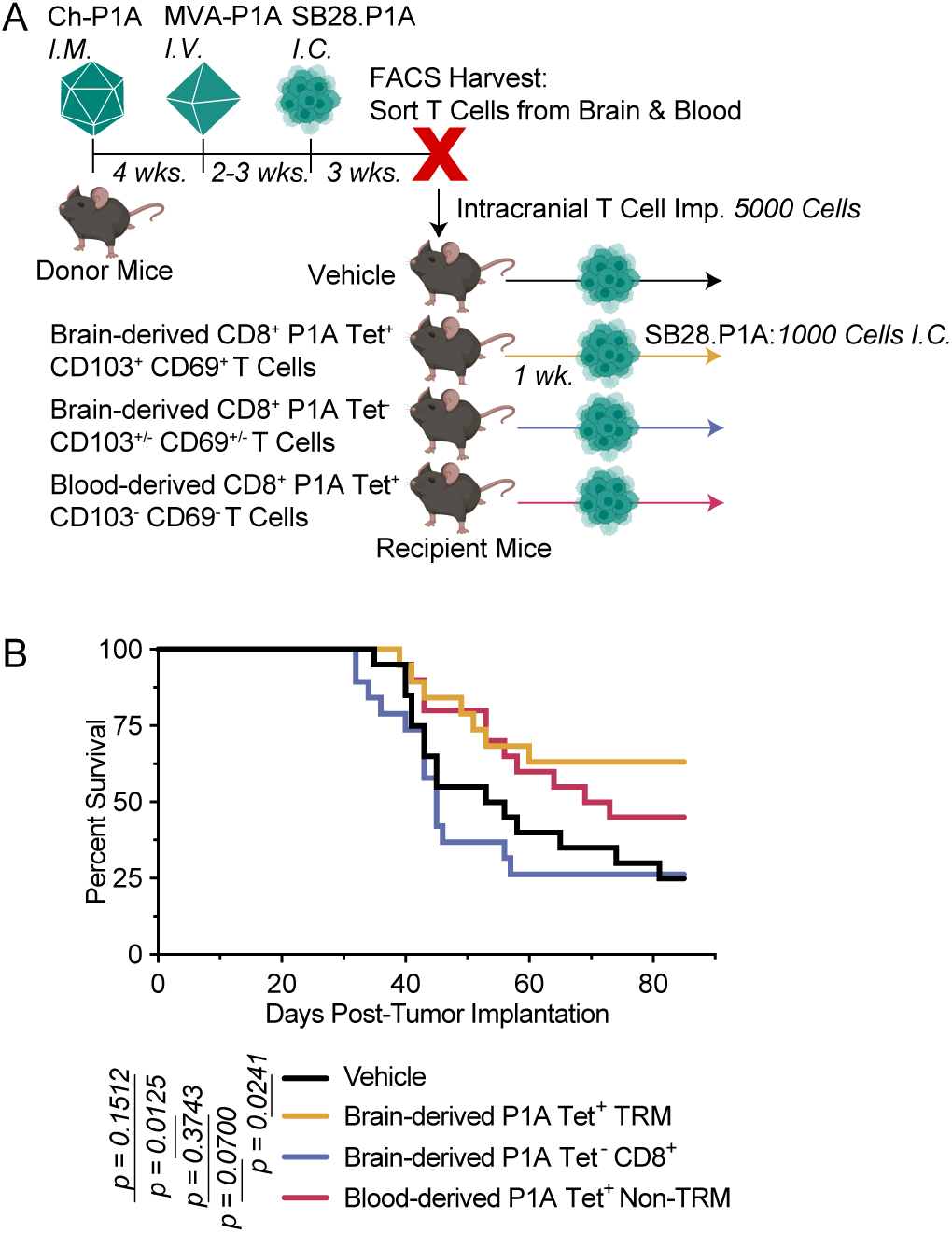
Antigen-specific brain TRM-like cells are sufficient to protect against glioblastoma. (A) Experimental scheme: specific CD8^+^ T cell subsets from the brains and blood of P1A-vaccinated, tumor-challenged mice were adoptively transferred into the brains of naïve recipients. Recipients were challenged with SB28.P1A tumors in the same intracranial location one week later. (B) Survival of SB28.P1A-challenged recipient mice (*n =* 19-20, pooled from 2 experiments).

## Discussion

Current vaccination approaches have failed to demonstrate clinical benefit in phase III trials for patients with glioblastoma, likely reflecting insufficient induction of tissue-infiltrating CD8^+^ T cells^12^. While it was unknown whether ChAdOx1/MVA vaccination could drive CD8^+^ T cells to the brain, accumulating evidence indicates that TRMs populate the brain post-immunization or post-infection, impacting neuroimmunity^22–30^.

TRMs have been identified in human glioblastomas and can mediate protection against CNS infection and tumor progression in experimental models^15,31–36^. However, clinically deployable vaccination strategies to induce brain TRMs and meditate therapeutic benefit against glioblastoma remained unidentified. Here, we demonstrate that ChAdOx1/MVA vaccination induces tumor-specific TRM-like CD8^+^ T cells in the brain that are sufficient to mediate anti-glioblastoma immunity.

These findings shift the framework for evaluating glioblastoma vaccines, underscoring tissue-residency, rather than the magnitude of circulating immune cells, as a determinant of efficacy. Circulating CD8^+^ T cell frequency and phenotype did not correlate with outcomes and did not reflect brain-infiltrating populations. Instead, vaccine-induced TRM-like cells more closely aligned with tumor control, consistent with evidence in other cancers linking TRMs to improved survival^12,13,36^. ChAdOx1/MVA vaccination is a clinically feasible, non-invasive strategy to drive antigen-specific TRM-like cells to the brain.

We demonstrate prophylactic and therapeutic efficacy of ChAdOx1/MVA vaccination against glioblastoma in multiple murine antigen contexts. Identification of the SB28 TAA Gpr149 enabled endogenous antigen-targeted vaccination in this immunologically stringent, ICB-refractory glioblastoma model, avoiding reliance on xenogeneic or supra-physiologically expressed transgenic murine antigens that may lead to overestimation of therapeutic efficacy. Targeting endogenous Gpr149 establishes a more rigorous standard for preclinical evaluation of antigen-targeted therapies in immunologically cold tumors and provides a resource for future mechanistic studies.

While Gpr149 is not upregulated in human glioblastoma, multiple shared TAAs, private and shared neoantigens, alternative splice variants, and non-canonical “dark genome” peptides have emerged as clinically relevant vaccine targets^37–47^. ChAdOx1/MVA vectors can encode large or multiplexed antigenic sequences, providing flexibility to accommodate inclusion of diverse antigen classes in future glioblastoma vaccines.

Surprisingly, ChAdOx1/MVA vaccination was effective as a monotherapy against glioblastoma. Unlike in other cancer models^6,8,9^, adjuvant ICB provided no additional benefit. While the clinical failure of both vaccines and ICB in glioblastoma has been presumed to be driven by a lack of combinatorial use, our data suggest that ChAdOx1/MVA-induced TRM-like cells do not require ICB licensing to exert cytotoxicity. However, vaccination efficacy may be further augmented by combinatorial strategies harnessing agents with particular potency in the brain TME, such as α2-adrenergic receptor agonists^48^, regulators of nonconventional T cells^49–51^, and neuromodulators^52^. Vaccination remodeled the tumor microenvironment, reducing immunosuppressive myeloid and T_reg_ populations. How vaccine-induced CD8^+^ TRMs interact with these compartments remains unresolved, but combinatorial therapies manipulating these interactions may further enhance efficacy.

The heterologous ChAdOx1/MVA platform offers several advantages, including strong CD8^+^ T cell priming, established safety in humans, and capacity for large antigenic inserts^9,53–55^. Despite increased population-level anti-vector immunity following widespread deployment of ChAdOx1-based COVID-19 vaccines, heterologous boosting, alternative routes of vaccination, and appropriately spaced dosing should preserve robust cellular immune responses. Notably, peripheral ChAdOx1/MVA vaccination drove antigen-specific T cell infiltration across multiple tissues, including the brain, without requiring local administration, distinguishing this approach from strategies that rely on site-restricted immunization to promote tumor control^56–58^.

The delineation of TRM-like TILs from both exhausted T cells and conventional TRMs remains an active area of investigation^13,59–61^. We observed that antigen-specific CD8^+^ T cells seeded in the brain were predominantly CD103^-^CD69^+^CD49a^+^ prior to tumor challenge and acquired persistent CD103 expression following tumor encounter. Tumor-derived factors including TGF-β^62^ and local antigen presentation likely contribute to this differentiation, consistent with previous work linking CD103 induction to local cues and TCR engagement^30^. Importantly, these TRM-like cells were polyfunctional and not terminally exhausted, distinguishing them from dysfunctional TRM-like TILs described in other contexts^60,61^. Why antigen-specific TRM-like cells appear in contralateral and tumor-challenged hemispheres at similar frequencies remains unresolved, but may reflect facile cell trafficking or dispersal of tumor antigens and factors, following a pattern distinct from tissues such as the mammary fat pad, where local breast cancer-induced TRMs do not migrate to confer contralateral protection^63^.

Though a prior study showed that intracranial transfer of bulk CD8^+^ T cells with TRM-like phenotypes derived from the brains of glioma-surviving mice can protect naïve mice against glioma challenge^35^, our findings demonstrate that TRM tumor-specificity is essential for effective immunity. Only antigen-specific and not bystander CD8^+^ T cells derived from tumor-challenged brains conferred protection, and antigen-specific CD103^+^CD69^+^ cells from tumor-challenged brains provided superior protection to circulating antigen-specific DN cells. These observations have important implications for adoptive cell therapies targeting brain tumors, and are consistent with preclinical work demonstrating that polarization of chimeric antigen receptor (CAR)-T cells toward TRM-like phenotypes improves anti-tumor efficacy^64^.

Our findings establish a key role for vaccine-induced TRM-like CD8^+^ T cells in glioblastoma control, though several considerations require further investigation. Preclinical evaluation of prime-boost vaccination strategies in aggressive transplantable tumor models is inherently constrained by limited therapeutic windows. As durable immunological memory typically develops over weeks, our study was not designed to fully resolve the contribution of TRM-like cells to therapeutic treatment. While human glioblastoma is extremely aggressive, the window available to treat patients exceeds that of our models, raising the possibility that extended prime-boost vaccination of patients could drive antigen-specific CD8^+^ T cells to the brain which would adopt a TRM-like phenotype with time. Future studies will be required to define the ontogeny, kinetics, and functional maturation of vaccine-induced TRM-like cells during therapeutic vaccination.

Overall, we show that viral vector vaccination induces antigen-specific, polyfunctional TRM-like cells in the brain with potent anti-tumor activity. Seeding the brain with TRM-like cells may enable treatment of glioblastoma and other primary brain tumors, and could extend to the treatment or prevention of brain metastasis and neurotropic infections. These findings provide a rationale for clinical investigation of ChAdOx1/MVA vaccination to treat patients with glioblastoma.

## Acknowledgements

We thank Ferenc Livac, Shafiuddin Siddiqui, and Kathy McKinnon at the Center for Cancer Research Flow Cytometry Core (NIH); Claire Powers at the Pandemic Sciences Institute Viral Vector Core Facility (University of Oxford); V. Clark and H. Gray at the Functional Genomics Facility (University of Oxford), and Devorah Gallardo at the Building 37 Animal Facility (NIH) for technical assistance. We thank Grégoire Altan-Bonnet, Remy Bosselut, Ronald Germain, and Giorgio Trinchieri for critical reading.

## Author Contributions

EES, CSKL, BJVdE, and MT conceived and designed the experiments. EES, LL, TH, AH, MC, JS, JM, VPA, AW, SAM, LN, CHW, JH, SP, HK, BA, CM, NB, MZ, WZ, DD, HS, and MT performed the experiments. EES, LL, TH, and MB analyzed data. BJVdE and MT acquired project funding. HO and MG contributed resources. CSKL, BJVdE, and MT supervised the study. EES administered the project, performed data visualization, and wrote the original manuscript draft. All authors reviewed and edited the manuscript.

## Methods

### Mice

Female 6-8-week-old C57BL/6 and BALB/c mice were purchased from Charles River (USA) or Envigo (UK). Mice received food and water ad libitum and were maintained in 12 hour light/dark cycles. All animal procedures reported in this study that were performed by NCI-CCR affiliated staff were approved by the NCI Animal Care and Use Committee (ACUC) and in accordance with federal regulatory requirements and standards. All components of the intramural NIH ACU program are accredited by AAALAC International. All animal procedures reported in this study that were performed at the University of Oxford were carried out in accordance with the terms of the UK Animals (Scientific Procedures) Act Project License (PPL) (PB050649E).

### Cell culture

The SB28 glioblastoma model was induced via intraventricular transfection of sleeping beauty (SB)-flanked mPDGF, NRasV12, Luciferase, and shRNA silencing of p53; a cell line was established from the tumor^65^. SB28 cells were cultured in DMEM supplemented with 10% fetal bovine serum, 100 U/ml penicillin, 100 μg/ml streptomycin, 2 mM L-glutamine, 100 mM Sodium pyruvate and 50 μM 2-mercaptoethanol. GL261 and HEK293T were maintained in DMEM supplemented with 10% FBS, 100 U/ml penicillin, 100 μg/ml streptomycin and 2mM L-glutamine. DC2.4 cells were maintained in RPMI 1640 supplemented with 10% fetal bovine serum, 100 U/ml penicillin, 100 μg/ml streptomycin, 2 mM L-glutamine, 1% Non-essential amino acids, 0.01 M HEPES Buffer Solution, and 50 μM 2-mercaptoethanol. All adherent cell lines were harvested via trypsination with 0.05% trypsin. T cell media was composed of RPMI 1640 supplemented with 10% FBS, 100 U/ml penicillin, 100 μg/ml streptomycin, 2 mM L-glutamine, 0.01 M HEPES, 100 mM Sodium pyruvate, 1% Non-essential amino acids, 50 μM 2-mercaptoethanol, and 20 U/ml rhIL-2 (Peprotech). All cell lines were maintained at 37°C 5% CO_2_. Cells were negative for mycoplasma contamination. Cell lines were tested negative for pathogen contamination prior to *in vivo* implantation.

### Generation of SB28.P1A cell line

pEF4/V5-HisA-P1A plasmids were diluted in Opti-MEM (Gibco) containing Lipofectamine 2000 (ThermoFisher) and cultured with SB28 target cells for 8 hours. Cells were then maintained in selection media containing neomycin for more than three weeks before manual single-cell plating, expansion of monoclonal cells, and validation of P1A expression.

### Western blot

Protein lysate was obtained from cells using RIPA buffer (Sigma) supplemented with 1X protease inhibitor (Roche) and 1X phosphatase inhibitor (ThermoFisher). Protein concentrations were determined using the Pierce BCA Assay Kit (ThermoFisher) and equal amounts of protein were prepared with NuPAGE LDS Sample Buffer (Invitrogen) and NuPage Reducing Agent (Invitrogen). Samples were run on a NuPage 4-12% Bis-Tris Gel (Invitrogen) in NuPage MES running buffer, then transferred to polyvinylidene difluoride membranes (Bio-Rad) with the Trans-Blot Turbo Transfer System (Bio-Rad). Membranes were subsequently blocked in 5% milk in phosphate buffered saline (PBS) with 1% Tween-20, followed by primary antibody (mouse anti-P1A, 1:1,000, LICR Brussels; rabbit anti-vinculin, 1:10,000, Cell Signaling) incubation overnight at 4°C and secondary antibody (anti-mouse IgG-HRP, 1:5,000, Santa Cruz; anti-rabbit IgG-HRP, 1:10,000, Cell Signaling) incubation for 30 minutes at room temperature. Blots were developed using the SuperSignal West Femto Chemiluminescent Substrate (ThermoFisher), and imaged on a Bio-Rad ChemiDoc Imager.

### Tumor implantations and survival

Intracranial tumors were implanted into mice anesthetized with a combination of xylazine (20 mg/ml) and ketamine (100 mg/ml) diluted in 0.9% NaCl at a 1:1:4 volume ratio, at a total dose of 0.1ml per 20g body weight. Cells in 2 μl of HBSS/DNase (200 μg/ml, Roche) were implanted stereotactically through a 25G Burr hole in the skull using a 33G Hamilton Syringe into the right parietal lobe, 4 mm anterior and 2 mm lateral to the bregma, at a depth of 2 mm, over a period of 10 minutes. Number of intracranially injected cells were as follows: 500 SB28 for prophylactic experiments, 250 SB28 for therapeutic experiments, and 1000 SB28.P1A for all experiments. Cells were pre-treated in vitro with 100 U/ml of IFNγ (Peprotech) for 48 hours prior to implantation for all experiments, unless otherwise specified. Mice were euthanized at humane endpoint when body weight fell below 15% of maximum body weight, or when dorsal cranial expansion resulting from tumor outgrowth exceeded a volume of 300 mm^3^ upon measurement with calipers. For experiments involving collection of brain tissue, “Non-challenged” controls were sham injected intracranial with vehicle only. Subcutaneous tumors (500,000 SB28.P1A cells) were implanted into the right flank of mice anesthetized with isoflurane. Subcutaneous tumor growth was measured 2–3 times per week with a caliper, and mice were euthanized when tumor size reached a mean diameter of 15 mm. Tumor volume and cranial expansion volume (V) were calculated according to the formula: V=((length (mm)×width^2^ (mm))×0.52). For rechallenge experiments, age-matched naïve controls were used.

### IVIS imaging

*In vivo* tumor measurement was enabled by expression of the luciferase transgene in SB28 cells^2^. D-Luciferin Potassium Salt (GoldBio) was dissolved in PBS (30 mg/ml) prior to use *in vivo*. Mice were injected IP with 100 μl Luciferin 10 minutes prior to imaging with an In Vivo Imaging System (IVIS) Lumina Series III (Revvity). Images were processed using Living Image software (Revvity). Background average radiance was subtracted from the average radiance centered on the head of each mouse. Any negative values resulting from background subtraction were set to 1 for data visualization. For therapeutic experiments, only mice with an IVIS signal above the reliable detection threshold (500 p/s/cm^2^/sr, indicated by dashed line on figures) on day 7 post-tumor implantation were used for experiments. Tumor-cleared mice were defined as having an IVIS signal below the detection threshold. Long-term survivor mice were defined as mice surviving at least 70 days post-tumor implantation with an IVIS signal below the detection threshold.

### Whole exome sequencing and identification of variants

DNA was extracted from cultured SB28 cells, cultured GL261 cells, and C57BL/6 brain tissue using the Qiagen DNeasy kit according to manufacturer’s protocols. Whole-exome sequencing was performed in duplicate with Illumina MiSeq at the NCI Sequence Core Facility. Bioinformatic analysis was performed using the Illumina DRAGEN pipeline (07.021.595.3.7.5). Variant calling was performed using Dragen Post Somatic Calling Filtering (Illumina) for all pairwise permutations of SB28 vs. brain replicates or GL261 vs. brain replicates. Variants passed if detected in at least two comparisons. The impact of the nucleotide variants was assessed using the Ensembl Variant Effect Predictor (EMBL European Bioinformatics Institute) and passed if the outcome was “High” or “Moderate.” Only those variants with a Variant Allele Frequency of ≥ 30% were considered for neoantigen analysis. Analysis was streamlined using the Mouse nEoanTigen pRedictOr (METRO) pipeline code^66^.

### RNA sequencing and identification of Tumor-Associated Antigen (TAA) candidates

RNA was extracted from cultured SB28 cells, cultured GL261 cells, C57BL/6 brain tissue, and mouse astrocytes using the Qiagen RNeasy kit according to manufacturer’s protocols. RNAseq was performed in triplicate for each sample by the Center for Cancer Research Sequencing Facility on NovaSeq 6000 SP using Illumina Stranded mRNA Prep and paired-end sequencing. Reads of the samples were trimmed for adapters and low-quality bases using Cutadapt before alignment with the reference genome (mm10) and the annotated transcripts using STAR. Gene expression quantification analysis was performed for all samples using STAR/RSEM tools. Post-alignment QA/QC, quantification to annotation model, and normalization were performed using Partek (Illumina). Only protein-coding and non-mitochondrial genes were included for consideration as TAAs. A student’s t-test comparing the TPM of triplicates for SB28 or GL261 vs. brain or astrocytes was performed to calculate p-values for each gene. The Benjamini-Hochberg procedure was performed on the p-value stacks for each comparison (SB28 vs. brain, GL261 vs. astrocytes, etc.) to calculate q-values and control the false discovery rate. Only those genes with q<0.05 were considered for further analysis. Genes were then further stratified for consideration based on absolute mean TPM (>0.5 in tumor cells, <0.5 in astrocytes and brain).

### RT-qPCR

Tissue samples were collected immediately upon animal euthanasia and snap-frozen in liquid nitrogen. RNA was extracted from samples using the RNeasy kit (Qiagen) according to manufacturer’s instructions. Total cellular RNA (0.5 μg) was used to synthesize first single-strand cDNA using the SuperScript III First-Strand Synthesis kit (ThermoFisher) according to the manufacturer’s protocol. The QuantiTect SYBR Green PCR kit (Qiagen) was used to quantify cDNA by qPCR according to the manufacturer’s instructions. Reactions were run in triplicate on the StepOnePlusTM Real-Time PCR System (Applied Biosystems) at the following conditions: 95°C for 15 mins, (94°C for 15s, 60°C for 30s, 72°C for 30s) x 40 cycles. Gene expression at the mRNA level for each target gene was quantified relative to an internal housekeeping gene control (*Tbp).* To determine relative mRNA expression, mean ΔCt (difference in cycle threshold number) was calculated for each target gene relative to *Tbp* using the formula ΔCt = Ct Target gene - Ct Housekeeping gene. Relative mRNA expression data are shown as 2^-ΔCt^. The primers used are shown in the Supplemental Data Table 3B.

### Evaluation of TAA Candidate Expression in Human Glioblastoma

RNA-sequencing data (TPM) and clinical annotations for glioblastoma were taken from TCGA-GBM through the Genomic Data Commons. The dataset included 295 tumor samples with RNA-seq data and 617 clinical records. Normal brain cortex expression values were taken from the GTEx dataset (n=270). All tumor samples were confirmed as glioblastoma based on clinical annotations. Nine samples were technical replicates from the same patients. Expression values for duplicates were averaged. After matching expression and clinical data and removing replicates, 286 patients remained for expression analysis. Human orthologs of murine TAAs (SHOX2, FBN2, CDH6, IGF2BP1, and GPR149) were identified using Ensembl gene orthology annotations. For each candidate antigen, fold-change was calculated as the ratio of median glioblastoma expression to median normal brain expression. Prevalence was the percentage of patients with expression above 1 TPM. Tumor expression was compared to the normal tissue median using one-sample t-tests (one-sided, tumor > normal). Multiple testing correction used the Benjamini-Hochberg FDR across five genes, with FDR <0.05 considered significant. Analyses were performed in Python 3.12.10 using pandas v2.3.1, numpy v1.26.4, scipy v1.13.1, and statsmodels v0.14.4. All tests were two-sided unless stated.

### Prediction of MHC binding

For neoepitopes, FASTA sequences corresponding to up to 13 amino acids up- and down-stream of the SNV were input into NetMHC4.1 and the IEDB Consensus Algorithms, which were used to predict 8-14mer epitopes binding to H-2K^b^ and H-2D^b^. For TAAs, FASTA sequences corresponding to the entire amino acid sequence for each gene were input into NetMHC4.1 and the IEDB Consensus Algorithms, which were used to predict 8-11mer epitopes binding to H-2K^b^ and H-2D^b^. All predicted TAA epitopes with an EL Rank < 0.5 (NetMHC4.1) and a Consensus Percentile Rank <1.5 (IEDB) were included for testing in stimulation assays. For prediction of MHC-II binding, Fbn2 minigene sequences were input into NetMHC4.1 and the IEDB Consensus Algorithms, which were used to predict 8-11mer epitopes binding to H-2K^b^ and H-2D^b^.

### Creation of ChAdOx1/MVA vectors expressing TAAs

The coding sequences for TAA constructs were generated via GeneArt Gene Synthesis (Thermo Fisher Scientific) and codon-optimized for mouse, or were PCR-amplified from SB28 cDNA in the case of Igf2bp1 and Shox2 using the Platinum SuperFi II PCR Master Mix (Invitrogen) and linked together using InFusion cloning (Takara). For all constructs, a sequence coding for the 26 amino acid transmembrane domain of the MHC-II invariant chain was linked to the N terminus of the transgenes. Sequences were inserted into the E1 locus of ChAdOx1 under a CMV immediate early promoter. MVA vectors were constructed with the F11 promoter driving transgene expression.

Sequences of cloning vectors were confirmed by Sanger Sequencing (Eurofins Genomics) and alignment in Snapgene. Viruses were produced by the Viral Vector Core Facility in the Pandemic Sciences Institute at the University of Oxford. The purity and identity of the viral vectors were confirmed by PCR. Creation of the P1A-expressing vectors was described previously^6^.

### Vaccinations and checkpoint inhibitor treatments

Unless otherwise specified, for prophylactic vaccination experiments, mice received 5×10^8^ infectious units (IU) of ChAdOx1 virus diluted in PBS injected IM in 50 μl total volume, followed 28 days later by 1×10^7^ plaque forming units (PFU) of MVA virus diluted in PBS injected IV in 100 μl total volume. For therapeutic vaccination experiments, mice received 5×10^8^ IU ChAdOx1 virus injected IM in 50 μl total volume, followed 7-14 days later by 1×10^7^ PFU MVA virus injected IV in 100 μl total volume. In other cases, vaccinations were administered as described, with IV injections always administered in 100 μl total volume and IM injections administered in 50 μl total volume. Checkpoint inhibitors (ICB) were administered intraperitoneally (IP) at a dose of 100 μg/dose in a total volume of 100 μl each. Antibodies were purchased from BioXcell (anti-PD-1 clone 29F.1A12 or Clone RPMI-14; anti-CTLA-4 clone 9H10 or 9D9). In all experiments using ICB, all non-ICB treated mice received 200 μg of rat IgG2a (Sigma) or BioXcell (BE0089) at ICB administration timepoints. In prophylactic vaccination experiments, anti-CTLA-4 and anti-PD-1 were administered on days 4, 7, 11, and 14 post-tumor implantation. For prophylactic vaccination experiments testing groups with or without anti-CTLA-4 and anti-PD-1 (ICB), vaccinated mice were randomized into ICB treatment groups to equalize based on frequencies of antigen-specific CD8^+^ T cells in blood prior to tumor challenge. In therapeutic vaccination experiments, ICB was administered on days 13, 20, 27, and 34 post-tumor implantation. For therapeutic vaccination experiments testing immunization with or without ICB, vaccinated mice were randomized into ICB treatment groups to equalize based on tumor size as measured by IVIS signal on the day 13.

### Perfusion

Prior to the collection of brain tissue for isolation of immune cells, mice received intracardiac perfusion of PBS using a Masterflex Ismatec pump (Avantor) at a rate of 0.15 ml/minute. Mice were injected with an overdose of ketamine/xylazine anesthetic prior to perfusion. In cases where liver was also collected, perfusion was performed using HBSS supplemented with 0.5 mM EDTA.

### Processing of blood and tissues for the isolation of immune cells

Brain collection was performed on day 21 post-tumor implantation in the prophylactic tumor setting and on day 28 post-tumor implantation in the therapeutic tumor setting unless otherwise specified. Blood was collected retro-orbitally in tubes containing EDTA (12.5 mM, Sigma). Two rounds of red blood cell lysis were performed using a buffer comprised of NH_4_Cl, KHCO_3,_ and tetrasodium EDTA (pH 7.3) prior to washing with PBS. Spleens were mashed over a 70 μm strainer or between two glass slides prior to 5 minutes of ACK lysis (Quality Biological) and washing with RPMI 1640. Cerebellums were cut away from brains prior to separation of hemispheres. Brains were dissociated with the GentleMACS Dissociator (Miltenyi) in RPMI 1640 containing 175 U/ml Collagenase Type I (Gibco) and 20 μg/ml DNase I (Roche) at 37°C prior to straining through a 70 μm strainer, followed by debris removal via density centrifugation with 30% Percoll (Cytiva) and washing with RPMI 1640. Skin was collected from ears with dorsal and ventral halves separated manually. Skin was digested dermal side down in RPMI 1640 supplemented with 0.5 μg/ml DNase I and 0.5 mg/ml Liberase TL at 37°C, followed by mechanical dissociation with the GentleMACS Dissociator (Miltenyi) and straining through a 70 μm strainer and washing. Lungs were collected without trachea and esophagus and manually dissociated with scissors prior to incubation with Collagenase Type IX and DNase I at 37°C, followed by straining through a 70 μm strainer, 5 minutes of ACK lysis (Quality Biological), and washing with RPMI 1640.

Muscle was collected from the thigh (IM vaccination site) and manually dissociated with scissors prior to incubation with 0.2 U / ml Liberase TM and 13 μg/ml DNase I at 37°C in DMEM, followed by straining through a 70 μm strainer, layering over 100% lymphoprep for density centrifugation, and washing with DMEM. Meninges were isolated manually using forceps under a stereoscope from intact skull prior to dissociation with 1 mg/ml Collagenase Type IV and 20 μg/ml DNase I at 37°C in RPMI, followed by straining through a 35 μm strainer and washing with RPMI 1640. Skull bone marrow was collected from calvaria after removal of meninges. The calvaria was manually cut into small fragments using scissors followed by multiple rounds of spin-extraction in a tabletop centrifuge (Eppendorf), followed by 30 seconds of ACK lysis (Quality Biological) and washing with PBS. Femoral bone marrow was collected via spin-extraction in a tabletop centrifuge (Eppendorf), followed by 1 minute of ACK lysis (Quality Biological) and washing with PBS. Lymph nodes were dissociated manually using microscope slides followed by straining through a 70 μm strainer. Livers were dissociated in HBSS supplemented with 0.5 mM EDTA with the GentleMACS Dissociator (Miltenyi) followed by digestion with 0.05% Collagenase Type IV and 10 U/ml DNase I in RPMI 1640 at 37°C. After straining through a 70 μm strainer, hepatocytes were separated from mononuclear cells by brief centrifugation followed by washing with RPMI 1640. Liver debris removal was performed using a 40/80% Percoll overlay and density centrifugation followed by washing with complete medium. Cell counting for brain, liver, muscle, meninges, skin, and lung was performed manually under a microscope with trypan blue. Cell counting for blood, spleen, lymph node, bone marrow was performed with an automated counter using trypan blue (with TC-20, Biorad) or propidium iodide (with Cellometer, Nelexcom).

### Surface staining, intracellular cytokine staining (ICS), and flow cytometry

Prior to surface staining, cells were incubated for 10 minutes at 4°C with 5 μg/ml anti-CD16/CD32 to block Fc receptors, or, for staining panels including tetramers, cells were incubated for 1 hour in the presence of tetramers and 5 μg/ml anti-CD16/CD32. For large spectral panels, viability staining was performed for 20 minutes at 4°C as an independent step, prior to surface staining for 30 minutes at 4°C with fluorescently conjugated antibodies diluted in Brilliant Stain Buffer (BD) and FACS Buffer (PBS supplemented with 0.05% sodium azide, 0.1% BSA, and 0.1 U/ml DNase I). For smaller panels, viability staining was performed simultaneously with surface staining for 30 minutes at 4°C with fluorescently conjugated antibodies diluted in PBS. For intracellular cytokine staining (ICS), cells were fixed and permeabilized (BD Cytofix/Cytoperm), then stained intracellularly for cytokine production with fluorescently conjugated antibodies diluted in permeabilization buffer for 20 minutes at 4°C. For intranuclear staining, cells were fixed and permeabilized (Tonbo Foxp3 / Transcription Factor Staining Kit), then stained intranuclearly with fluorescently conjugated antibodies diluted in permeabilization buffer overnight at 4°C. Samples were acquired on an Aurora spectral flow cytometer (Cytek) with unmixing performed in SpectroFlo (Cytek) or an LSR Fortessa (BD) with compensation applied in FACSDiva (BD). Data were analyzed with FlowJo software v10 (BD) or OMIQ (Dotmatics). Tetramer expression was not included as a parameter for generation of UMAPs or for unsupervised clustering. Unsupervised clustering of CD8^+^ and P1A_43-51_/H-2D^b^-tetramer^+^ T cell populations was performed in OMIQ using FlowSOM with the Euclidean distance metric and elbow metaclustering.

Analysis of polyfunctional CD8^+^ T cell responses was performed via a Boolean analysis of IFNγ^+^, TNF^+^, IL-2^+^, and in some cases CD107a^+^ and Granzyme B^+^ events in the CD3^+^CD8^+^ gate using FlowJo. Pestle and SPICE (Vaccine Research Centre, NIH, Bethesda) software were used to generate graphical representations of average proportions of T cells expressing combinations of cytokines^67^. The antibodies and tetramers used are shown in the Supplemental Data Table 3A. Gating strategies used for flow cytometry are shown in Supplemental Data 4.

### Peptide restimulation for evaluation of antigen-specific T cells in blood and spleen

For the detection of antigen-specific T cells after vaccination, freshly isolated PBMCs or splenocytes were stimulated with peptides at a concentration of 4 μg/ml per peptide (Genscript). For evaluation of responses against TAAs, ≥60% purity peptides corresponding to top predicted minimal epitopes were pooled for each gene (see Supplemental Data Table 2), and cells were stimulated with pools corresponding to vaccination group; DPY-vaccinated controls were stimulated with all minimal epitopes for all TAAs simultaneously. For immunogenicity screening of individual TAA epitopes, mice were vaccinated according to the prophylactic vaccination schedule, a splenocytes were isolated 2 weeks post-boost. For evaluation of responses against P1A, a pool of overlapping 15-mer peptides spanning the entire P1A protein sequence, each at 4 μg/ml and ≥75% purity, was used for stimulation. Stimulations occurred in the presence of DNase I (20µg/ml, Roche) and costimulatory anti-CD28 (2 µg/ml, Biolegend) at 37°C for 5 hours, adding brefeldin A (10 µg/ml, BioLegend) for the last 4 hours to allow accumulation of intracellular cytokines prior to ICS.

### Co-culture stimulation assay with *ex vivo* expanded P1A-specific CD8^+^ T cells

P1A-specific CD8^+^ T cells were expanded *ex vivo* from splenocytes of mice primed IM with 1×10^8^ IU ChAdOx1-P1A and boosted 28 days later IM with 1×10^7^ PFU MVA-P1A. Splenocytes from vaccinated mice were cultured in the presence of irradiated, syngeneic splenocytes from naïve mice which had been pre-pulsed with 4 μg/ml of P1A_43-51_ peptide (Peptides and Elephants) (i.e., antigen-presenting cells, APCs) to expand P1A_43-51_-specific CD8^+^ T cells in T cell media. T cells were restimulated every 9 days with peptide-pulsed APCs for a total of three simulations prior to use in assays. For the co-culture assay, expanded P1A_43-51_-specific CD8^+^ T cells were cultured at a 4:1 ratio with tumor cell targets for 14 hours at 37°C, adding 10 µg/ml brefeldin A for the last 12 hours to allow accumulation of intracellular cytokines, prior to ICS. Some tumor cells were pre-treated with 100 U/ml of IFNγ (Peprotech) for 48 hours prior to use in the assay, and some tumor cells were pre-pulsed with 4 μg/ml of P1A_43-51_ peptide.

### Restimulation of brain and blood T cells for polyfunctionality analysis

For the evaluation of T cell activation, freshly isolated PBMCs or immune cells from whole brains were stimulated with a pool of overlapping 15-mer peptides spanning the entire P1A protein sequence, each at 4 μg/ml and ≥75% purity. Stimulations occurred in the presence of DNase I (20 µg/ml, Roche), costimulatory anti-CD28 (2 µg/ml, Biolegend), and fluorescently conjugated anti-CD107a antibodies (1:1600, Biolegend) at 37°C for 16 hours, adding brefeldin A (10 µg/ml, BioLegend) for the last 12 hours to allow accumulation of intracellular cytokines prior to ICS.

### RMA-S MHC-I binding assay

RMA-S cells were obtained as a generous gift from Dr. David Margulies at NIH. RMA-S cells were cultured in the presence of human β2-microglobulin (5 μg/ml, Sigma) and increasing concentrations of peptides (≥85% purity, Genscript) for 16 hours. Cells were then stained simultaneously with fluorescently conjugated anti-H-2D^b^ and anti-H-2K^b^ antibodies prior to flow cytometry.

### Generation of SB28.Gpr149.KO cell line

TrueGuide Synthetic gRNA targeting Gpr149 (AAGUCAUAAUUCUACGGAGA, ThermoFisher), TrueCut Cas9 Protein v2 (ThermoFisher), and Lipofectamine CRISPRMAX (ThermoFisher) were combined in Opti-MEM (Gibco), then added to SB28 cells for 24 hours. Single cells were plated into 96-well plates by flow-assisted cell sorting (BD FACS Aria) to establish clones. To screen for complete Gpr149 knockout, DNA was isolated from each clone, the genetic region flanking the cut site was amplified via PCR, and the PCR product was purified via agarose gel electrophoresis prior to Sanger Sequencing. Knockout efficiency was measured with the SeqScreener Gene Edit Confirmation App (ThermoFisher). One monoclonal line with complete Gpr149 knockout was selected for use in all experiments.

### Generation of transgenic DC2.4 cell lines

DC2.4 cells were obtained as a kind gift from Dr. Jeffrey Schlom at NIH. DC2.4 lines expressing each TAA were created via lentiviral transduction. The sequences for Gpr149, Cdh6, Shox2, Igf2bp1, or the 11 Fbn2 minigenes (including spacer sequences) were PCR amplified from the same entry vector plasmids used to create the corresponding ChAdOx1/MVA viral vectors, then inserted into pLVX-TetOne-Puro lentiviral plasmids using InFusion cloning (Takara). Plasmids were verified via Sanger Sequencing before being used for co-transfection of HEK293T-LentiX cells alongside psPAX2 and pMD2.G with X-tremeGENE 9 Transfection Reagent (Sigma). Viral supernatants were collected and used to transduce DC2.4 cells, which were then selected for transgene expression with Puromycin (Gibco). Post-selection, single cells were isolated via FACS to establish monoclonal lines. TAA expression on monoclonal DC2.4 lines was verified via RT-qPCR. Clones expressing each TAA were selected for use in assays. DC2.4 cells were pre-treated for 48 hours with doxycycline (0.1 μg/ml, Sigma) to drive expression of the relevant TAA transgene before use in assays.

### DC2.4 co-culture assay

Mice were primed IM with 5×10^6 IU with a given ChAdOx1-TAA vaccine and boosted 28 days later with 1×10^6 IU of the same ChAdOx1-TAA vaccine IM. Splenocytes were isolated 2 weeks post-boost. To expand TAA-specific CD8^+^ T cells, splenocytes pooled from 3-5 vaccinated mice were co-cultured with irradiated DC2.4 lines expressing the relevant TAA transgene in the presence of human recombinant IL-2 (20 U/ml, Roche). Splenocytes from mice vaccinated with ChAdOx1-Combi were separately stimulated with either DC2.4.Shox2 or DC2.4.Igf2bp1. After 9 days, T cells were restimulated with the same cognate irradiated DC2.4-TAA cells or WT DC2.4 cells for 12 hours in the presence of anti-CD28 (2 µg/ml) and DNase (20 µg/ml), then for an additional 10 hours in the presence of Brefeldin A (10 µg/ml). IFNγ secretion was measured by ICS and flow cytometry.

### Stimulation assay of tetramer-labeled T cells

T cells were sorted from the spleens of ChAdOx1/MVA-Gpr149 vaccinated mice via FACS after staining with Gpr149_128-135_/H-2K^b^ and Gpr149_447-455_/H-2D^b^ tetramers and surface staining antibodies. Control CD3^+^CD8^+^ T cells were negative for both tetramers. For the data shown in Fig. 2H, T cells were FACS-isolated from the pooled splenocytes of tumor-naïve mice (*n* = 3) vaccinated according to the prophylactic ChAdOx1/MVA-Gpr149 six weeks post-boost. For the data shown in Supplementary Fig. 5G, T cells were sorted individually from splenocytes of individual tumor-naïve mice vaccinated according to the prophylactic ChAdOx1/MVA-Gpr149 two weeks post-boost. T cells were rested overnight at 37°C in T cell media supplemented with 20 U/ml IL-2 prior to use in the assay. Some target SB28 cells or SB28.Gpr149.KO cells were treated with rm-IFNγ (100 U/ml, Peprotech) for 48 hours prior to use in assay. 25,000 T cells were co-cultured with 50,000 tumor cells for 5 hours in T cell media in round-bottom 96-well plates in the presence of IL-2 (20U/ml), DNase (20 μg/ml, Roche), anti-CD28 (2 μg/ml, Biolegend), and CD107a-PE-Cy7 (1D4B, 125 ng/ml, BD). Brefeldin A (5 μg/ml, Biolegend) was added for the last four hours. ICS and flow cytometry was then performed as described above.

### Real-time monitoring of cell growth

Target SB28 cells or SB28.Gpr149.KO cells were treated with rm-IFNγ (100 U/ml, Peprotech) for 48 hours prior to harvest. Upon harvesting, cells for some test conditions were pulsed with Gpr149 peptides (4 μg/ml, 85% purity, Genscript) for 2 hours at 37°C, then washed thoroughly with PBS. Non-pulsed cells were handled identically. 2500 cells were plated in 100 μl T cell media supplemented with IL-2 (20 U/ml) in 16-well E-Plates (Agilent) and electrical impedance was measured every 15 minutes using an XCELLigence Real-Time Cell Analysis MP instrument (Agilent). A total of 25,000, 5000, or 1250 T cells in a volume of 100 μl T cell media were added 24 hours after initial plating of tumor cells. T cells were derived from the splenocytes of tumor-naïve mice vaccinated according to the prophylactic ChAdOx1/MVA-Gpr149 regimen two weeks post-boost. T cells were FACS-isolated after staining with anti-CD3 (17A2, Biolegend), anti-CD8 (53-6.7, Biolegend), and Gpr149_128-135_/H-2K^b^- and Gpr149_447-455_/H-2D^b^-tetramers. Tetramer-negative control T cells were negative for both tetramers. T cells were rested overnight at 37°C in T cell media supplemented with 20 U/ml IL-2 prior to use in the assay. ΔCell Index was calculated by the XCELLigence Real-Time Cell Analysis software (Agilent) from the time point directly prior to the addition of T cells.

### *In vivo* lysis assay

Splenocytes from naïve C57BL/6 mice were harvested and split three ways prior to pulsing for 2 hours with Gpr149_128-135_ or Gpr149_447-455_ peptides (10 μg/ml, Genscript). Cells were then labeled with 1 μM, 0.33 μM, or 0.11 μM CFSE (ThermoFisher) (CFSE^hi^: Gpr149_128-135_ pulsed, CFSE^mid^: non-pulsed, CFSE^lo^: Gpr149_447-455_ pulsed, respectively), mixed at a 1:1:1 ratio in PBS, and 2×10^6^ cells/condition were infused IV into vaccinated or naïve recipients. Vaccinated recipients were treated with ChAdOx1/MVA-Gpr149 according to the prophylactic dosing scheme prior to the infusion of CFSE-labeled cells 6 weeks post-boost. 72 hours post-infusion, spleens were harvested from recipients and run on a flow cytometer without additional staining. Percent lysis was calculated as (#Non-pulsed - #Pep-Pulsed)/#Non-Pulsed x 100.

### FTY720 treatment

Mice were injected IP daily with 1 mg/kg FTY720 in 100 μl, beginning two days prior to tumor challenge and extending through the study duration. FTY720 (Cayman Chemical) was reconstituted in DMSO and suspended in 10% DMSO, 40% polyethylene glycol 300, 5% Tween-80, and 45% Saline. Controls received daily injections of vehicle only.

### Intracranial adoptive transfer of T cells

Donor mice were generated via the prophylactic ChAdOx1/MVA-P1A dosing scheme and implanted with SB28.P1A tumors intracranially. Immune cells were isolated from blood and brains of perfused mice on 21 days post-tumor implantation followed by surface staining with fluorescently conjugated antibodies and tetramers as described above (Supplemental Data 4). T cell populations were isolated via FACS on a FACSAria sorter (BD) and washed prior to suspension in HBSS for intracranial implantation. Naïve recipient mice were stereotactically implanted with 5,000 T cells in 2 μl using the same experimental procedures as described for tumor implantations above. One week later, T cell recipient mice were implanted with 1,000 SB28.P1A tumor cells through the same Burr hole according to the procedures described above.

### Definition of T cell subsets

CD8^+^ T cells were distinguished by expression of CD45^hi^Lin^-^CD3^+^CD8^+^ by flow cytometry. CD8αβ^+^ T cells were identified as CD45^hi^CD11b^-^CD3^+^TCRβ^+^5-OP-RU/MR-1-tetramer^-^PBS-57 /CD1d-tetramer^-^CD8^+^CD4^-^. Resident memory T cells (TRMs) were CD44^hi^CD62L^-^CD8^+^ and CD103^+^CD69^+^, CD103^-^CD69^+^, or Runx3^+^CD49a^+^CD39^+^ T cells localized within tissues, as demonstrated by perfusion prior to tissue collection and immune cell isolation. CD44^hi^CD62L^-^CD8^+^ were referred to as double-positive (DP) cells when CD103^+^CD69^+^, as single-positive (SP) cells when CD103^-^CD69^+^, and as double-negative (DN) cells when CD103^-^CD69^-^. Short-lived effector cells (SLECs) were defined as CD8^+^CD44^hi^CD62L^-^KLRG1^+^CD127^-^. Terminally exhausted (T_TEX_) cells were defined as CD8^+^PD-1^+^TIM-3^+^. Progenitor exhausted cells (T_PEX_) were defined as CD8^+^PD-1^+^TCF-1^+^. P1A_43-51_-specific T cells were defined as P1A_43-51_/H-2D^b^-tetramer^+^CD3^+^CD8^+^. Gpr149_128-135_-specific T cells were defined as Gpr149_128-135_/H-2K^b^-tetramer^+^CD3^+^CD8^+^. Gpr149_447-455_-specific T cells were defined as Gpr149_447-455_/H-2D^b^-tetramer^+^CD3^+^CD8^+^.

### Statistical analysis and figure preparation

Statistical analysis was performed using GraphPad PRISM (V10) except for EdgeR analysis, which was conducted with OMIQ on equally subsampled data, and for analysis of TAAs in humans, which was performed in Python. All significant statistics are shown on all graphs when calculated unless otherwise indicated. Non-significant statistics are not shown in most cases, though they are included in some graphs for emphasis. For datasets in which all performed comparisons were non-significant, indication of “All *n.s.*” is written in figure legends. Figures were prepared using OMIQ, FlowJo, Adobe Illustrator, GraphPad PRISM, SPICE, and Biorender. UMAP plots display equally subsampled data from the samples indicated. For histograms created in OMIQ, a smoothing factor of 8 was applied.

---

**Supplementary Figure 1.**
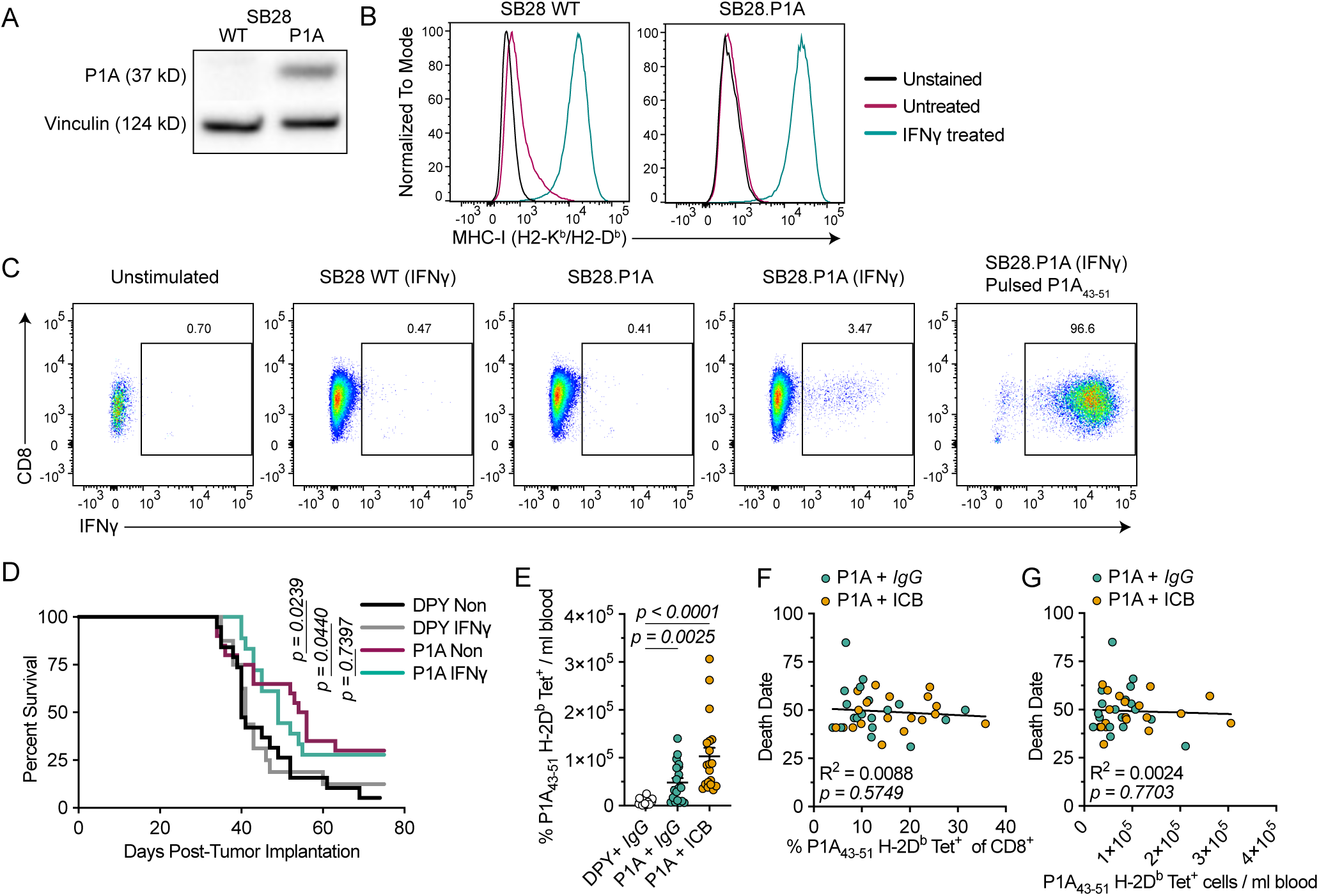
ChAdOx1/MVA-P1A vaccination is prophylactically and therapeutically effective against P1A-expressing SB28 tumors. (A) Western blot showing P1A expression in WT and P1A-transfected SB28 lines; vinculin housekeeping gene shown as loading control. (B) Representative histograms showing MHC-I expression on WT or P1A-expressing SB28 lines after 48 hours of IFNγ treatment. (C) Representative reactivity of P1A-specific T cells against WT, P1A-transfected, or P1A_43-51_-peptide pulsed SB28 cells, some of which were pre-treated with IFNγ. (D) Survival of DPY- or P1A-prophylactically vaccinated mice challenged with non-pretreated (Non) or IFNγ-pretreated SB28.P1A tumor cells (*n* = 16-20, pooled from 2 experiments). (E) Number of P1A_43-51_/H-2D^b^-tetramer^+^CD8^+^ T cells/ml of blood one-week post-boost (D28) for mice in Figure 1I-L. Data are presented as mean ± SEM (*n = 19*, pooled from 2 experiments). (F) Spearman rank correlation for date of death and proportion of P1A_43-51_/H-2D^b^-tetramer^+^CD8^+^ T cells in blood one-week post-boost (D28) for P1A + *IgG* (*aqua*) and P1A + ICB (*gold)* treated mice in Figure 1I-L (*n = 38*, pooled from 2 experiments). (G) Spearman rank correlation for date of death and number P1A_43-51_/H-2D^b^-tetramer^+^CD8^+^ T cells/ml of blood one-week post-boost (D28) for P1A + *IgG* (*aqua*) and P1A + ICB (*gold)* treated mice in Figure 1I-L (*n = 38*, pooled from 2 experiments). Statistical analysis was performed using log-rank test for (D), Kruskal-Wallis test with Dunn’s multiple comparisons for (E), and simple linear regression for (F-G).

**Supplementary Figure 2.**
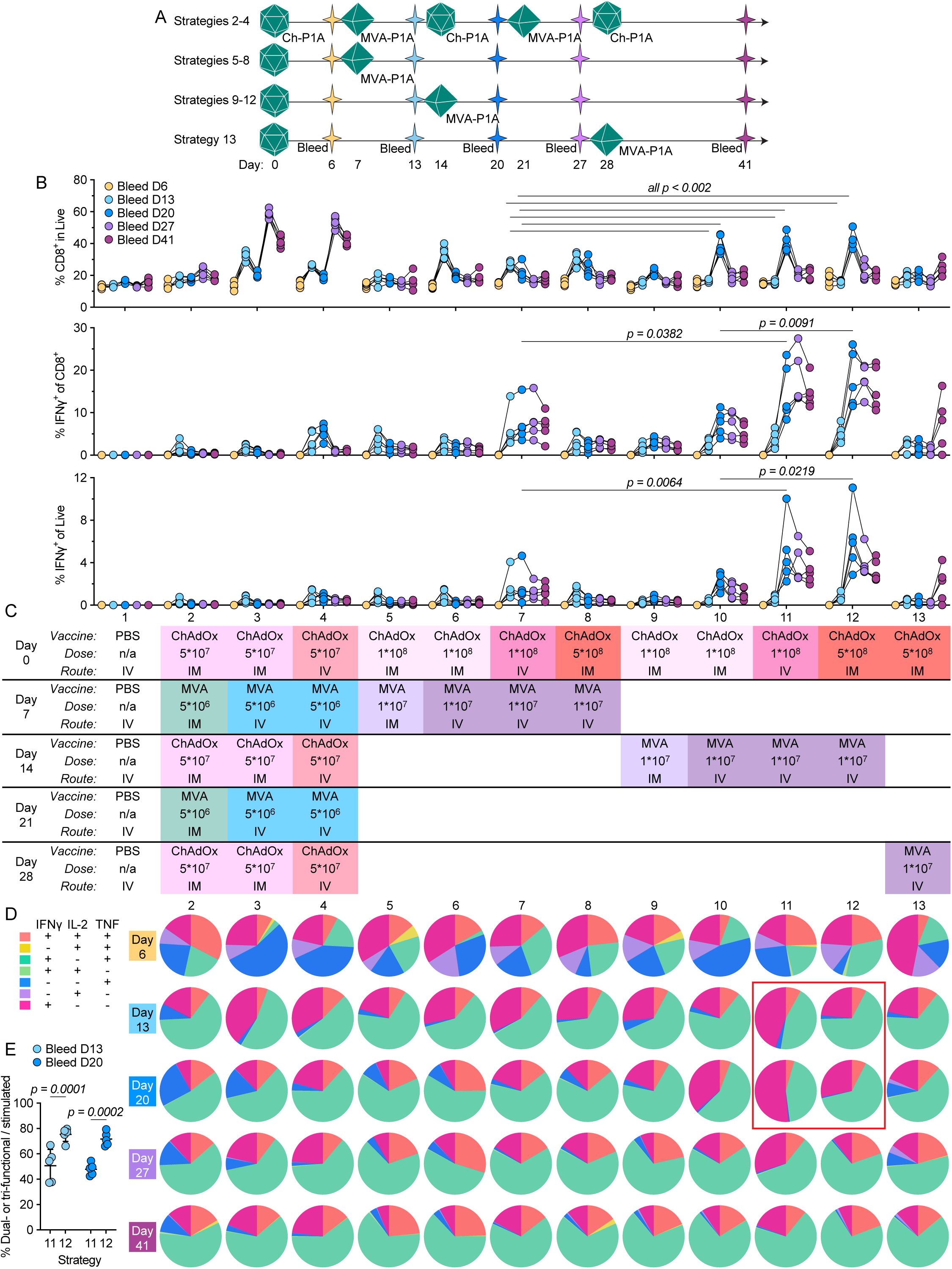
Optimization of vaccine dosing schedule and route for therapeutic treatment settings. (A) Schematic overview of vaccination and blood sampling timepoints for the experiment shown in (B-E). See (C) for dose and route details. (B) Frequency of CD8^+^ T cells among live cells (*top*), IFNγ^+^ cells among CD8^+^ T cells (*middle*), and IFNγ^+^ cells among live cells (*bottom*) from PBMCs after *ex vivo* stimulation with pooled P1A peptides at multiple sampling timepoints (*designated by color)* for mice treated via 13 different vaccination strategies detailed in (A) and (C) (*n = 5*). Only statistically significant comparisons between strategies 7, 10, 11, and 12 at blood sampling D13 and D20 timepoints are shown for space. (C) Table depicting vaccines administered by experimental day for 13 different vaccination strategies, listing vaccine (ChAdOx1-P1A or MVA-P1A), dose in IU (International Units) for ChAdOx1 or PFU (Plaque Forming Units) for MVA, and route of administration (IV, intravenous; IM, intramuscular). Empty white blocks for a given treatment day indicate that no treatment was given. (D) Polyfunctionality of cytokine-expressing CD8^+^ T cells for each vaccination strategy (*columns)* at each blood sampling timepoint (*rows*) (*n* = 5). (E) Proportion of cytokine-expressing CD8^+^ T cells expressing 2 or 3 cytokines at blood sampling days 13 and 20 for vaccination strategies 11 and 12; data are presented as mean ± SD (*n* = 5). Statistical analysis was performed using two-way repeated measures ANOVA with Fisher’s LSD multiple comparisons test for (B), (E).

**Supplementary Figure 3.**
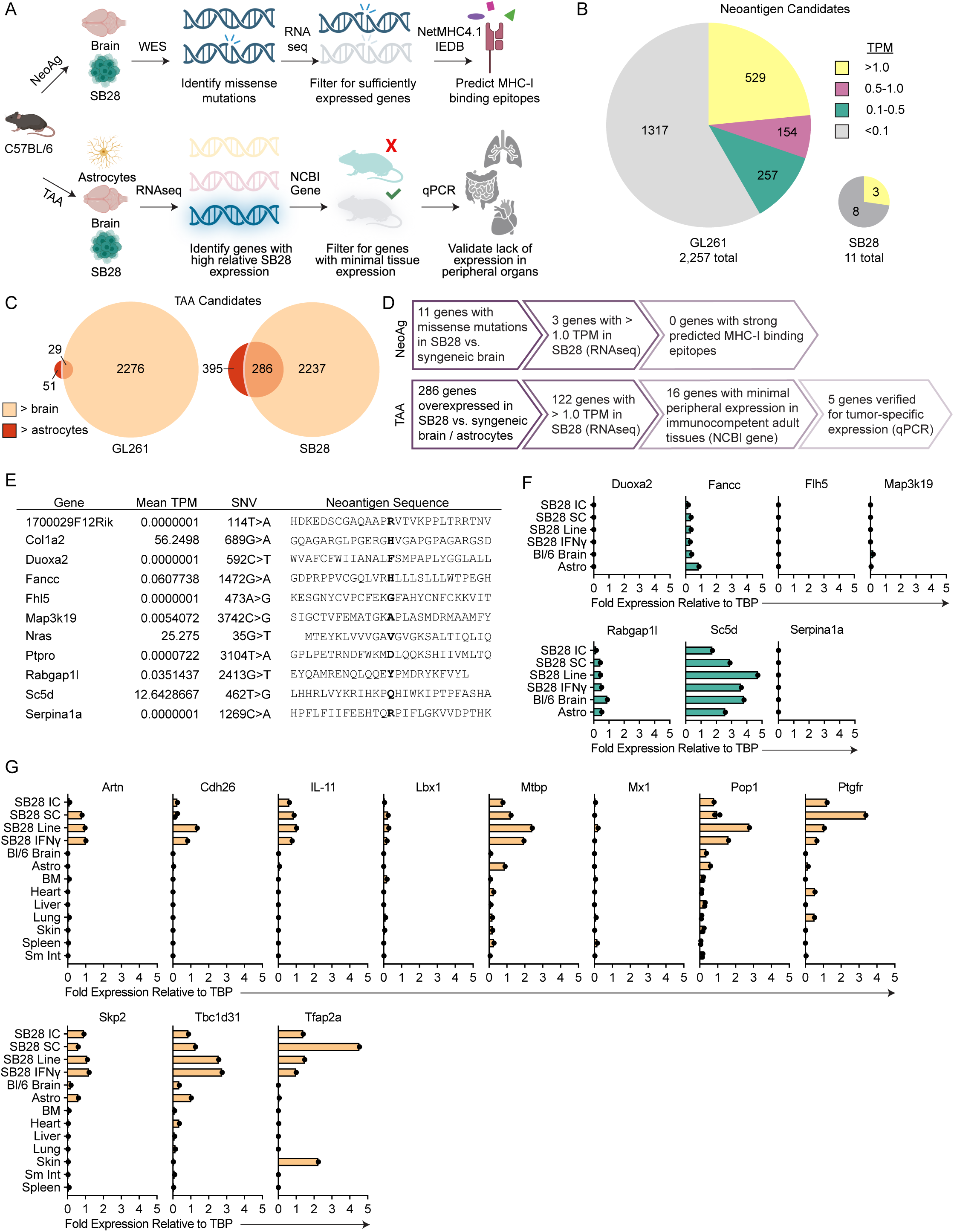
Identification of vaccine target candidates in the SB28 glioblastoma cell line. (A) Schematic depicting pipeline for identification of neoantigen (top) and TAA (bottom) candidates. (B) Numbers of neoantigen candidate genes in SB28 and GL261 cell lines. Candidates were determined as missense mutations in protein-coding genes based on comparative whole exome sequencing (WES). Relative expression level of each neoantigen candidate based on RNASeq data is summarized by color (Yellow > 1.0 Transcripts per Million (TPM), Pink 0.5-1.0 TPM, Aqua 0.1-0.5 TPM, Gray < 0.1 TPM). (C) Venn diagrams depicting relative numbers of overexpressed genes in SB28 and GL261 cell lines by comparative RNAseq, with area of circles proportional to the number of genes. Dark (left) circles represent genes with > 0.5 TPM in tumor cells and <0.5 TPM in astrocyte cells; light (right) circles represent genes with > 0.5 TPM in tumor cells and < 0.5 TPM in whole C57BL/6 brain; overlap represents genes with > 0.5 TPM in tumor cells and < 0.5 TPM in both astrocytes and whole brain. (D) Numbers of neoantigen (top) and TAA (bottom) candidates passing each stage of the pipeline for SB28. (E) Expression and sequences of all SB28 neoantigen candidates based on RNAseq of SB28 cell line. Values are mean expression of triplicates. TPM, transcripts per kilobase million. (F) Fold change in neoantigen candidate gene expression in SB28 samples or control tissues relative to a housekeeping gene (*Tbp*) as measured by RT-qPCR (*n =* 1). TBP, TATA-Box Binding Protein; IC, intracranial; SC, subcutaneous; IFNγ, cell line treated for 48 hours with IFNγ; Bl/6, C57BL/6; Astro, astrocytes. (G) Fold change in TAA candidate gene expression in SB28 samples or peripheral tissues relative to a housekeeping gene (*Tbp*) as measured by RT-qPCR. Data are presented as mean (*n =* 1-2). BM, bone marrow; Sm Int, small intestine; other abbreviations as in (F).

**Supplementary Figure 4.**
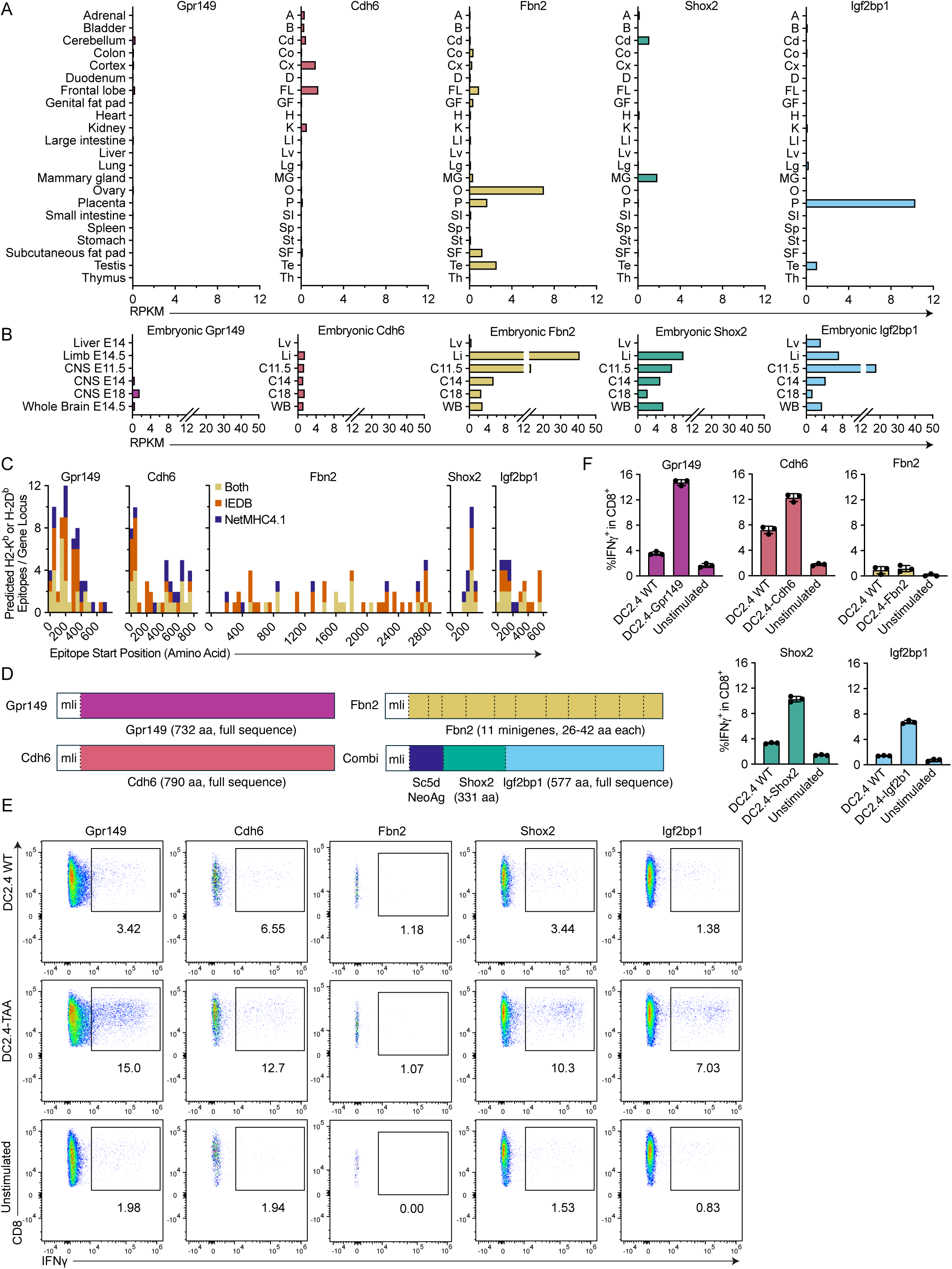
Validation of candidate SB28 tumor-associated antigens. (A) Transcriptomic expression of top 5 SB28 TAA candidate genes in various murine adult tissues in the Mouse ENCODE transcriptome dataset from NCBI Gene. RPKM, reads per kilobase million. (B) Transcriptomic expression of selected genes in various murine embryonic tissues from the Mouse ENCODE transcriptome dataset from NCBI Gene. Number following “E” designates age of embryo in days. (C) Histograms depicting number and distribution of unique 8-11mer epitopes beginning within a given range of amino acids (bin width = 50) for each gene predicted to bind H-2K^b^ or H-2D^b^ by the IEDB Consensus Algorithm (*orange*, Consensus Percentile Rank <1.5), NetMHC4.1 (*navy*, EL Rank < 0.5), or both (*yellow*). (D) Schematics representing the design of each vaccine. Dashed lines represent spacer amino acid sequences between gene segments (GGGPGGG). mIi, major histocompatibility complex class II-associated invariant chain. (E) Representative reactivity of *ex vivo* expanded CD8^+^ T cells after co-culture of T cells with WT or transgenic DC2.4 cells expressing cognate TAA gene inserts. FACS plots are pre-gated on Singlets/Live/CD3^+^CD8^+^. (F) Frequency of IFNγ-expressing CD8^+^ T cells after co-culture as in (E). Data are presented as mean ± SD (*n* = 3 technical replicates).

**Supplementary Figure 5.**
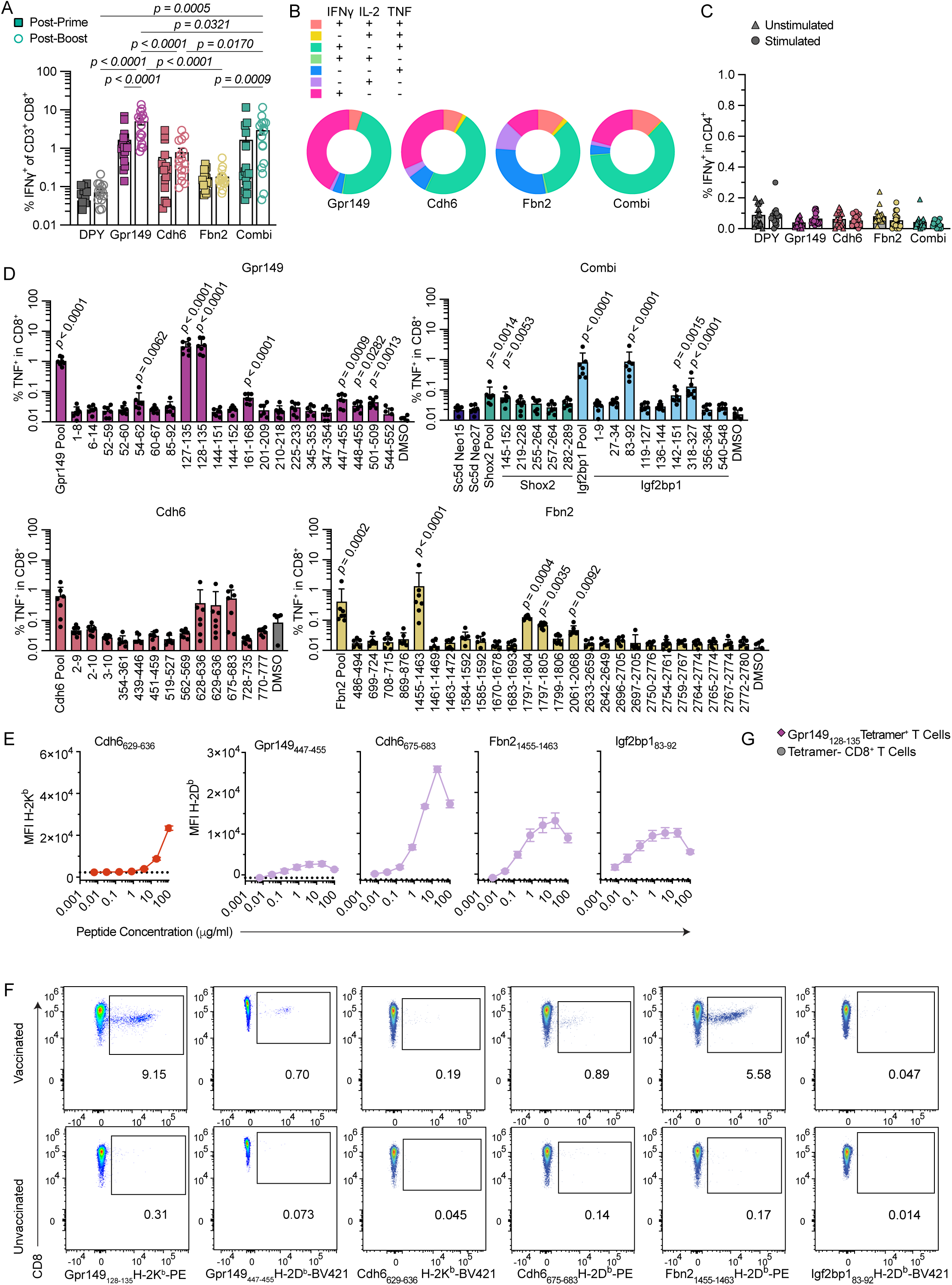
Immunogenicity of SB28-TAA vaccines. (A) Proportion of CD8^+^ T cells expressing IFNγ after *ex vivo* restimulation of PBMCs with relevant TAA peptide pools two weeks post-prime (*squares*) or two weeks post-boost (*circles*) (*n* = 16, pooled from 2 experiments). (B) Polyfunctionality of cytokine-expressing CD8^+^ T cells for each vaccination condition two weeks post-boost (*n* = 16, pooled from 2 experiments). (C) Proportion of CD4^+^ T cells expressing IFNγ after *ex vivo* restimulation of PBMCs with relevant TAA peptide pools (*circles*) or DMSO (*triangles*, unstimulated controls) two weeks post-boost (*n* = 16, pooled from 2 experiments). All *n.s*. (D) Frequency of CD8^+^ T cells expressing TNF after *ex vivo* restimulation of splenocytes from TAA-vaccinated mice with minimal epitope peptides (*n* = 7, pooled from 2 experiments). (E) Median fluorescence intensity (MFI) of surface H-2D^b^ or H-2K^b^ on RMA-S cells after culture with the indicated peptide at increasing concentrations; dotted line indicates baseline. All peptides eliciting a statistically significant response in (D) were tested; data shown only for those with successful MHC-I stabilization (*n* = 3 technical replicates). (F) Representative flow cytometry plots pre-gated on CD3^+^CD8^+^ T cells depicting tetramer labeling of splenocytes from untreated mice or mice vaccinated with the relevant set of ChAdOx1/MVA TAA vaccines. (G) Proportion of IFNγ^+^CD107a^+^ cells among sorted Gpr149_128-135_-specific or nonspecific T cells from Gpr149-vaccinated mice after co-culture with SB28 cells (*n =* 4 mice). Statistics not shown for Gpr149 vs. nonspecific T cell comparisons. Data are presented as mean ± SEM for (A), (C), (G), mean for (B), and mean ± SD for (D-E). Statistical analysis was performed using two-way ANOVA with Tukey’s multiple comparisons test for (A), (G), two-way ANOVA with Sídǎk’s multiple comparisons test (Unstimulated vs. Stimulated) for (B), and Kruskal-Wallis test with Dunn’s multiple comparisons for each stimulation condition vs. DMSO controls for (D-E).

**Supplementary Figure 6.**
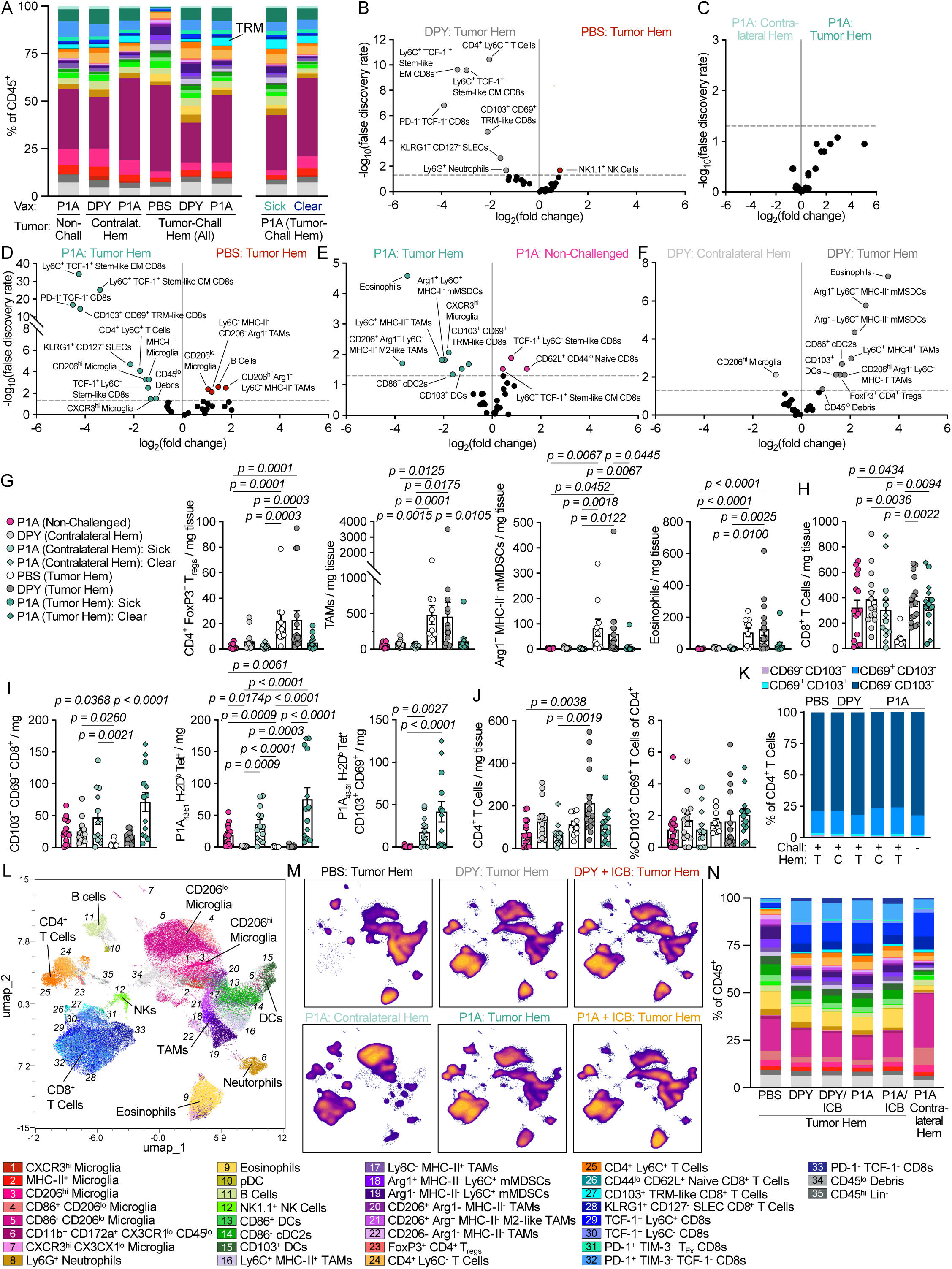
Prophylactic and therapeutic P1A-targeted vaccination remodels tumor immune subsets. For panels (A-K), mice were prophylactically vaccinated with ChAdOx1/MVA-P1A or - DPY, or were non-vaccinated (PBS-treated), then challenged with SB28.P1A tumors (except non-challenged group), as shown in the experimental scheme in Figure 3A. Brains and blood were collected 21 days post-tumor challenge for flow cytometry. (A) Proportion of CD45^+^ cells in brains corresponding to subsets in (Fig. 3B). Subsets are stacked ascending from bottom in the order listed in (Fig. 3B), except clusters 31 and 32 placed at bottom (*n =* 6-17, pooled from 3 experiments). (B-F) Volcano plots showing relative expression of immune subsets listed in Figure 3B for two-way comparisons between treatment groups designated at the top of each graph (*n =* 5-10). (G-J) Flow cytometry analysis of immune subsets in brains of mice, shown by number of cells / mg brain tissue or percent of parent gates, all utilizing the same color/group key as in (G). Datapoints from “tumor-cleared” mice (see Figure 3C) are designated by diamonds (*n* = 10-17, pooled from 3 experiments). (K) Proportion of CD4^+^ T cells in brains of mice expressing CD103 or CD69 for P1A-vaccinated mice (*n* = 10-17, pooled from 3 experiments). T, tumor-challenged hemisphere; C, contralateral hemisphere. For panels (L-N), mice bearing intracranial SB28.P1A tumors were therapeutically vaccinated with ChAdOx1/MVA-P1A or -DPY. Some mice also received anti-PD-1 + anti-CTLA-4 (ICB) as shown in the experimental scheme in Figure 3K. Brains were collected 28 days post-tumor challenge for flow cytometry. (L) UMAP of flow cytometry data from 54 brain samples. (M) Heatmaps showing distribution of immune subsets as in (A) by treatment group and brain hemisphere (Hem) (*n* = 4-12, pooled from 2 experiments, except for “PBS” and “P1A Contralateral Hem” groups). (N) Proportion of CD45^+^ cells in brains corresponding to subsets in (A). Subsets are stacked ascending from bottom in the order listed in (A), except clusters 34 and 35, which are placed at bottom. Data are presented as mean (*n =* 6-19, pooled from 2-3 experiments, except for “P1A Contralateral Hem” group). Data are presented as mean for (A), (K), (N), and mean ± SEM for (G-J). Statistical analysis was performed using edgeR for (B-F), and Kruskal-Wallis test with Dunn’s multiple comparisons for (G-J).

**Supplementary Figure 7.**
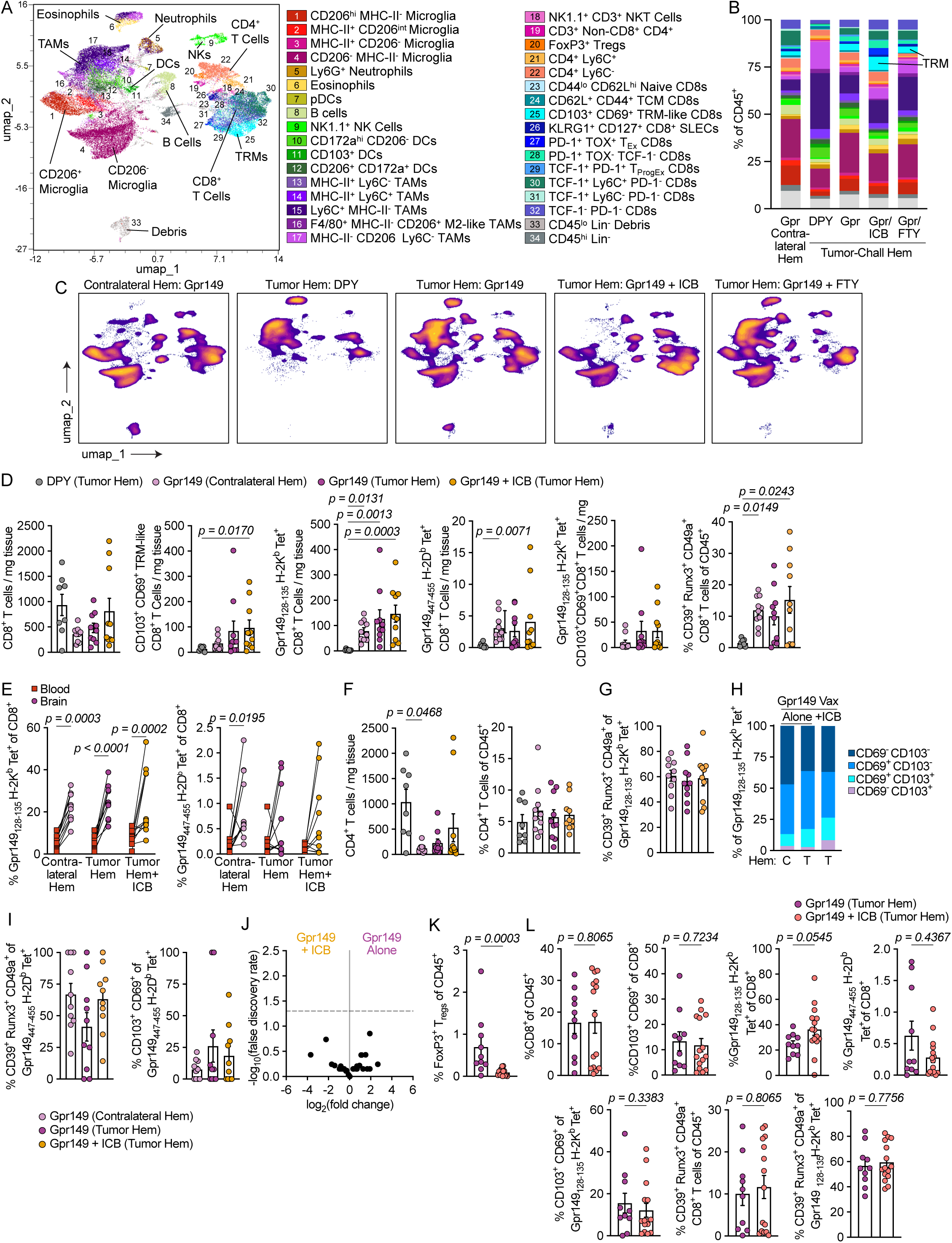
Prophylactic Gpr149-targeted vaccination remodels tumor immune subsets and induces high levels of CD103^+^CD69^+^ TRM-like CD8^+^ T cells. For all panels, mice were prophylactically vaccinated with ChAdOx1/MVA-Gpr149 or - DPY, then challenged with SB28 tumors. Some mice also received anti-PD-1 + anti-CTLA-4 (ICB) as shown in the experimental scheme in Figure 2A. Other mice also received FTY720 (FTY) daily as shown in the experimental scheme in Figure 7Q. Brains and blood were collected 21 days post-tumor challenge for flow cytometry. (A) UMAP of flow cytometry data from 23 brain samples. (B) Proportion of CD45^+^ cells in brains corresponding to subsets in (A). Subsets are stacked ascending from bottom in the order listed in (A), except clusters 33 and 34, which are placed at bottom (*n =* 8-15, pooled from 2 experiments). (C) Heatmaps showing distribution of immune subsets as in (A) by treatment group and brain hemisphere (Hem) (*n* = 3-5). (D) Flow cytometry analysis of immune cells in brains, shown by percent of parent gates or number of cells/mg brain tissue (*n* = 8-10, pooled from 2 experiments). (E) Proportion of CD8^+^ T cells binding Gpr149_128-135_/H-2K^b^-tetramers (*left*) or Gpr149_447-455_ H-2D^b^-tetramers (*right*) in brains or blood of Gpr149-vaccinated mice at D21 (*n* = 10, pooled from 2 experiments). (F-G) Flow cytometry analysis of immune cells in brains, shown by percent of parent gates or number of cells/mg brain tissue, using the group key shown above (D) (*n* = 10, pooled from 2 experiments). (H) Proportion of Gpr149_128-135_/H-2K^b^-tetramer^+^CD8^+^ T cells in brains expressing CD103 and/or CD69 for Gpr149-vaccinated mice (*n* = 10, pooled from 2 experiments). C, contralateral hemisphere; T, tumor-challenged hemisphere. (I) Flow cytometry analysis of immune cells in brains, shown by percent of parent gates, (*n* = 10, pooled from 2 experiments). (J) Volcano plot showing relative expression of immune subsets listed in (A) for two-way comparisons between treatment groups (*n =* 5). (K-L) Flow cytometry analysis of immune cells in brains, shown by percent of parent gates, utilizing the color/group key shown above (L) (*n* = 10-15, pooled from 2 experiments). Data are presented as mean for (B), (H) and mean ± SEM for (D), (F-G), (I), (K-L). Statistical analysis was performed using Kruskal-Wallis test with Dunn’s multiple comparisons for (D), (F-G), (I), RM two-way ANOVA with Sídǎk’s multiple comparisons test (Blood vs. Brain) for (E), edgeR for (J), and two-tailed Mann-Whitney test for (K-L).

**Supplementary Figure 8.**
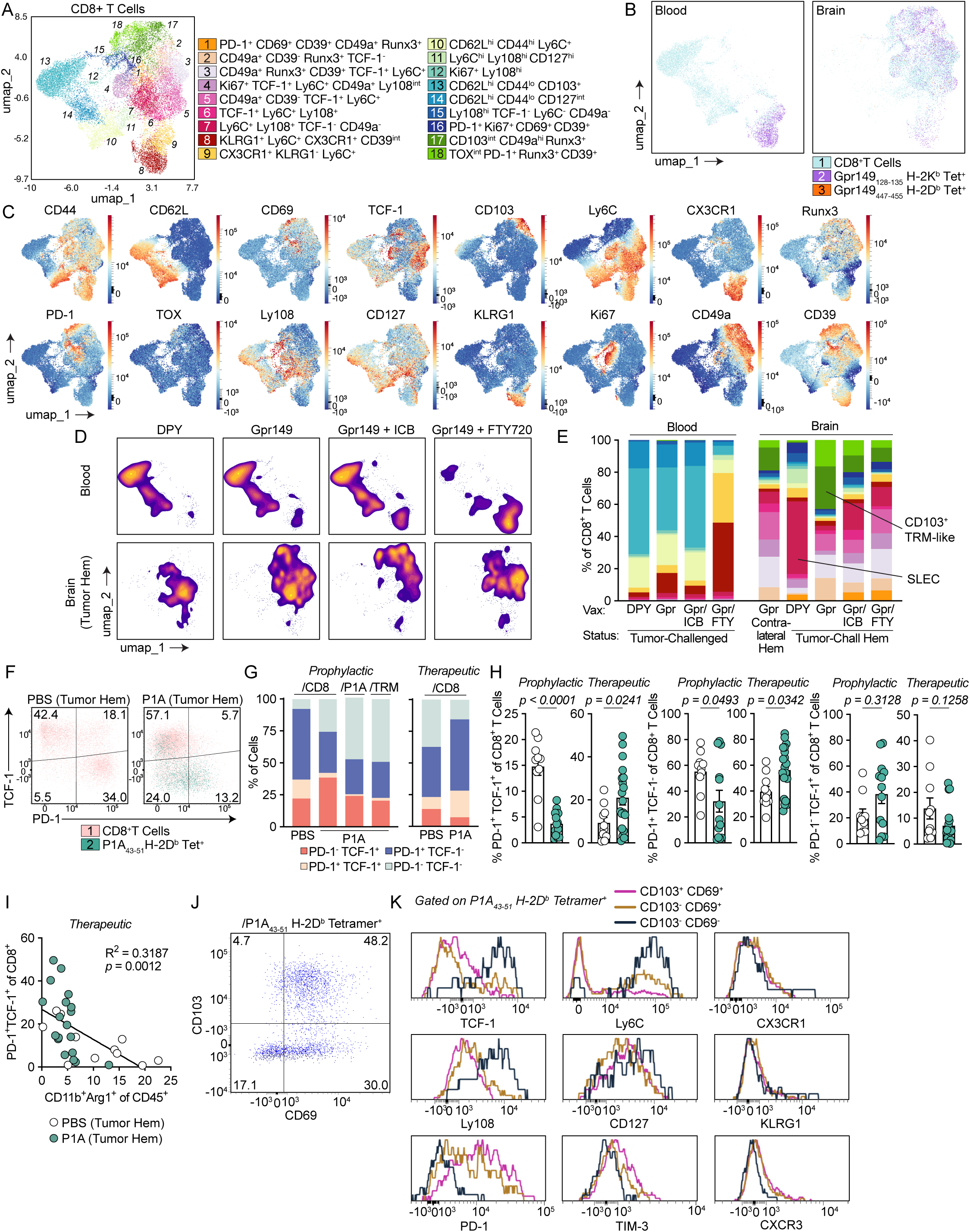
Vaccination induces TRM-like CD8^+^ T cell subsets in brain tumors with a distinct phenotypic signature. Panels (A-E) refer to flow cytometry data from brains and blood of Gpr149-prophylactically vaccinated mice as described in Supplementary Figure 7 and 7O. Panels (F-H and J-K) refer to flow cytometry data from brains of untreated (PBS) or P1A-prophylactically vaccinated mice as described in Figure 3A. Panels (G-I) additionally refer to flow cytometry data from brains of untreated (PBS) or P1A-therapeutically vaccinated mice as described in Supplementary Figure 6. (A) UMAP of CD8^+^ T cells from 23 brain and 18 blood samples. (B) Overlay of Gpr149 tetramer^+^ T cells atop CD8^+^ T cell UMAP for blood or brain (*n =* 18-23). (C) Heatmap overlay of phenotypic markers for UMAP in (A) (n = 41). (D) Heatmaps showing distribution of cells as in (A) by treatment group and tissue (*n* = 3-5). (E) Proportion of CD8^+^ cells in brains or blood corresponding to subsets listed in (A), stacked ascending from bottom (*n =* 5-10). (F) TCF-1 vs. PD-1 expression on CD8^+^ and P1A_43-51_/H-2D^b^-tetramer^+^ T cells from brains of untreated or P1A-vaccinated mice (concatenated from *n =* 5-10). (G) Proportion of CD8^+^, P1A_43-51_/H-2D^b^-tetramer^+^, or CD103^+^CD69^+^ T cells from brains of untreated or P1A-vaccinated mice expressing TCF-1 and PD-1 (*n* = 10-19, each pooled from 2-3 experiments). (H) Proportion of brain CD8^+^ T cells expressing phenotypic markers for untreated or P1A-vaccinated mice (n = 10-19, each pooled from 2-3 experiments). (I) Correlation between proportions of PD-1^+^TCF-1^+^ among CD8^+^ T cells and Arg1^+^CD11b^+^ among CD45^+^ cells in brains of PBS-treated or P1A-therapeutically vaccinated mice. (J) CD103 vs. CD69 expression on P1A_43-51_/H-2D^b^-tetramer^+^ T cells from brains of tumor-challenged P1A-vaccinated mice (concatenated from *n =* 10). (K) Histograms showing expression of phenotypic markers on brain P1A_43-51_/H-2D^b^-tetramer^+^ T cells by CD103 and CD69 expression from tumor-challenged P1A-vaccinated mice (concatenated from *n =* 10). Data are presented as mean for (E), (G), and mean ± SEM for (H). Statistical analysis was performed using two-tailed Mann-Whitney test for (H) and a simple linear regression for (I).

**Supplementary Figure 9.**
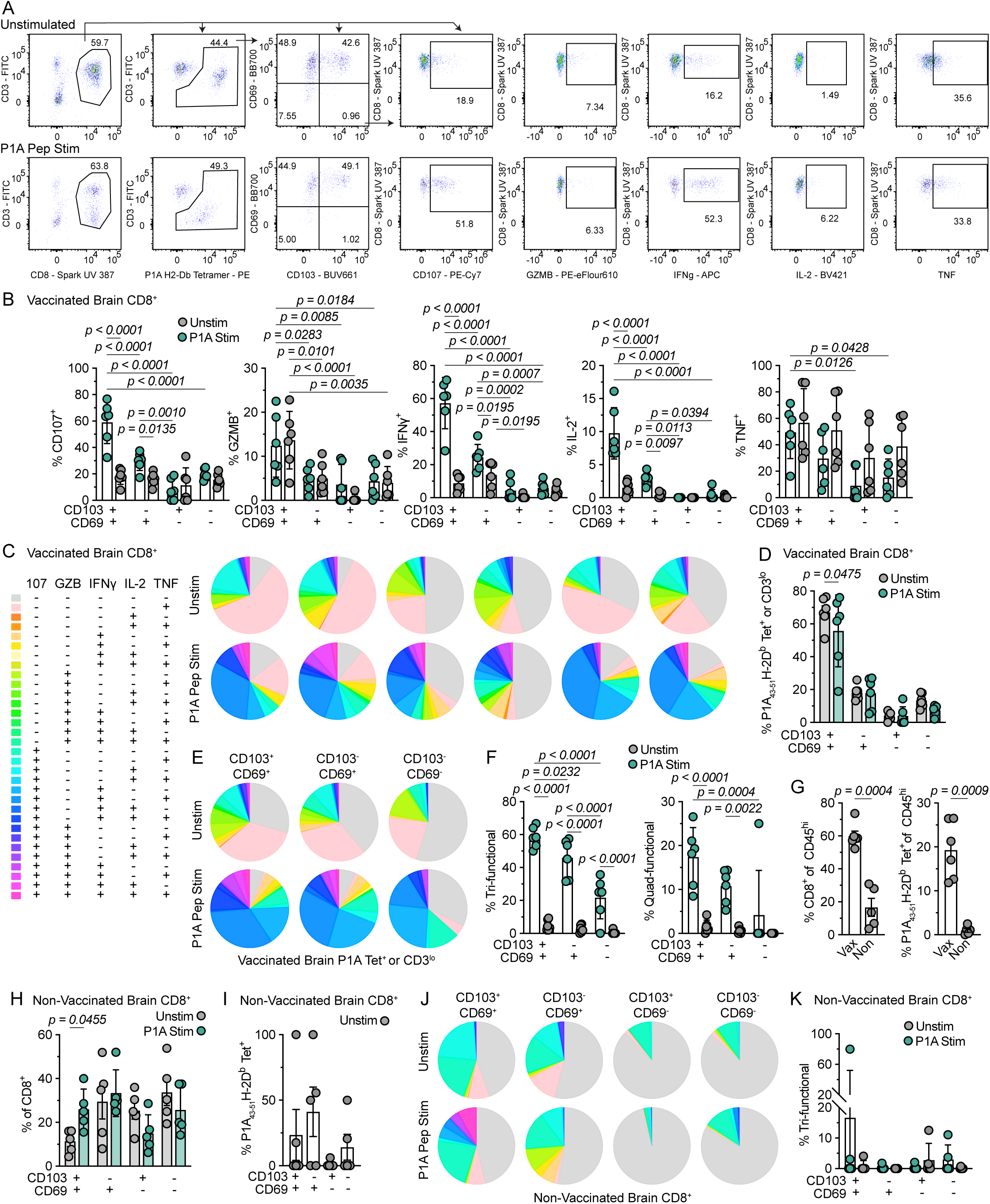
CD103^+^ CD69^+^ CD8^+^ T cells in brain tumors are highly polyfunctional. For all panels, mice were prophylactically vaccinated with ChAdOx1/MVA-P1A or were unvaccinated, then challenged with SB28.P1A tumors prior to isolation of immune cells from brains 21 days post-tumor challenge (as in Figure 3A) for *ex vivo* restimulation and flow cytometry. (A) Representative gating strategy for T cell phenotyping, displayed for a matched unstimulated vs. peptide-stimulated sample. (B) Monofunctionality of CD8^+^ T cells by CD103 and CD69 expression from brains of vaccinated mice, with or without stimulation (*n* = 6). (C) Polyfunctionality of all CD8^+^ T cells from brains of vaccinated mice, showing each matched sample individually with (*bottom*) or without (*top*) stimulation. (D) Proportion of CD8^+^ T cells from brains of vaccinated mice which are P1A_43-51_/H-2D^b^-tetramer^+^ or CD3^lo^ (as in second gate from left in (A)) by CD103 and CD69 expression, with or without stimulation (*n* = 6). Statistical comparisons between CD103/CD69 subsets not shown. (E) Polyfunctionality of P1A_43-51_/H-2D^b^-tetramer^+^ or CD3^lo^ CD8^+^ T cells (as in (I)) from brains of vaccinated mice by CD103 and CD69 expression (*columns)* with or without stimulation (*rows*), utilizing the same color key as in (C) (*n* = 6). (F) Proportion of P1A_43-51_/H-2D^b^-tetramer^+^ or CD3^lo^ CD8^+^ T cells (as in (I)) from brains of vaccinated mice simultaneously expressing at least 3 or 4 markers (as in E) by CD103 and CD69 expression, with or without stimulation (*n* = 6). (G) Proportion of CD8^+^ or P1A_43-51_/H-2D^b^-tetramer^+^ T cells among CD45^hi^ cells in brains of vaccinated or non-vaccinated mice without stimulation (*n* = 5-6). (H) Proportion of CD8^+^ T cells in brains of non-vaccinated mice expressing CD103 and/or CD69 with or without stimulation (*n =* 5). Statistical comparisons between CD103/CD69 subsets not shown. (I) Proportion of CD8^+^ T cells from brains of non-vaccinated mice binding P1A_43-51_/H-2D^b^-tetramers by CD103 and CD69 expression, without stimulation (*n =* 5). (J) Polyfunctionality of CD8^+^ T Cells from brains of non-vaccinated mice by CD103 and CD69 expression (*columns)* with or without stimulation (*rows*), utilizing the same color key as in (C) (*n* = 5). (K) Proportion of CD8^+^ T cells from brains of non-vaccinated mice simultaneously expressing at least 3 markers (as in J) by CD103 and CD69 expression, with or without stimulation (*n* = 5). Data are presented as mean ± SD for (B), (F), (K), and mean ± SEM for (D), (G-I), and mean for (E), (J). Statistical analysis was performed using two-way ANOVA with Tukey’s multiple comparisons test for (B), (D), (F), (H), (K), two-tailed Welch’s t test for (G), and Kruskal-Wallis test with Dunn’s multiple comparisons for (I).

**Supplementary Figure 10.**
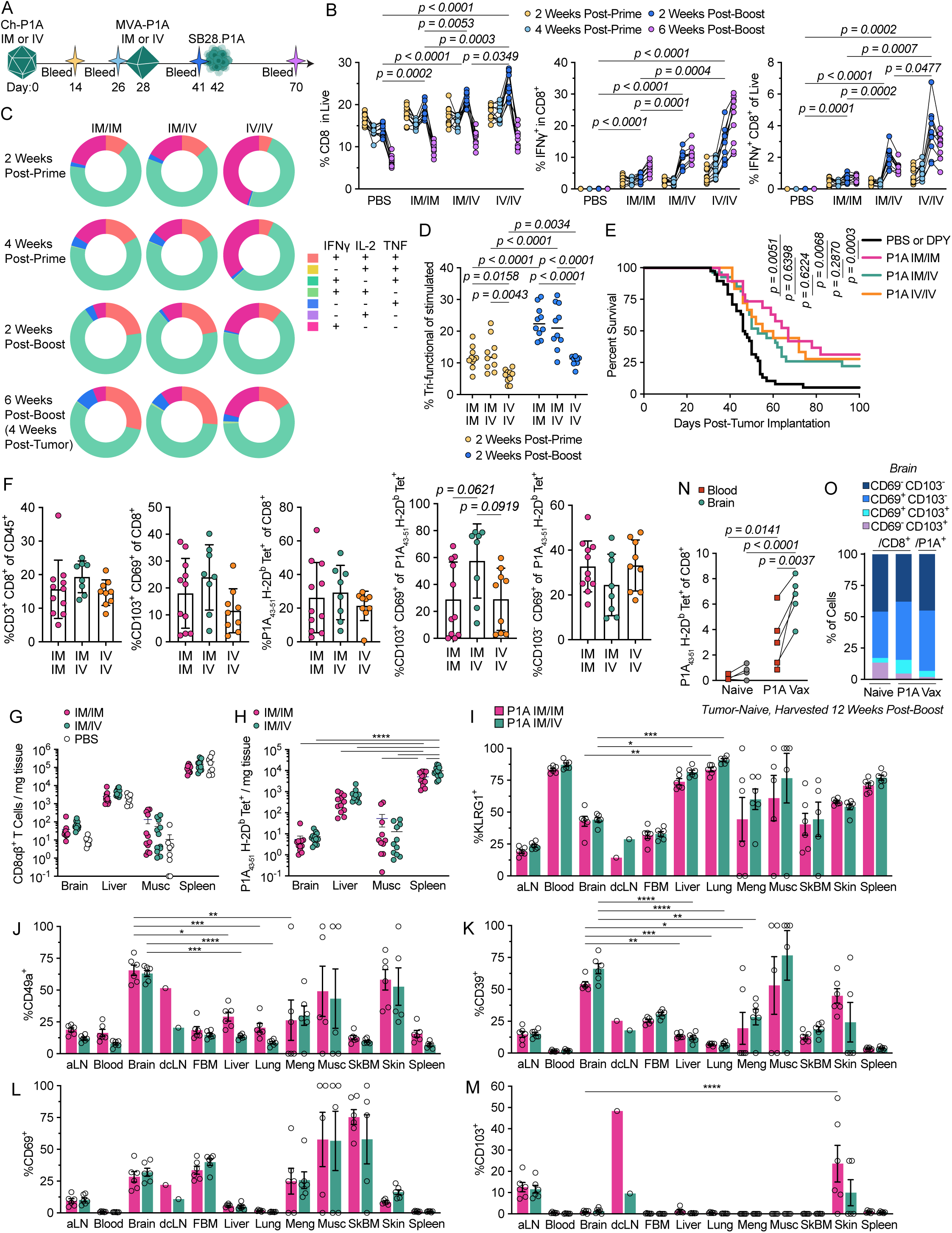
Impact of route of prophylactic vaccination on tissue-infiltrating CD8^+^ T cells. (A) Experimental scheme for (B-F): mice were prophylactically vaccinated with ChAdOx1/MVA-P1A via different routes of injection, or were non-vaccinated (PBS-treated), then challenged intracranially with SB28.P1A tumors. Mice used for (F) were bled only on D41, and brains were collected 21 days post-tumor challenge for flow cytometry. (B) Frequency of CD8^+^ T cells among live cells (*left*), IFNγ^+^ cells among CD8^+^ T cells (*middle*), and IFNγ^+^ CD8^+^ T cells among live cells (*right*) from PBMCs after *ex vivo* stimulation with P1A peptides at multiple sampling timepoints (*designated by color)* for mice treated via different vaccination strategies detailed (*n =* 9-10). (C) Polyfunctionality of cytokine-expressing CD8^+^ T cells for each vaccination strategy (*columns)* at each blood sampling timepoint (*rows*) (*n* = 9-10). (D) Proportion of cytokine-expressing CD8^+^ T cells expressing all 3 cytokines at blood sampling days 14 and 41 (*n* = 10). (E) Survival of prophylactically challenged mice (*n =* 18-38, pooled from 4 experiments). (F) Flow cytometry analysis of immune cell subsets in brains of prophylactically vaccinated mice, shown by percent of parent gates (*n* = 8-11, pooled from 3 experiments, all *n.s.*). For panels (G-M), mice were primed with ChAdOx1-P1A IM and boosted with MVA-P1A IM or IV 28 days later; controls received PBS (as in Fig. 6). Tissues were harvested 14 days post-boost for flow cytometry. Tissue abbreviations are as follows: aLN, axillary lymph nodes; dcLN, deep cervical lymph nodes; Meng, meninges; Musc, muscle; SkBM, skull bone marrow. (G) CD8αβ^+^ T cells/mg of tissue by tissue and route of vaccination (*n =* 8-12, pooled from 2 experiments). (H) P1A_43-51_/H-2D^b^-tetramer^+^ CD8αβ^+^ T cells/mg of tissue from vaccinated mice by tissue and route of vaccination (*n =* 12, pooled from 2 experiments). Statistical comparisons between vaccination strategies are shown only within a given tissue. (I-M) Proportion of P1A_43-51_/H-2D^b^-tetramer^+^CD8αβ^+^ T cells expressing phenotypic markers by tissue and vaccination strategy (*n =* 5-6). For (N-O), brains and blood of naïve or prophylactically P1A-vaccinated mice were harvested 12 weeks post-boost. (N) Proportion of P1A_43-51_/H-2D^b^-tetramer^+^ cells among CD8^+^ T cells in blood (*squares*) or brains (*circles*) (*n =* 4-5). (O) Proportion of CD8^+^ or P1A_43-51_/H-2D^b^-tetramer^+^ T cells in brains expressing CD103 or CD69 (*n* = 4-5). Data are presented with line at median for (D), mean ± SD for (F), mean ± SEM for (G-M), and mean for (O). Statistical analysis was performed using a mixed-effects model with Tukey’s multiple comparisons (by timepoint) for (B), two-way repeated measures ANOVA with Tukey’s multiple comparisons test for (D), log-rank test for (E), Kruskal-Wallis test with Dunn’s multiple comparisons for (F), two-way ANOVA with Tukey’s multiple comparisons for (H-M), and two-way ANOVA with Fisher’s LSD multiple comparisons test for (N). For (I-M), significant comparisons are shown only for brain vs. liver, lung, meninges, and skin within a given vaccination strategy, and brain IM vs. IV. \**p* < 0.05, \*\**p* < 0.01, \*\*\**p* < 0.001, \*\*\*\**p* < 0.0001.

**Supplementary Figure 11.**
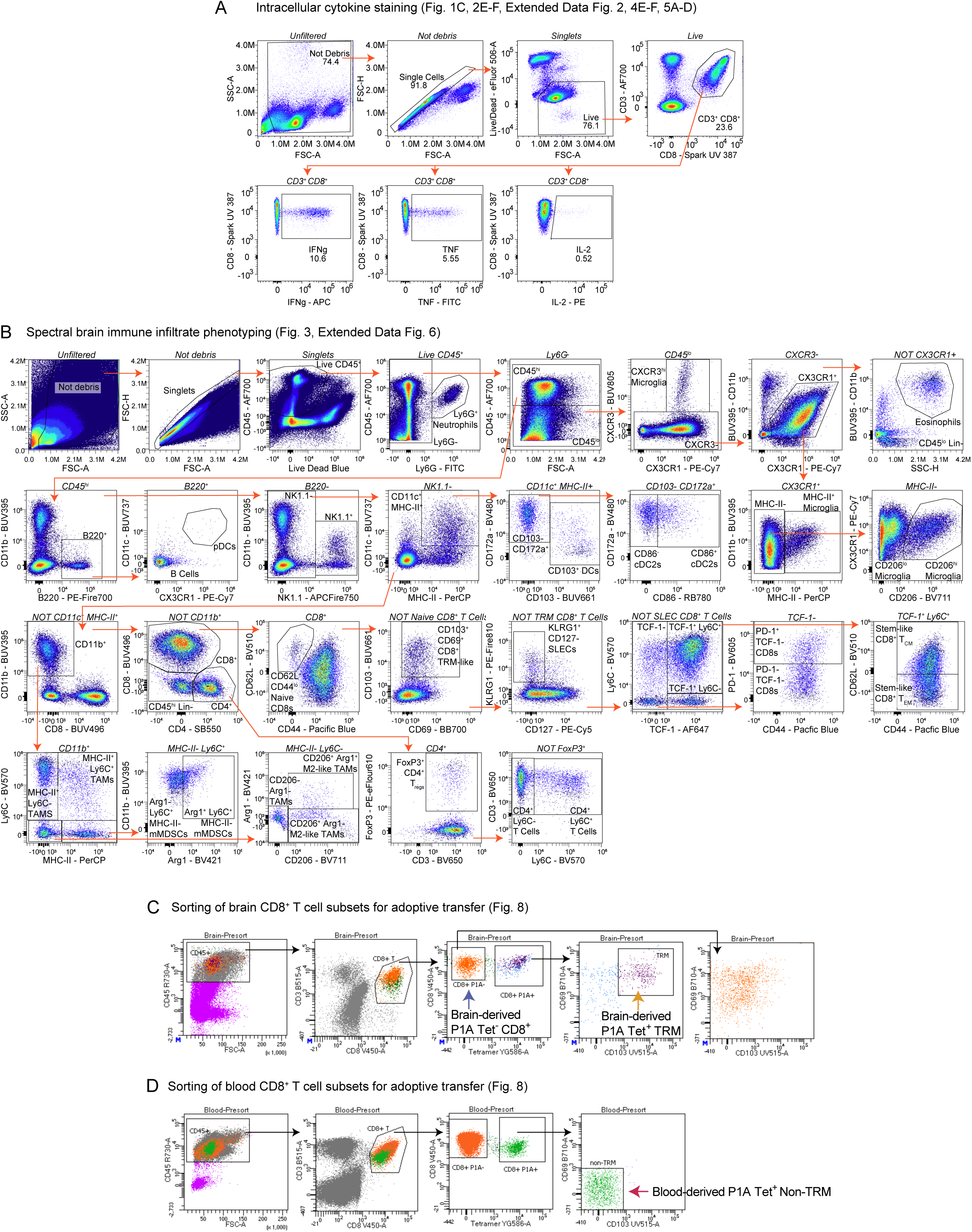
Representative flow cytometry and sorting gating strategies.

**Supplementary Data Table 1A.**
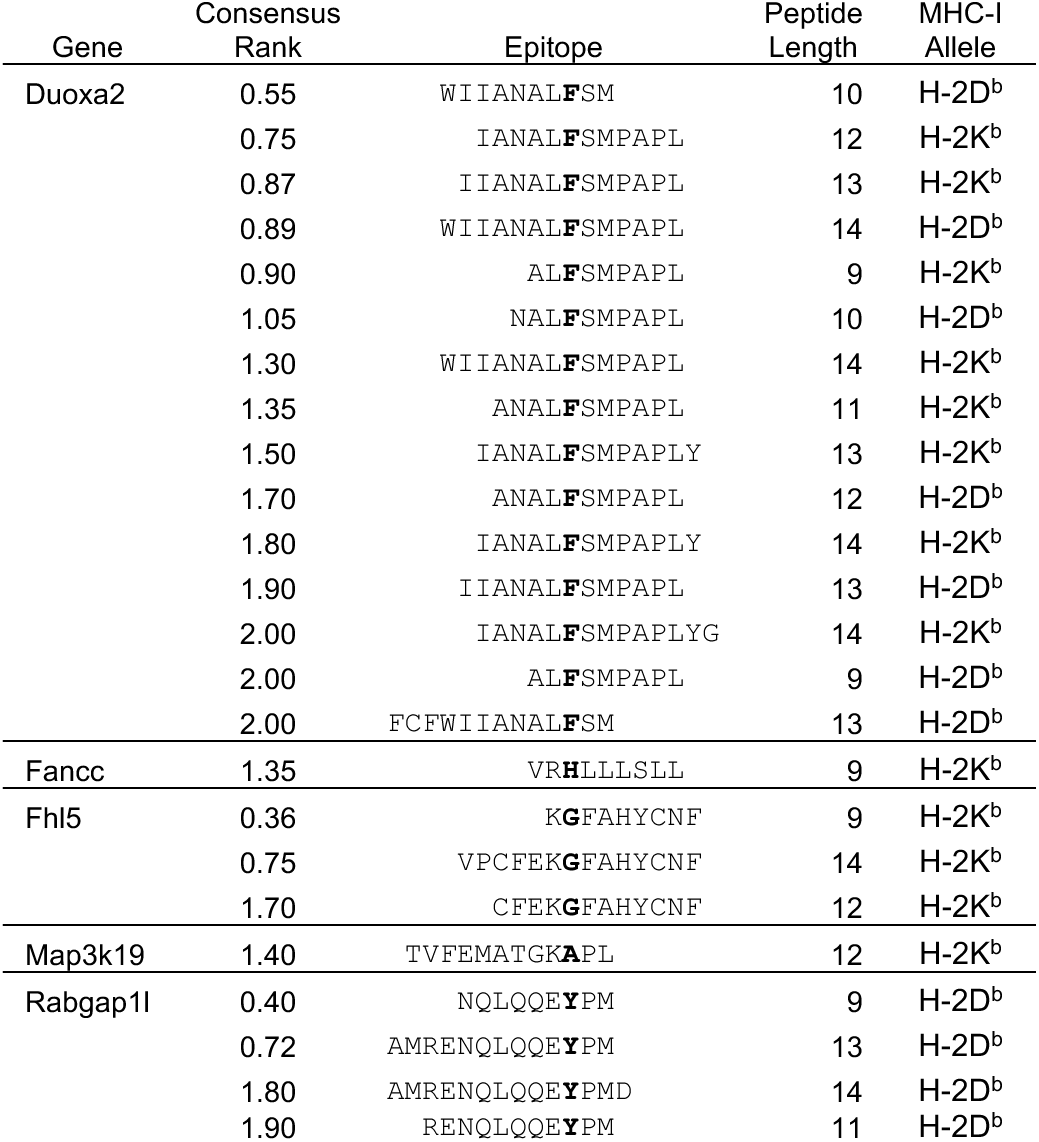

**Supplementary Data Table 1B.**
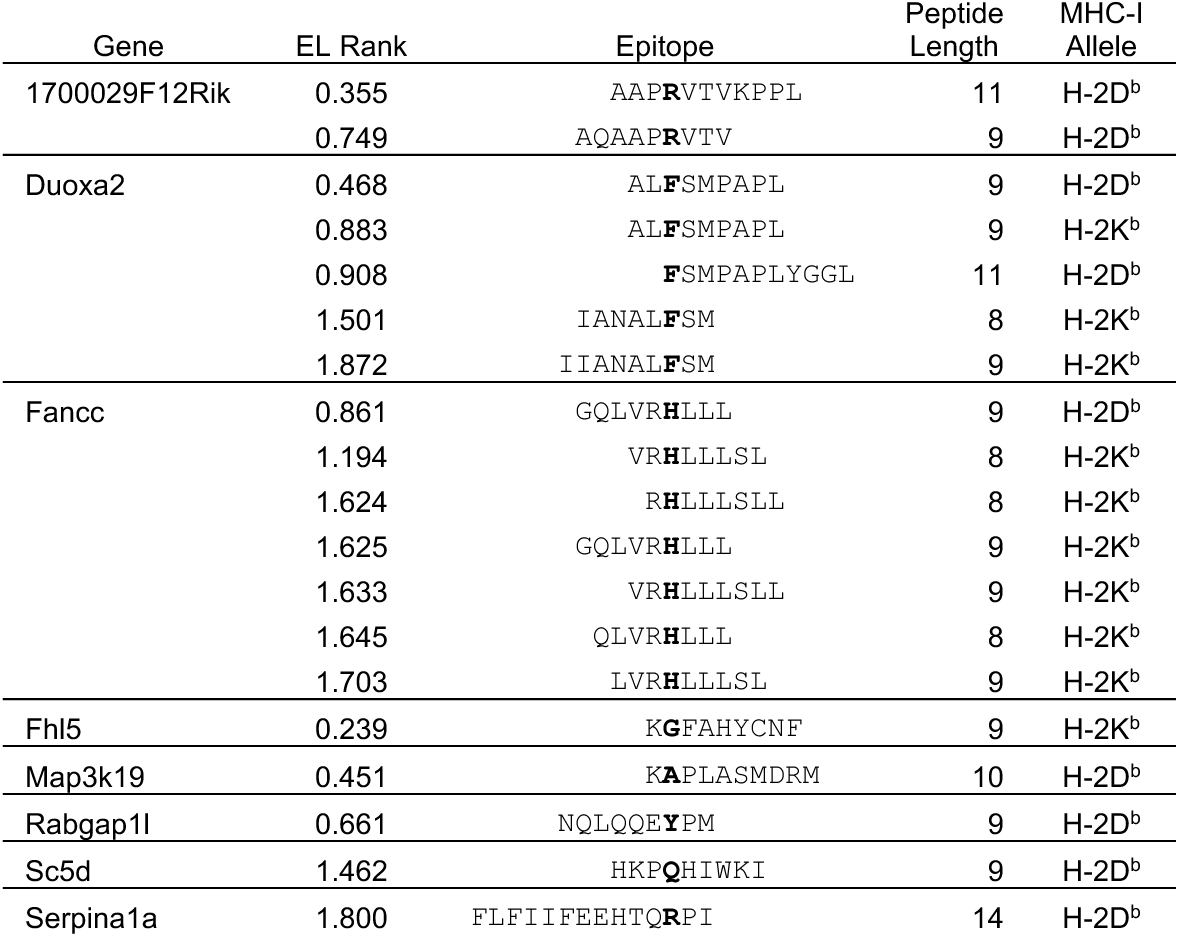

**Supplementary Data Table 1C.**
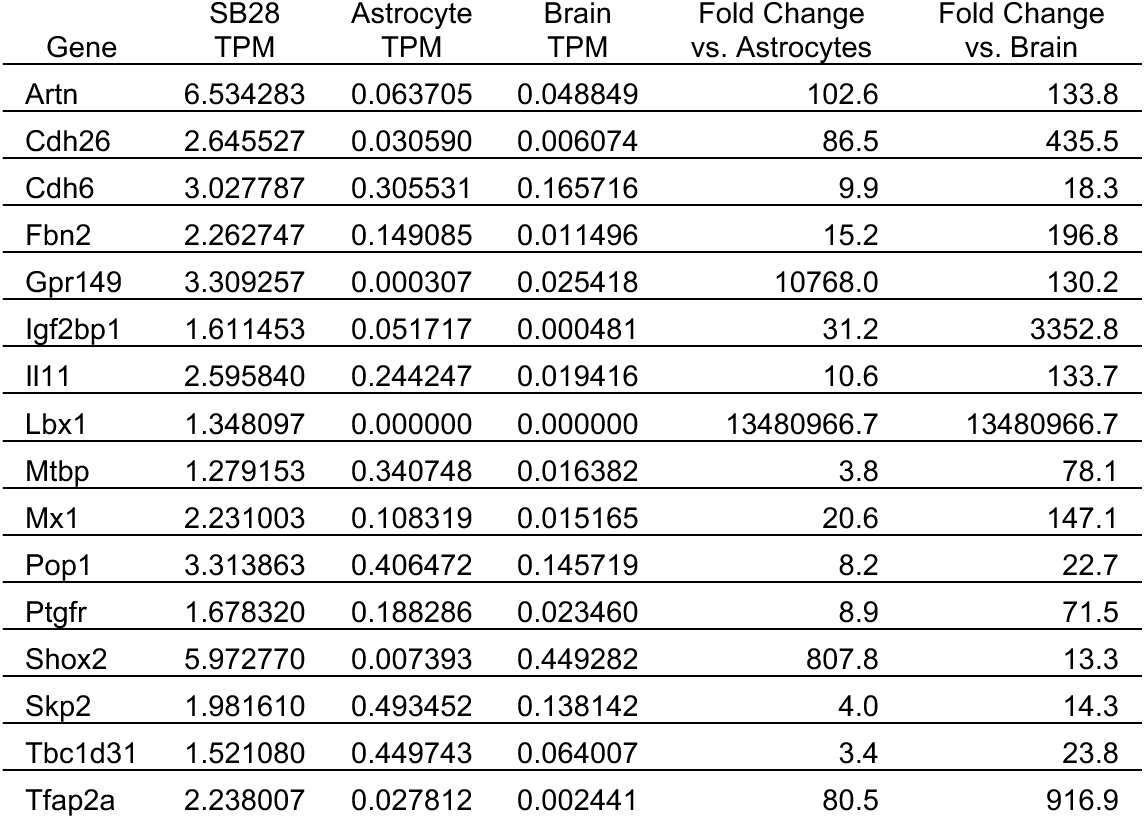

**Supplementary Data Table 1D.**
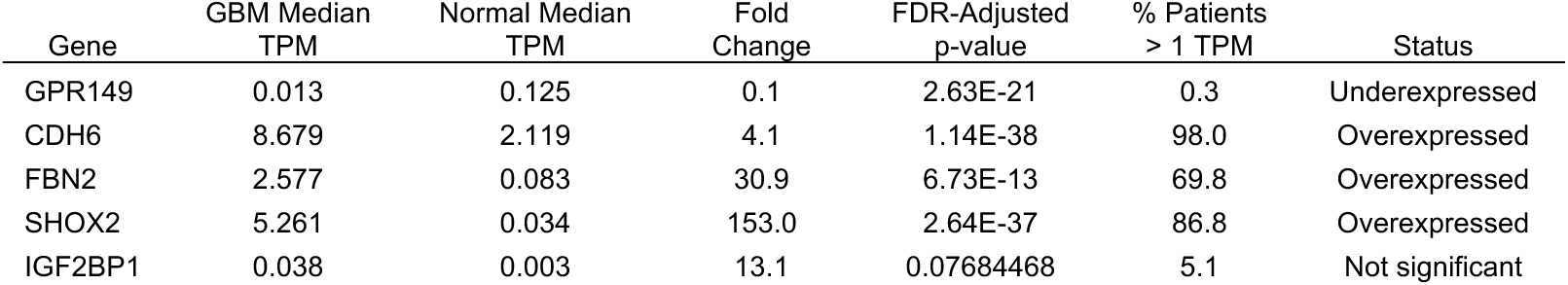

**Supplementary Data Table 1E.**
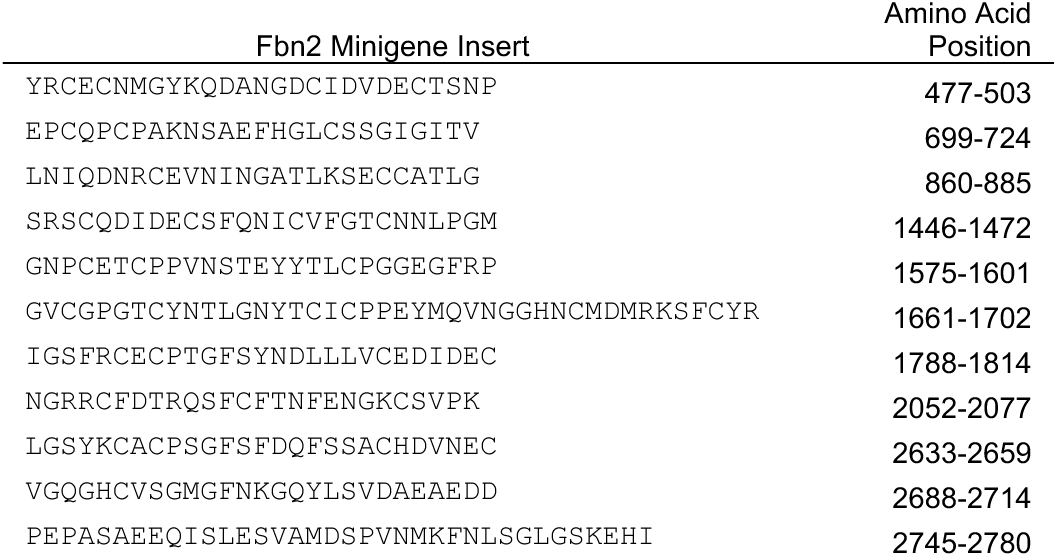

**Supplementary Data Table 2.**
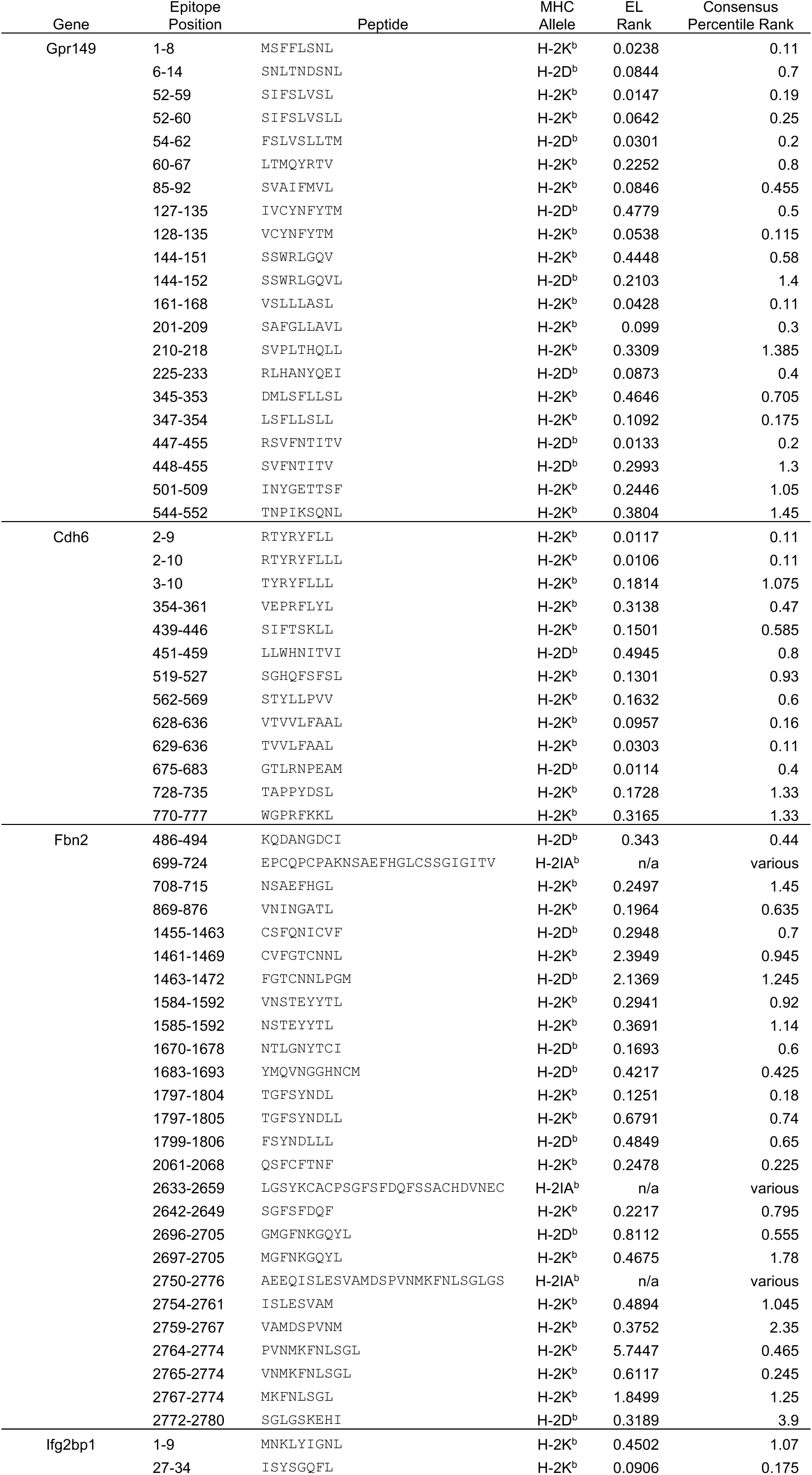
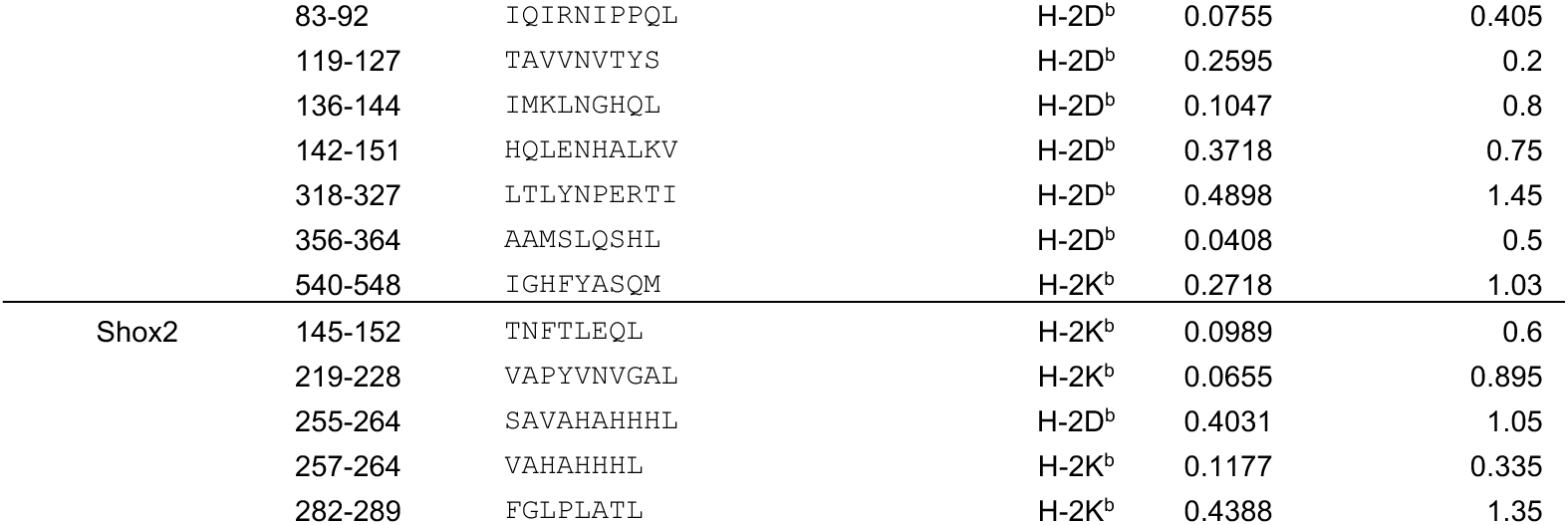

**Supplementary Data Table 3A.**
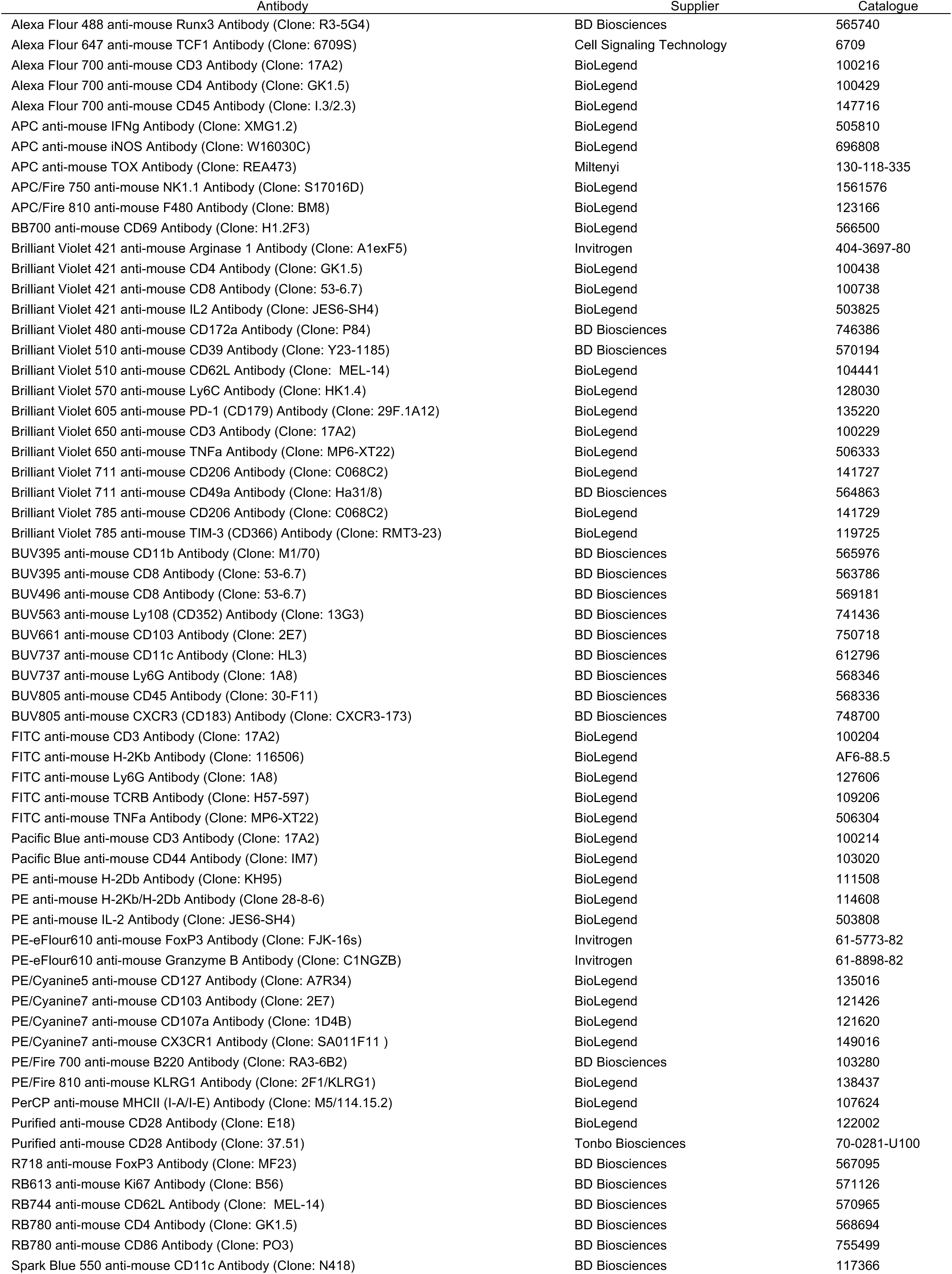
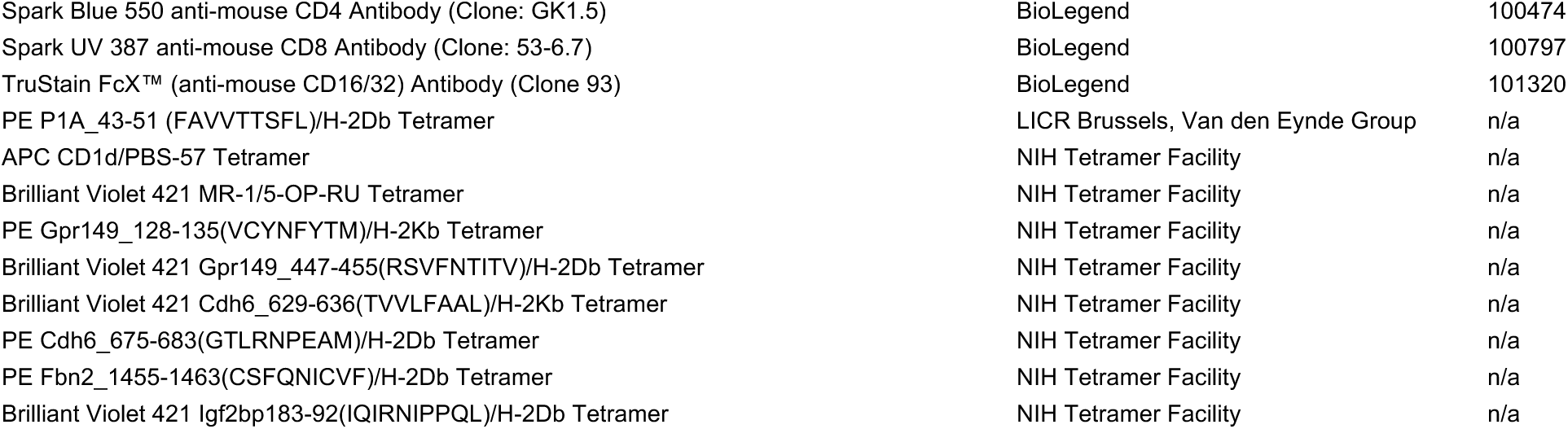

**Supplementary Data Table 3B.**
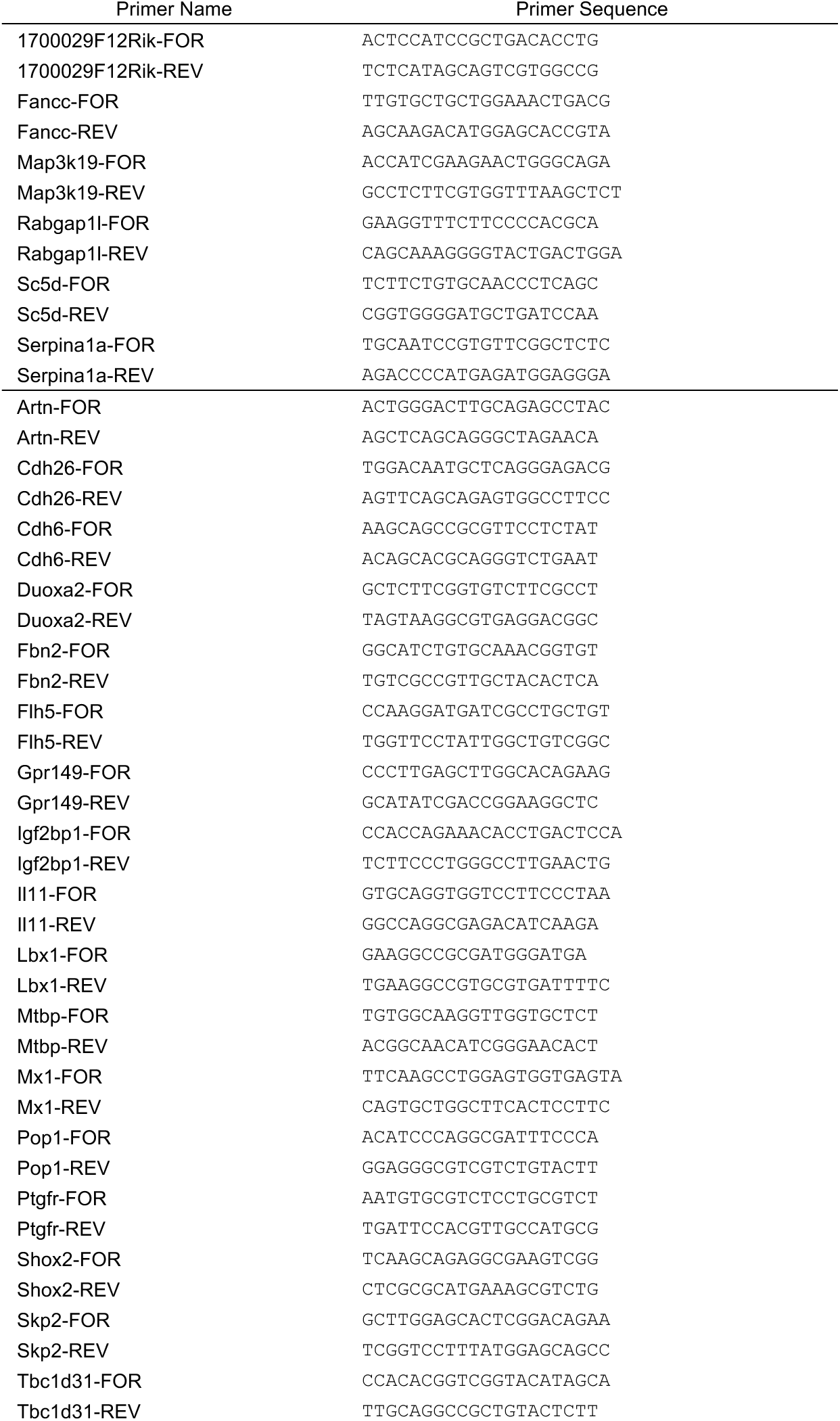
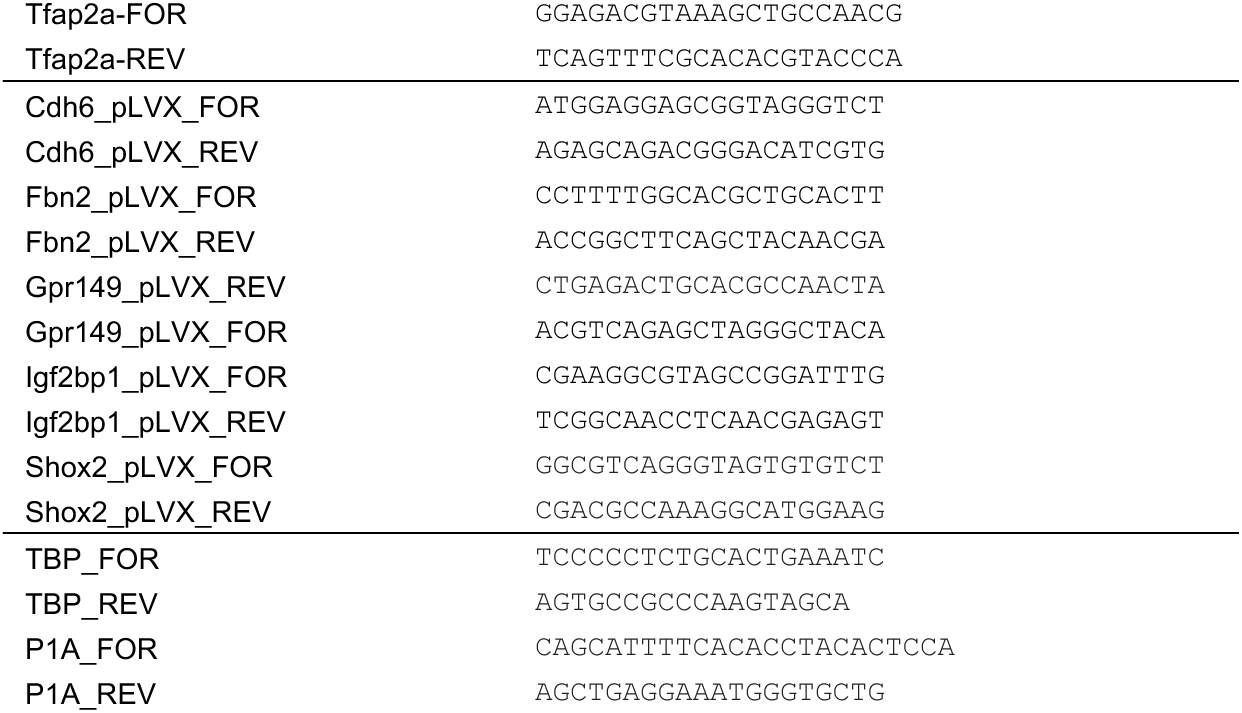

**Supplementary Table 1.**

**Tables 1A-B. Prediction scores for SB28 neoantigen candidates**

(A) Table showing 8-14mer minimal epitopes predicted by the IEDB Consensus algorithm to bind H-2D^b^ or H-2K^b^ with Consensus Percentile Rank ≤ 2.0.

(B) Table showing 8-14mer minimal epitopes predicted by the NetMHC4.1 algorithm to bind H-2D^b^ or H-2K^b^ with EL Rank ≤ 2.0.

**Table 1C. Expression of top 16 SB28 TAA candidates by RNASeq**

RNA from SB28 cells, astrocytes, or C57BL/6 brain was sequenced in triplicate. Expression level for each gene is shown by mean TPM (transcripts per million). Fold change was calculated as mean SB28 TPM / mean astrocyte or brain TPM.

**Table 1D. Expression of top 5 SB28 TAA candidates in human glioblastoma**

Expression of candidate genes in human IDH mutant glioblastoma (*n =* 295) compared to healthy cortical tissue (*n* = 270), based on TCGA and GTEx datasets, respectively.

**Table 1E. Minigene inserts included in the Fbn2 vaccine constructs**

Amino acid sequence of Fbn2 minigenes and position of minigene within whole Fbn2 gene is shown.

**Supplementary Table 2.**

**Table 2. Peptides used for immunogenicity testing of TAAs**

Amino acid position within original gene, peptide amino acid sequence, predicted MHC binding allele, and predicted MHC binding ranks are shown (EL Rank calculated by NetMHC4.1 Algorithm; Consensus Percentile Rank calculated by IEDB Consensus Algorithm). For Gpr149, Cdh6, Shox2, and Igf2bp1, all minimal epitopes with both an EL Rank < 0.5 and Consensus Percentile Rank < 1.5 were tested; for Fbn2, all predicted minimal epitopes within the minigene sequences with an EL Rank < 0.5 or Consensus Percentile Rank < 1.5 were tested. The top 48 H-2IA^b^ epitopes predicted by the IEDB MHC-II 2.22 Consensus Algorithm within the Fbn2 minigenes were all contained within three minigenes. Long 26- or 27-mer peptides corresponding to these three Fbn2 minigenes were also tested.

**Supplementary Table 3.**

**Table 3A. Antibodies used for flow cytometry.**

**Table 3B. Primers used for RT-qPCR.**

